# Fission yeast RPA–TERT–Tpz1^TPP1^ complex promotes telomere extension and suppresses telomere recombination

**DOI:** 10.64898/2026.04.26.720725

**Authors:** Bettina A. Moser, Madeline Points, Sourav Agrawal, Adam C. Didier, Amanda K. Mennie, Ci Ji Lim, Yong-jie Xu, Toru M. Nakamura

## Abstract

Telomerase maintains chromosome ends by extending telomeric DNA, yet how recruited telomerase becomes productively engaged remains poorly understood. Recent studies found that Replication Protein A (RPA) contributes to telomerase stimulation through interaction with TERT in humans and with the TPP1 ortholog Est3 in budding yeast, suggesting a direct role in telomerase activation. Here, we provide genetic and structural modeling evidence for a RPA-Trt1^TERT^-Tpz1^TPP1^ ternary complex that promotes telomere extension while suppressing recombination in fission yeast. Guided by results from genetic screen, followed by AlphaFold3 modeling and systematic mutagenesis of RPA, Trt1, and Tpz1, we identify four key interfaces supporting telomerase function: Ssb1^RPA1^-Trt1, Ssb2^RPA2^-Trt1, Ssb2^RPA2^-Tpz1, and the TEL-patch-mediated Trt1-Tpz1 interaction. Notably, Tpz1-R81, previously assigned as the TEL patch, instead contacts Ssb2 in the complex. Epistasis and suppressor analyses indicate that the newly identified RPA-Trt1 and RPA-Tpz1 interfaces collaborate with the Trt1-Tpz1 interface to allow telomerase activation after recruitment. Furthermore, comparative analyses using AlphaFold3 suggest that these interactions are likely conserved in budding yeast and humans. Collectively, these findings support a model in which RPA serves as an essential component of the active telomerase complex, coordinating TERT and TPP1-like factors to enable productive telomerase engagement.

**Author Summary:** Telomeres are specialized dynamic protective structures at the ends of eukaryotic chromosomes that must be properly maintained to preserve genome stability. Telomerase extends telomeric DNA, but how recruited telomerase becomes fully activated to promote telomere extension remains poorly understood. In this study, we use fission yeast to investigate the role of the conserved single-stranded DNA-binding protein complex Replication Protein A (RPA) in this process. We find that RPA forms a functional complex with the telomerase catalytic subunit TERT and Tpz1, a component of the telomere protection complex shelterin and the fission yeast ortholog of human TPP1. Genetic and structural analyses identify multiple interactions within the RPA-TERT-Tpz1 complex that are required for efficient telomere extension. Disrupting these interactions allows telomerase recruitment but prevents productive telomerase action at chromosome ends. Our results further suggest that similar mechanisms may operate in other organisms, including budding yeast and humans. These findings provide insight into how telomerase activity is regulated at chromosome ends.

## Introduction

Telomeres, the ends of linear chromosomes, consist of G-rich repetitive double-stranded DNA (dsDNA) terminating in a 3’ single-stranded overhang (G-tail). Telomerase, composed of the catalytic subunit TERT, an associated RNA template, and multiple additional regulatory and structural subunits, extends this overhang by synthesizing telomeric repeats to overcome the end-replication problem [1, 2].

In fission yeast *Schizosaccharomyces pombe*, telomere protection and telomerase regulation are mediated by a conserved shelterin complex consisting of the dsDNA-binding protein Taz1 (TRF1/TRF2 homolog), the G-overhang-binding protein Pot1, and the bridging components Rap1, Poz1 (TIN2 homolog), Tpz1 (TPP1 ortholog), and Ccq1 [3–5]. Telomerase recruitment to chromosome ends depends on Ccq1, whose phosphorylation at T93 by the checkpoint kinases Rad3^ATR^ and Tel1^ATM^ promotes interaction with the telomerase subunit Est1 [6, 7].

Tpz1^TPP1^ plays dual roles in telomerase regulation. First, conserved TEL-patch residues promote telomerase activity. For example, the *tpz1-K75A* mutant maintains only very short telomeres despite increased Trt1 recruitment, suggesting a defect in telomerase activation after recruitment [8]. Another proposed TEL-patch mutant, *tpz1-R81E*, completely loses telomeres and has been reported to impair telomerase recruitment [9]. Secondly, Tpz1 is SUMOylated at K242, a modification that negatively regulates telomerase recruitment [10, 11]. Together with a conserved SWSSS motif, the K242 SUMOylation promotes interaction with the Stn1–Ten1–Polα complex to facilitate C-strand synthesis, thereby limiting Replication Protein A (RPA) binding, Rad3^ATR^ signaling, and telomerase association at telomeres [12].

Telomere DNA replication is tightly coordinated with the regulation of the shelterin and Stn1–Ten1–Polα complexes. As replication forks progress through telomeric DNA, exposed ssDNA is first bound by RPA [13, 14]. RPA is an abundant heterotrimeric ssDNA-binding complex consisting of RPA1/RPA70, RPA2/RPA32, and RPA3/RPA14 that resolves secondary DNA structures during DNA replication and repair. It interacts directly with multiple DNA damage response factors, including the checkpoint kinase ATR [15, 16]. At telomeres, RPA is subsequently replaced by POT1, and this transition is essential for suppressing checkpoint-mediated cell-cycle arrest and promoting CST (CTC1–STN1–TEN1)–Polα function [13, 17–19]. Because RPA transiently occupies the telomeric ssDNA prior to POT1 loading, it is uniquely positioned to coordinate ATR-dependent signaling and telomerase action at chromosome ends.

RPA binds ssDNA through four oligosaccharide-binding (OB)-fold domains (DBD-A, -B, and -C of RPA1 and DBD-D of RPA2) [15, 20]. Mutations in Rfa1^RPA1^ and Rfa2^RPA2^ cause substantial telomere shortening and replicative senescence in budding yeast *Saccharomyces cerevisiae* [21, 22], while mutations in human RPA1 and RPA2 are associated with telomere shortening and premature aging [23–25]. Recent studies have begun to explain these observations: human RPA stimulates telomerase processivity through a direct RPA2–TERT interaction [26], and budding yeast RPA2 interacts with the TPP1 ortholog Est3 to regulate telomerase activity [22]. These findings parallel the role of the telomerase-specific RPA-like TEB (Teb1–Teb2–Teb3) complex in *Tetrahymena* [27, 28]. However, it remained unclear whether canonical RPA functions equivalently to the telomerase-specific TEB complex in *Tetrahymena* by simultaneously forming a functional complex with both TERT and TPP1 orthologs.

In fission yeast, only a single mutation in RPA1, *ssb1-D223Y*, has been reported to cause telomere shortening [29]. However, this mutation substantially reduces Ssb1 protein levels [30], making it difficult to distinguish telomere-specific defects from broader consequences of limiting the essential ssDNA-binding factor Ssb1. Here, we identify multiple mutations in Ssb1^RPA1^ and Ssb2^RPA2^ that cause severe telomere shortening or complete telomere loss without substantially reducing protein levels. Guided by AlphaFold3 structural predictions [31], we provide genetic evidence for a functional RPA–Trt1–Tpz1 complex at fission yeast telomeres. Our analyses indicate that the predicted Ssb1^RPA1^–Trt1^TERT^, Ssb2^RPA2^–Trt1^TERT^, Ssb2^RPA2^–Tpz1^TPP1^, and TEL-patch-mediated Trt1^TERT^–Tpz1^TPP1^ interfaces make partially redundant yet essential contributions to telomerase activation and telomere extension while suppressing telomere recombination. We further provide evidence for functional conservation of the Rfa1^RPA1^–Est2^TERT^ interface in budding yeast and the RPA1-TERT interface in humans, and reinterpret earlier mutational studies in the context of conserved RPA–TERT–TPP1-like regulatory complexes in both budding yeast and humans. Collectively, these findings identify the RPA–Trt1–Tpz1 ternary complex as a functional module that promotes telomerase activation by facilitating access of the telomeric 3’ end to the TERT catalytic center while suppressing telomere recombination.

## Results

### Ssb1^RPA1^ mutations reveal residues critical for telomere maintenance

We previously reported 25 mutants of Ssb1, the largest subunit of RPA in fission yeast, that are sensitive to the genotoxins hydroxyurea (HU) and/or methyl methane sulfonate (MMS) [30]. To determine whether any of these mutations also affect telomere maintenance, we examined telomere length by Southern blot analysis (Fig S1A). This analysis identified eight mutants (*ssb1-1*, *2*, *7-11*, and *13*) with very short telomeres (Fig S1B-C). Two of these mutants, *ssb1-2* (*D223Y*) and *ssb1-7* (*R11C-C69S-D223N*), exhibited substantially reduced Ssb1 protein expression, similar to the budding yeast equivalent *rfa1-D228Y* [32]. Nevertheless, prior work indicates that mutations at D223 itself are likely causative for telomere shortening [29, 33, 34]. It should also be noted that there are other mutant alleles (*ssb1-4*, *18*, and *23*) that caused substantial reduction in Ssb1 protein levels but showed only moderate to no reduction in telomere length (Fig S1C).

Among all mutants analyzed, *ssb1-12* (*W523C*) showed the most severe telomere phenotype. Despite near wild-type (wt) Ssb1 protein expression and robust cell growth [30], this mutant completely lost telomeres (Fig S1B), suggesting the formation of circular chromosome survivors [35]. To confirm this phenotype, we integrated *W523C*, *W523A,* or *W523S* at the endogenous *ssb1* locus (Figs 1A-B and S2A) and immediately introduced a *ssb1*^+^ plasmid to prevent telomere loss during strain construction. After subsequent loss of the plasmid, all three alleles showed similar early telomere shortening followed by complete telomere loss after further outgrowth (Fig 1B). These results establish that W523 is essential for telomere maintenance in fission yeast. On the other hand, since only circular chromosome survivors, but not earlier-generation cells with linear chromosomes, showed severe sensitivity to MMS, HU, or Camptothecin (CPT), *ssb1-W523C* does not appear to have a major impact on general DNA damage repair or checkpoint functions of the RPA complex (Fig S4A).

**Fig 1.**
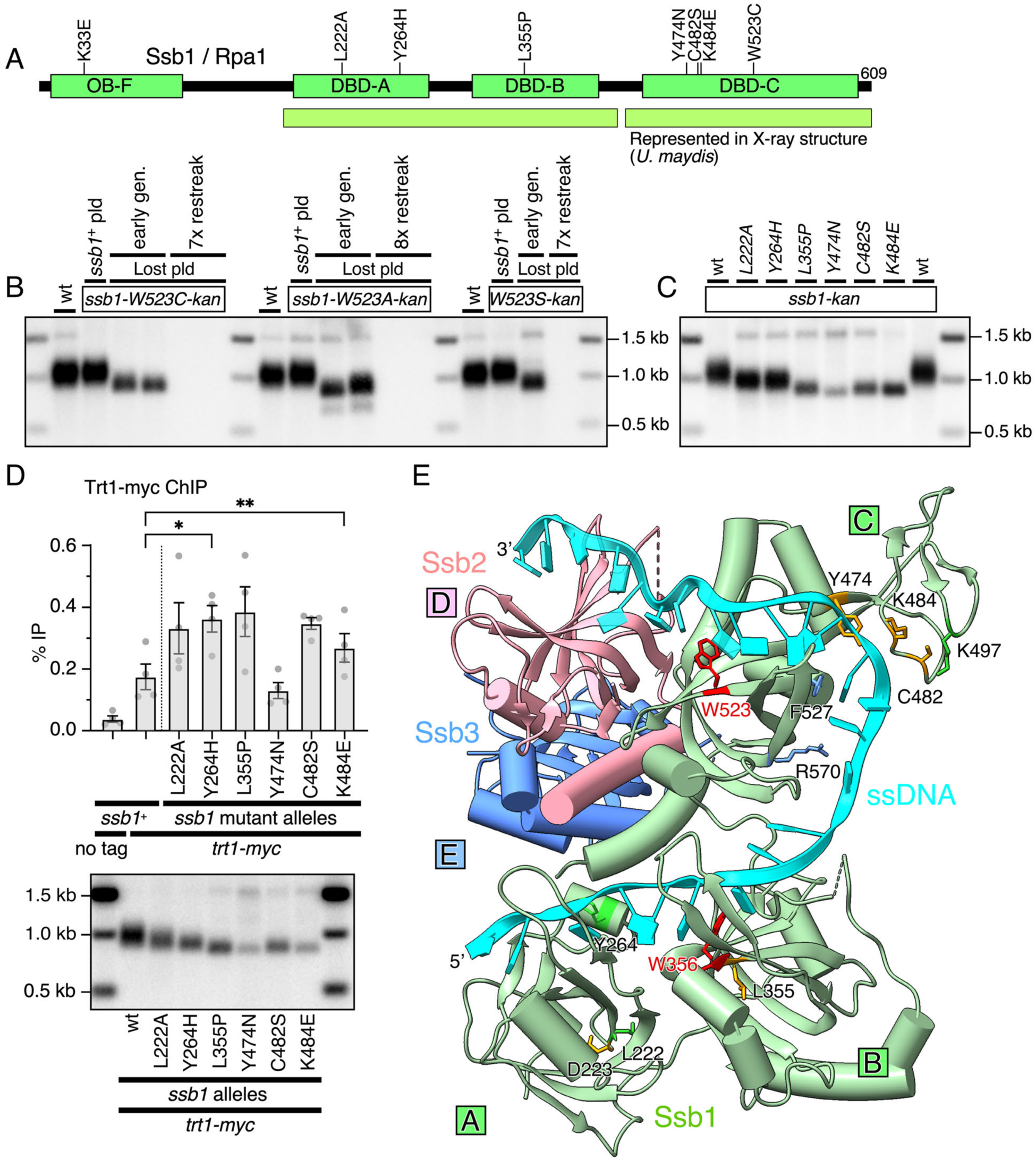
Ssb1 mutants with severe telomere maintenance defects. **(A)** Schematic representation of fission yeast Ssb1/Rpa1 showing the N-terminal OB-Fold domain (OB-F), DNA-Binding-Domains A-C (DBD-A, -B, -C), and the location of mutated residues for integrated mutants. The region represented in the X-ray structure from *U. maydis* RPA, shown in (E), is indicated below. **(B)** Telomere length analysis by Southern blot of *ssb1-W523* mutant strains in the presence or after loss of a *ssb1*^+^ plasmid. **(C)** Telomere length analysis of the indicated *ssb1* mutant strains. All strains were extensively restreaked to obtain terminal telomere phenotypes. Southern blot analysis for additional independently generated and restreaked *ssb1* mutant strains is shown in Fig S2B. **(D)** Trt1 binding to telomeres in the indicated *ssb1* mutant strains, analyzed by Trt1-myc ChIP (top), with corresponding telomere length analysis of the strains used for ChIP (bottom). Statistical significance was assessed by ANOVA with Dunnett’s comparison test (*P ≤ 0.05; **P ≤ 0.01). See S1 Data for individual % IP values and additional statistical analysis. **(E)** X-ray structure of *U. maydis* RPA bound to ssDNA [38], labeled using fission yeast nomenclature. Positions of Ssb1 residues analyzed in this study, as well as additional residues of interest, are indicated.

Because the *ssb1*^+^ plasmid allows *W523C* cells to maintain wt-length telomeres, we used this system to identify the causative single-site mutations within several multi-missense alleles. We focused on *ssb1-1*, *8*, *10*, *11*, and *13* because these alleles maintain near-wt Ssb1 protein expression [30] (Fig S1B). Plasmid-based complementation analysis revealed that *L355P*, *Y474N*, *C482S*, and *K484E* cause severe telomere shortening, while *Y264H* and *K497E* cause milder shortening (Fig S1D).

We also tested fission yeast equivalents of human RPA1 mutations associated with telomere shortening, namely *V227A*, *E240K*, and *T270A* [24]. In Ssb1, these correspond to *L222A*, *D235K*, and *T265A*, respectively (Fig S3A). In the plasmid assay, *L222A* caused modest telomere shortening, whereas *T265A*, despite being adjacent to Y264, had no detectable effect on telomere length (Fig S1E). *D235K* also did not cause telomere shortening (Fig S1E). Interestingly, budding yeast *rfa1-F269L* (Y264^Ssb1^/F269^RPA1^) and *rfa1-L227S* (L222^Ssb1^/V227^RPA1^) mutations (Fig S3A) hardly affected telomere length on their own but promoted HR-dependent survivor generation in telomerase RNA mutant *tlc1Δ* and suppressed temperature sensitivity of *cdc13-1* [36, 37].

Single-site mutations that caused short telomeres in the plasmid assay were then integrated at the endogenous *ssb1* locus (Fig S2A). Western blot analysis revealed near-wt expression of all mutant proteins except L355P, which showed a slight reduction like *ssb1-8* (*R330S-L355P*) (Fig S2C). Consistent with the plasmid assay, *L355P*, *Y474N*, *C482S*, and *K484E* caused very short telomeres, whereas *L222A* and *Y264H* caused moderate shortening (Fig 1C and S2B). Because *K33E* and *Y264H* each caused only mild telomere shortening, the stronger phenotype of *ssb1-1* (*K33E-Y264H*) indicates that these two mutations contribute additively to telomere shortening (Figs 1C and S2D). Unlike *W523C*, all integrated single-site mutants maintained stable telomeres even after extensive outgrowth.

We next asked whether the telomere shortening in these *ssb1* mutants reflects defective telomerase recruitment. Chromatin immunoprecipitation (ChIP) assays for Trt1 detected near-wt-level or increases in telomere binding for all mutants (Figs 1D and S2E). Thus, telomere shortening in these strains cannot be explained by reduced telomerase recruitment and instead might reflect impaired ability of telomerase to extend telomeres.

Because the fission yeast Tpz1 TEL-patch mutant *K75A* is also defective in telomerase activation [8], we asked whether the identified Ssb1 residues act redundantly with the Tpz1 TEL-patch. When combined with *tpz1-K75A*, milder *ssb1* mutants (*L222A* and *Y264H*) maintained short telomeres comparable to *tpz1-K75A* alone (Fig S2F). By contrast, stronger telomere-shortening mutants (*L355P*, *Y474N*, *C482S*, and *K484E*) showed severe telomere dysfunction in the *tpz1-K75A* background. Multiple double mutant clones either lost telomeres completely or displayed subtelomeric rearrangements and telomere amplification (Fig S2G-H). These results identify L355, Y474, C482, K484, and especially W523 as key Ssb1 residues for telomere maintenance.

To place these residues in a structural context, we mapped them onto the X-ray crystal structure of *Ustilago maydis* RPA bound to ssDNA [38] (Figs 1E and S3A). In the DBD-C domain (Fig 1A), W523 contacts a DNA base directly, whereas Y474, C482, and K484 cluster around the ssDNA. Nearby K497, which causes mild telomere shortening, lies in the same structural region. In the DBD-B domain, L355 lies adjacent to W356, which also contacts a DNA base, suggesting that *L355P* may disrupt the ssDNA-binding function of the neighboring W356. Consistent with this idea, recent work in budding yeast showed that *rfa1-W360A* (W356 in Ssb1) causes progressive telomere loss and replicative senescence [22]. On the other hand, budding yeast *rfa1-Y483L* (Y474 in Ssb1) was found to maintain extremely short, yet stable, telomeres [22]. Among the DBD-A mutants, Y264 likewise contacts a DNA base. Although not yet tested in fission yeast, the DBD-C residues F527 and R570, corresponding to the budding yeast *rfa1-F537A* and *R580E*, which cause very short telomeres [22], are also positioned adjacent to ssDNA. These observations indicate that Ssb1/RPA1 residues proximal to ssDNA are among the most important determinants of RPA function at telomeres.

### Ssb1-W523C uncouples telomerase recruitment from telomerase activation

We next asked why *ssb1-W523C* cells lose telomeres. In fission yeast, the Rad3^ATR^–Rad26^ATRIP^ kinase complex promotes telomerase recruitment by stimulating the Ccq1–Est1 interaction [6]. Because RPA-coated ssDNA recruits ATR–ATRIP and promotes phosphorylation of downstream checkpoint and DNA repair targets [39–41], we tested whether the *W523C* mutation affects recruitment of RPA, Rad3–Rad26, or telomerase to telomeres. Since telomeres in *W523C* cells show progressive shortening, we constructed and used a series of isogenic strains spanning longer-than-wt to progressively shorter telomeres (see Materials and Methods and Fig S5) and monitored telomere-length-dependent changes in protein association by ChIP.

Both RPA (Ssb2 and Ssb3) and the ATR complex (Rad26) exhibited robust binding to telomeres in *ssb1-W523C* cells, and this binding increased significantly as telomeres shortened (Fig 2A-C). Consistently, *ssb1-W523C* cells exhibit a higher fraction of cells with intense Ssb3-GFP foci upon telomere shortening, while the frequency of Ssb3-GFP foci formation in the isogenic strain with longer telomeres was comparable to *ssb1^+^* cells, indicating that the W523C mutation does not likely cause a general defect in DNA replication or DNA damage responses (Fig S4B-C).

**Fig 2.**
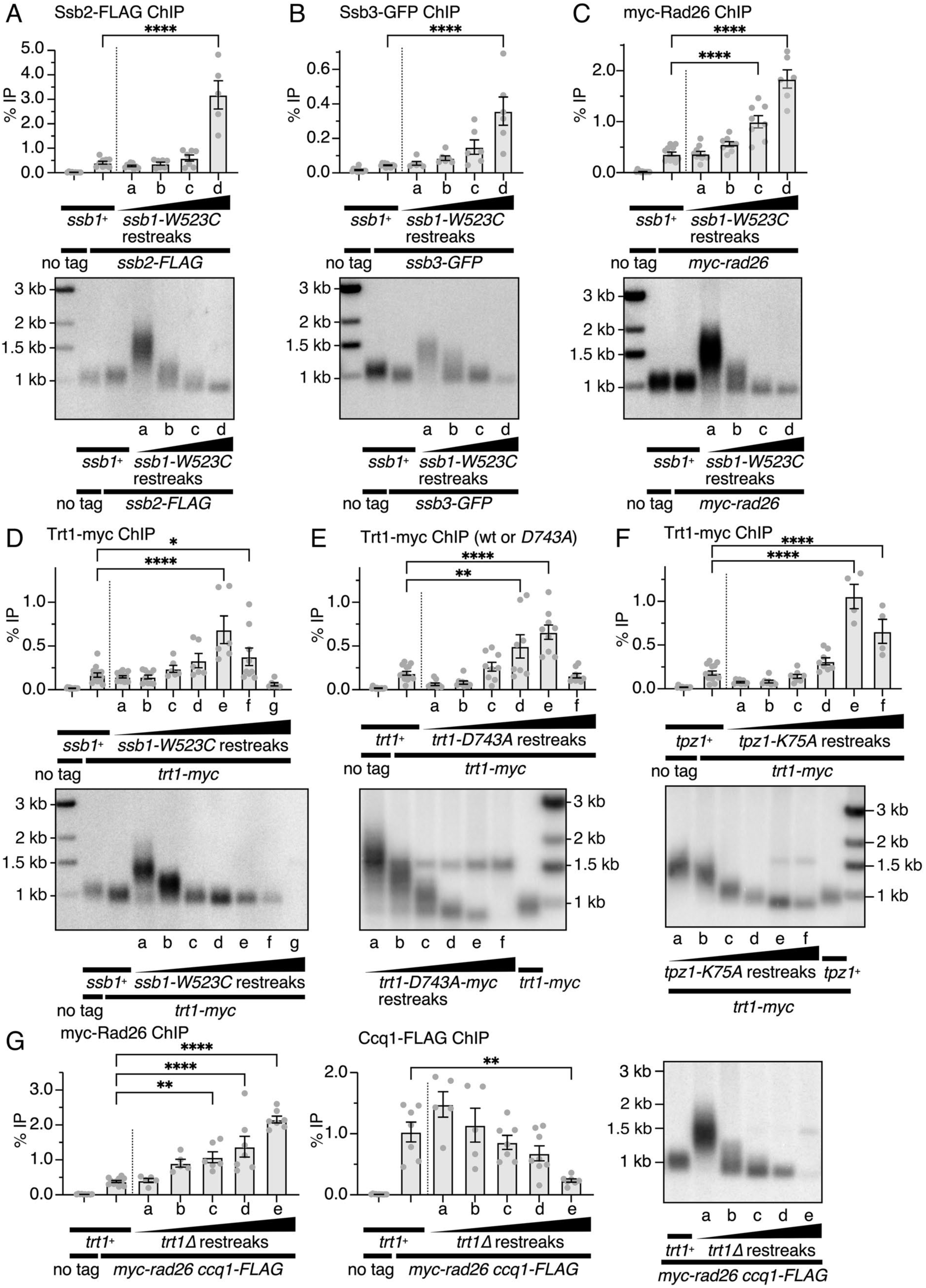
*ssb1-W523C* cells exhibit a telomerase activation defect. **(A-G)** Binding of the indicated proteins to telomeres in isogenic *ssb1-W523C* (A-D), *trt1-D743A* (E), *tpz1-K75A* (F), and *trt1Δ* (G) strains carrying different telomere lengths, analyzed by ChIP (top or left), with corresponding telomere length analysis of the strains used for ChIP (bottom or right). Statistical significance was assessed by ANOVA with Dunnett’s comparison test (*P ≤ 0.05; **P ≤ 0.01; ****P ≤ 0.0001). See S1 Data for individual % IP values and additional statistical analysis.

Trt1 recruitment also initially increased as telomeres shortened, paralleling the accumulation of RPA and Rad26^ATRIP^ (Fig 2D). However, once telomeres became extremely short, enhanced Trt1 binding was lost. A similar pattern of initial increase, followed by reduction, of telomere binding of Trt1 was observed in the catalytically inactive *trt1-D743A* mutant (Fig 2E), indicating that *ssb1-W523C* cells are not defective in telomerase recruitment, but instead fail to engage in productive elongation of telomeres. Notably, the defect in *ssb1-W523C* is more severe than in *tpz1-K75A* cells, because *K75A* cells showed substantial increases in Trt1 binding as telomeres shortened, yet stably maintained telomeres and enhanced association of Trt1 with telomeres (Fig 2F).

Previous work showed that Rad3^ATR^/Tel1^ATM^ kinase-dependent Ccq1-T93 phosphorylation initially increases as telomeres shorten in *trt1Δ* cells and subsequently disappears once telomeres become very short [42]. By contrast, phosphorylation of the checkpoint kinase Chk1 is initially suppressed but becomes strongly induced in later generations of *trt1Δ* cells [42]. ChIP assays explain this transition: when telomeres become very short, Rad26^ATRIP^ binding remains robust, whereas Ccq1 binding drops dramatically (Fig 2G). Thus, loss of Ccq1 accumulation and Ccq1-T93 phosphorylation likely explain why Trt1 binding is lost in *ssb1-W523C* and *trt1-D743A* cells when telomeres become critically short. Moreover, because disruption of the Tpz1–Ccq1 interaction activates checkpoint signaling by reducing Ccq1 accumulation at telomeres [43–45], loss of Ccq1 also explains why cells with very short telomeres eventually activate Chk1.

Taken together, these results identify *ssb1-W523C* as a separation-of-function mutant that uncouples telomerase recruitment from telomerase activation.

### AlphaFold3 modeling predicts a functional RPA–Trt1–Tpz1 ternary complex

The RPA2–TERT and Rfa2^RPA2^–Est3^TPP1^ interactions that promote telomerase activity in humans [26] and budding yeast [22] have not been examined in fission yeast. We therefore used AlphaFold3 [31] to test whether fission yeast RPA might form analogous interactions with Trt1^TERT^ and Tpz1^TPP1^.

When Trt1, the N-terminal OB-fold domain of Tpz1 (aa1-219), and all three RPA subunits were included in the modeling, the top-scoring AlphaFold3 models converged on a highly similar overall architecture (Fig S6A-B). Although full-length Ssb1^RPA1^ was modeled, we focused primarily on aa177-609 because the orientation of the N-terminal OB-F domain, connected by a long linker, varied substantially across models (Fig 3 and S6A-B). Likewise, although full-length Ssb2 was modeled, the N-terminal unstructured region (aa1-49) and the C-terminal winged-helix (WH) domain (aa182-279) were excluded from further analysis due to their variable orientations. Moreover, truncation analysis showed that the Ssb2 WH domain is dispensable for telomere maintenance in fission yeast (Fig S7A-B), unlike the human RPA2 WH domain, which affects telomere maintenance but not RPA’s role as telomerase processivity factor [23, 26]. Similarly, Ssb3 did not directly contact either Trt1 or Tpz1 in the AlphaFold3 models, in agreement with the observation that *ssb3Δ* cells maintain wt telomeres (Fig S7C).

**Fig 3.**
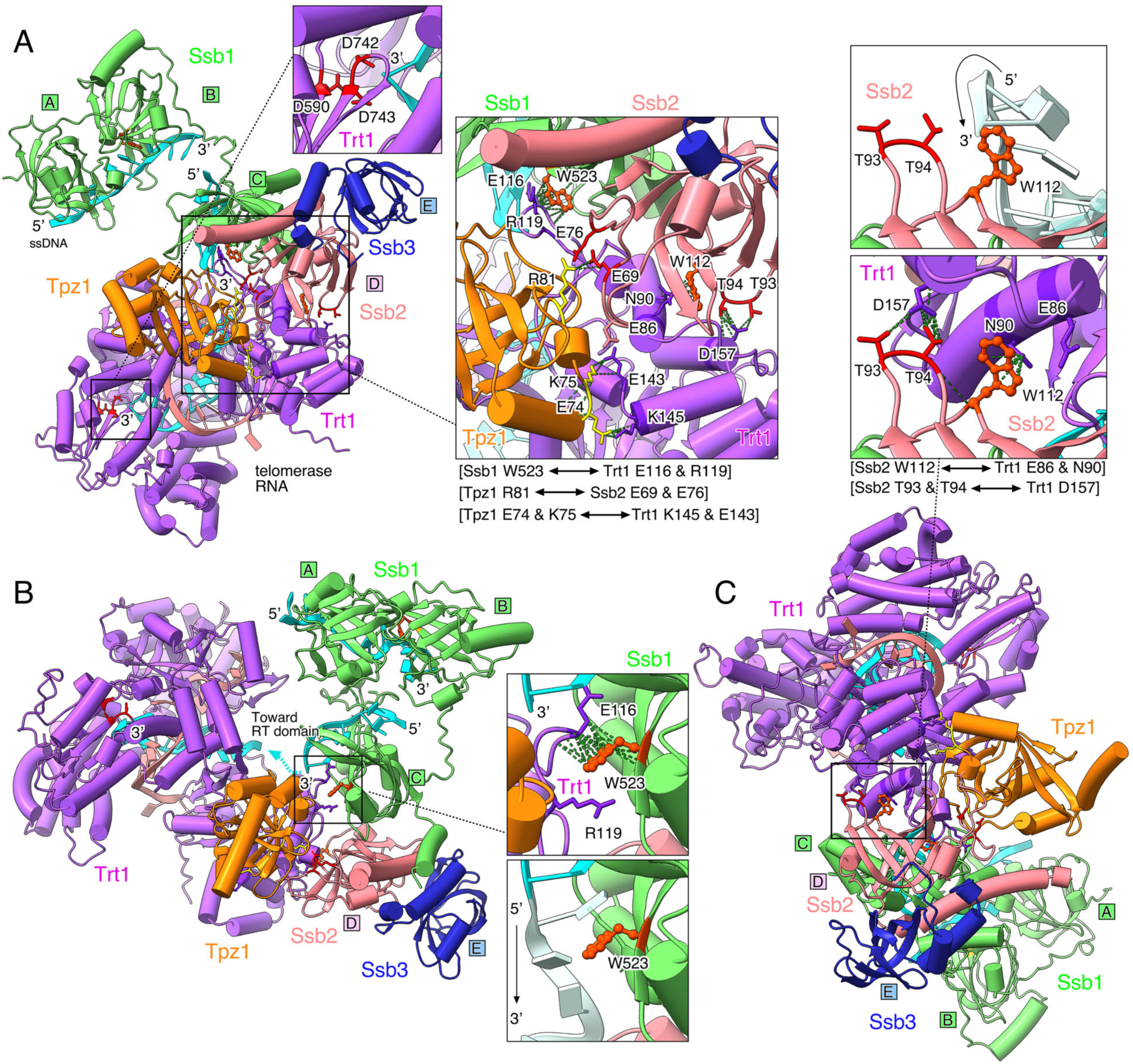
AlphaFold3 model of the RPA–Trt1–Tpz1 complex. **(A-C)** AlphaFold3 models containing Ssb1 (aa177-609), Ssb2 (aa50-181), full-length Ssb3, Tpz1 (aa1-219), and full-length Trt1, shown in different orientations. The models include the 3’ end of telomeric ssDNA (turquoise) and a short telomerase RNA template region (pink). DBD-A, -B, and -C of Ssb1, DBD-D of Ssb2, and OB-E of Ssb3 are indicated. (A) Overall model highlighting Trt1 catalytic site aspartates D590, D742, and D743 positioned adjacent to the 3’ end of telomeric ssDNA (small inset) and the predicted Ssb1–Trt1, Ssb2–Tpz1, and Trt1–Tpz1 interaction surfaces (large inset). (B) Detailed view of the predicted interaction of Ssb1-W523 with Trt1-E116/R119 (top inset) or with telomeric ssDNA (bottom inset). (C) Detailed view of the predicted interaction of Ssb2-W112 with telomeric ssDNA (top inset) or with Trt1-E86/N90 (bottom inset). The predicted interaction between Ssb2-T93/T94 and Trt1-D157 is also shown (bottom inset). The predicted interactions of Ssb1 and Ssb2 with Trt1 are structurally incompatible with their simultaneous engagement of ssDNA in the canonical RPA–ssDNA binding configuration.

The predicted fission yeast complex resembles the overall organization of the *Tetrahymena* telomerase complex, which includes TERT, the telomerase processivity factor complex TEB, and the TPP1-like factor p50 [28, 46–48] (Fig S6C). Because the AlphaFold3 model of RPA aligned well with the *U. maydis* RPA–ssDNA structure [38] (Fig S7D-F), we overlaid ssDNA from the crystal structure onto the RPA–Trt1–Tpz1 model (Fig 3). Based on the strong structural conservation of TERT, we also overlaid the RNA template region and telomeric ssDNA from the *Tetrahymena* telomerase structure, which positioned the 3’ end of telomeric ssDNA adjacent to catalytic aspartate residues D590, D742, and D743 of Trt1 (Fig 3A), required for 3’ nucleotide addition [49].

The model further predicted that Tpz1 TEL-patch residues E74 and K75 interact with Trt1 K145 and E143, respectively, thereby linking the TEL-patch to the TEN-domain of Trt1 (Figs 3A and S6D). Additional nearby interactions, including K75^Tpz1^–E116^Tpz1^ and S85^Trt1^–E143^Trt1^, may stabilize this interface (Fig S6D), and Trt1-R119 may also contact Tpz1-N122 (Fig S6E). These predictions are consistent with previous observations that *tpz1-K75A* and *tpz1*-*K75E* cause very short telomeres, whereas *tpz1-E74R* causes milder but detectable telomere shortening [8, 9].

In addition to the TEL-patch interface, AlphaFold3 predicted several RPA residues to interact directly with Trt1. These include Ssb1-W523, positioned near Trt1-E116 and R119 (Fig 3B), and Ssb2-W112, positioned adjacent to Trt1-E86 and N90 (Fig 3C). The predicted W112^Ssb2^–E86/N90^Trt1^ interaction is analogous to the W107^RPA2^–R83^TERT^ interaction shown to be critical for human RPA-mediated stimulation of telomerase processivity [26]. Importantly, both Ssb1-W523 and Ssb2-W112 also contact ssDNA in the canonical RPA–ssDNA structure (Fig S7D-E). Thus, if these residues engage Trt1 as predicted, they are unlikely to simultaneously occupy the canonical RPA–ssDNA-binding configuration (Fig 3B-C). Indeed, persistent binding of Ssb2 to ssDNA, as in the canonical RPA–ssDNA structure, would position the 3’ telomeric end away from the Trt1 reverse transcriptase (RT) domain (Fig 3B). This suggests a mechanism in which the formation of the RPA–Trt1–Tpz1 ternary complex releases the 3’ end from RPA and channels it toward the telomerase active site. The model also predicts a stabilizing interaction between Trt1-D157 and Ssb2-T93/T94, which lie close to the W112^Ssb2^–Trt1 interface but do not contact ssDNA (Figs 3C and S7D). Notably, this arrangement resembles the previously identified human D147^TERT^–T88^RPA2^ interaction [26].

The model also predicted that Tpz1-R81 interacts with Ssb2 E69 and/or E76 (Figs 3A and 4A). This was surprising because earlier work classified R81 as part of the Tpz1 TEL-patch, based on complete telomere loss and failure to recruit telomerase in *tpz1-R81E* cells [9]. However, our analysis found that *tpz1-R81E* cells still exhibit robust increases in Trt1 recruitment as telomeres shorten and lose Trt1 binding only when telomeres become extremely short (Fig 4B), like *ssb1-W523C* and *trt1-D743A* cells (Fig 2D-E). Thus, our data instead suggest that Tpz1-R81 and Ssb1-W523 likely contribute to a post-recruitment step of telomerase that is essential for telomerase activation.

**Fig 4.**
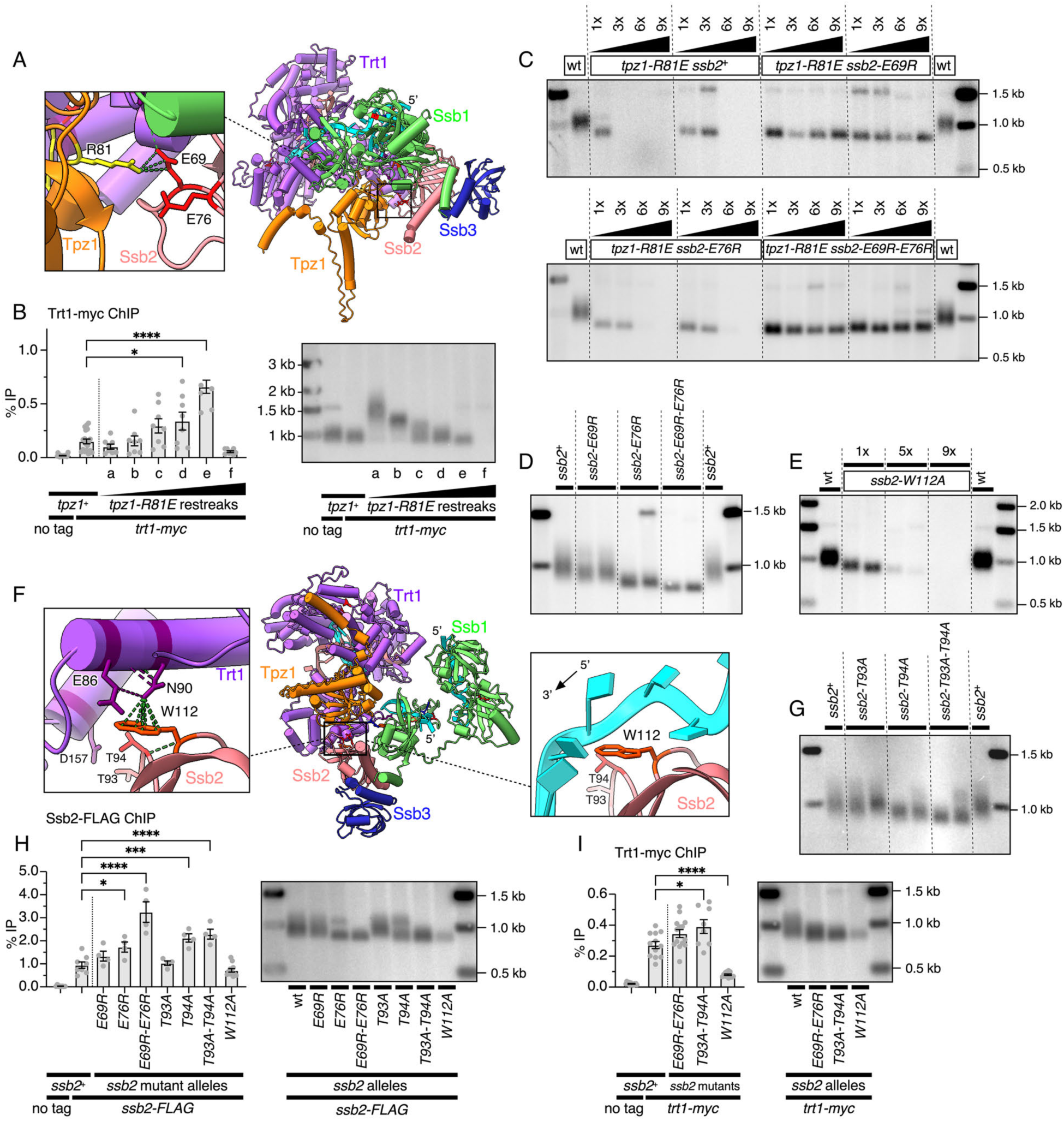
Characterization of *ssb2* mutants predicted to affect Ssb2–Tpz1 and Ssb2–Trt1 interactions. **(A)** AlphaFold3 model highlighting the predicted interaction of Tpz1-R81 with Ssb2-E69/E76. **(B)** Trt1-myc ChIP analysis of telomere binding in isogenic *tpz1-R81E* strains with different telomere lengths, with corresponding telomere length analysis. **(C)** Telomere length analysis by Southern blot of the indicated *ssb2* mutant strains in combination with *tpz1-R81E* after the indicated numbers of restreaks following sporulation of heterozygous diploid strains. Two independent clones are shown for each genotype. **(D)** Telomere length analysis by Southern blot of the indicated *ssb2* mutant strains after extensive restreaks to obtain terminal telomere phenotypes. **(E)** Telomere length analysis by Southern blot of *ssb2-W112A* mutant cells after indicated numbers of restreaks following sporulation of heterozygous diploid strains. **(F)** AlphaFold3 model highlighting the predicted interaction of Ssb2-W112 with Trt1-E86/N90 (left inset) or with telomeric ssDNA (right inset). **(G)** Telomere length analysis by Southern blot of the indicated *ssb2* mutant strains. All strains were extensively restreaked to obtain terminal telomere phenotypes. **(H, I)** ChIP analysis of telomere binding by Ssb2-FLAG (H) and Trt1-myc (I) in the indicated *ssb2* mutant strains, with corresponding telomere length analysis of the strains used for ChIP. For all ChIP assays, statistical significance was assessed by ANOVA with Dunnett’s comparison test (*P ≤ 0.05; ***P ≤ 0.001; ****P ≤ 0.0001). See S1 Data for individual % IP values and additional statistical analysis.

Taken together, the AlphaFold3 analysis provides a mechanistic framework for a RPA–Trt1–Tpz1 ternary complex in which RPA–TERT and RPA–Tpz1 interactions collaborate with the Tpz1 TEL-patch to promote engagement of the telomeric 3’ end with the Trt1 catalytic center.

### Ssb2^RPA2^-mediated interfaces with Tpz1 and Trt1 contribute to productive telomere maintenance

To test the predicted Ssb2–Tpz1 interface, we mutated Ssb2 residues E69 and E76, which are predicted to interact with Tpz1-R81. If the AlphaFold3 prediction is correct, charge-reversal mutations in Ssb2 might rescue the interaction defect caused by *tpz1-R81E*. Indeed, *ssb2-E69R* and *ssb2-E69R-E76R*, but not *ssb2-E76R*, rescued telomere loss in *tpz1-R81E* cells (Fig 4C). However, they did not restore wt telomere length, indicating that Tpz1-R81 remains important even when the charge interaction is partially compensated. This suppression required telomerase, because the corresponding *tpz1 ssb2* double mutants failed to maintain telomeres in a *trt1Δ* background (Fig S8A).

In a *tpz1*^+^ background, *ssb2-E69R* caused milder telomere shortening than *ssb2-E76R*, whereas *ssb2-E69R-E76R* produced the shortest telomeres (Fig 4D), indicating that E76 is more important than E69, although both contribute to telomere maintenance. Consistent with this interpretation, *E76R* and *E69R-E76R*, but not *E69R*, exhibited synergistic telomere loss when combined with *tpz1-K75A* (Fig S8B).

We next examined the predicted Ssb2–Trt1 interface (Figs 3C and 4F). We found that *ssb2*-*W112A* causes progressive telomere shortening followed by telomere loss (Fig 4E), consistent with the importance of the equivalent residue in the human system [26]. By contrast, *ssb2*-*T93A-T94A* caused telomere shortening but still allowed stable telomere maintenance (Fig 4G). T94 is found to be more important than T93, because *ssb2-T94A*, but not *ssb2-T93A*, caused substantial telomere shortening on its own and synthetic telomere loss with *tpz1-K75A* (Fig S8C).

To determine whether these Ssb2 mutants still bind telomeres efficiently, we performed ChIP assays. Cells carrying *E69R/E76R* or *T93A/T94A* mutations showed increased Ssb2 binding that inversely correlated with the severity of telomere shortening (Fig 4H), consistent with intact RPA binding to short telomeres. In contrast, *ssb2-W112A* did not show increased telomere binding to short telomeres despite robust Ssb2 protein expression (Figs 4H and S8D). Because W112 lies adjacent to ssDNA in the RPA–ssDNA complex (Figs 4F and S7D-E), this mutation might weaken not only telomeric binding but also RPA association with non-telomeric ssDNA, consistent with the poor growth of *ssb2-W112A* cells.

ChIP assays for Trt1 showed that both *ssb2-E69R-E76R* and *ssb2-T93A-T94A* cells recruit moderately increased levels of telomerase to short telomeres (Fig 4I), indicating that telomere shortening in these strains reflects reduced telomerase activity per recruited complex rather than defective recruitment. Indeed, even *tpz1-R81E ssb2-T93A-T94A* double mutants retained Trt1 binding (Fig S8E), indicating that robust telomerase recruitment can still occur even when both Ssb2–Tpz1 and Ssb2–Trt1 interfaces are simultaneously compromised. In contrast, *ssb2-W112A* cells showed a severe reduction in Trt1 binding in early generations before eventual telomere loss (Fig 4I), indicating that this mutation affects telomerase recruitment, consistent with an impairment of the RPA–Rad3^ATR^-dependent Ccq1–Est1 pathway due to poor RPA association at telomeres. On the other hand, because mutation of the equivalent residue in humans (W107^RPA2^) causes the most severe defect in RPA-dependent stimulation of TERT processivity [26], *ssb2-W112A* may also impair telomerase activation.

Taken together, these experiments provide strong support for the functional roles of the two predicted Ssb2–Tpz1 and Ssb2–Trt1 interfaces in productive telomere maintenance.

### Redundant RPA-telomerase-shelterin interactions ensure productive telomerase engagement

To complement the Ssb2 mutational analysis, we next mutated Trt1 residues predicted to reside at the Ssb1–Trt1 and Ssb2–Trt1 interfaces. Mutations *E86A*, *E86A-N90A*, *E116A*, *R119A*, *E116A-R119A*, and *D157A* in Trt1 all caused telomere shortening, whereas *N90A* maintained near wt telomere length (Figs 5A and S8F). Western blot analysis confirmed that these Trt1 mutations do not impair Trt1 protein expression (Fig S8G), and all mutants stably maintained their shortened telomeres.

**Fig 5.**
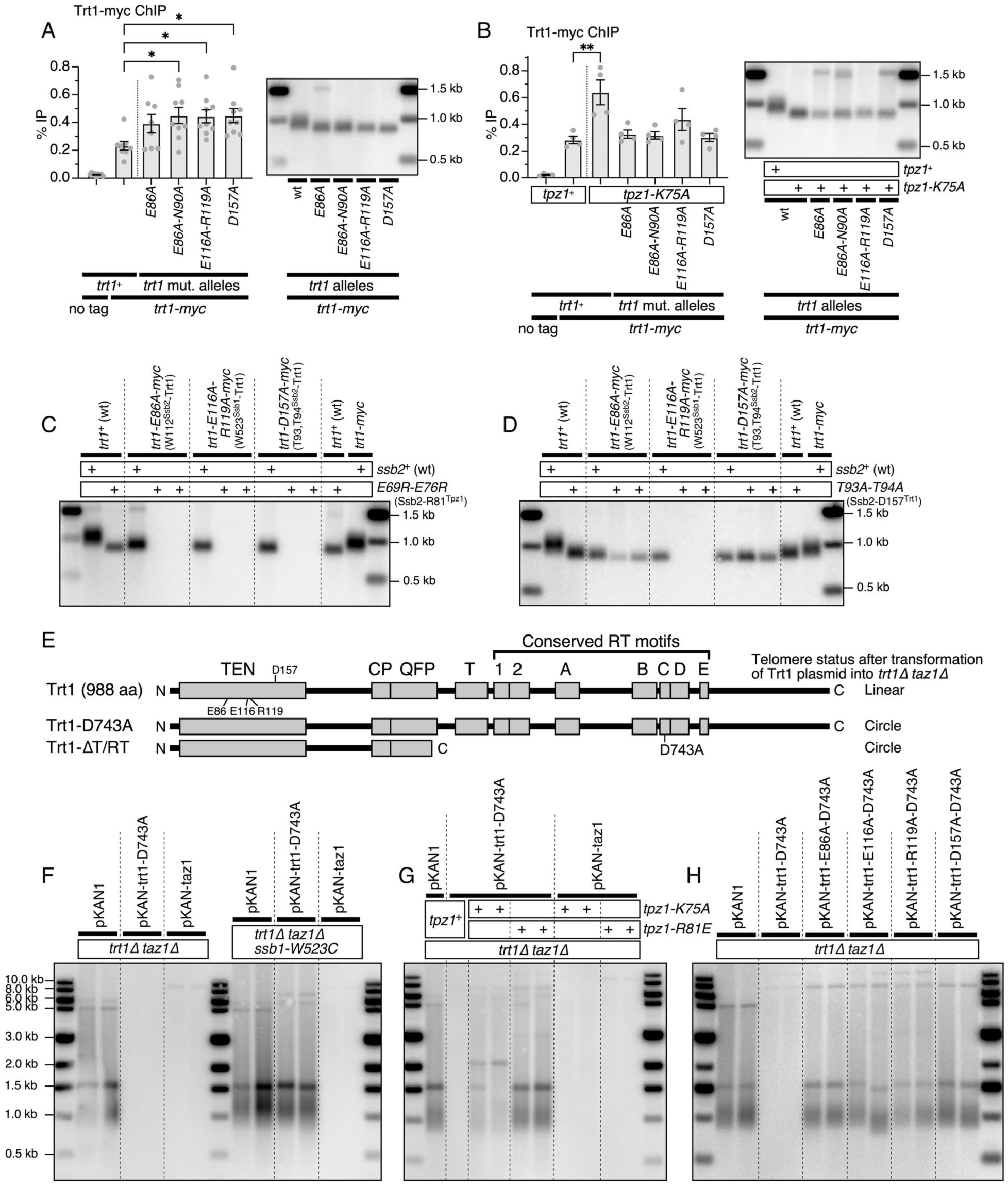
Ssb1–Trt1, Ssb2–Trt1, Trt1–Tpz1, and Ssb2–Tpz1 interactions redundantly regulate productive telomerase engagement with telomeric 3’ ends. **(A, B)** ChIP analysis of the indicated *trt1* mutants in *tpz1^+^* (A) or *tpz1-K75A* (B) background, with corresponding telomere length analysis by Southern blot of the strains used for ChIP. Statistical significance was assessed by ANOVA with Dunnett’s comparison test (*P ≤ 0.05; **P ≤ 0.01). See S1 Data for individual % IP values and additional statistical analysis. **(C, D)** Telomere length analysis by Southern blot of the indicated *trt1* mutants in *ssb2-E69R-E76R* (C) and *ssb2-T93A-T94A* (D) backgrounds. All strains were extensively restreaked to obtain terminal telomere phenotypes. **(E)** Schematic representation of fission yeast Trt1 constructs previously shown to induce chromosome circularization upon reintroduction into *trt1Δ taz1Δ* survivor cells [51, 52]. **(F-H)** Analysis of telomere loss in *trt1Δ taz1Δ* cells after reintroduction of plasmids expressing catalytically inactive *trt1-D743A* or *taz1*^+^ in either *ssb1*^+^ or *ssb1-W523C* backgrounds (F), *tpz1-K75A* or *tpz1-R81E* backgrounds (G), or various *trt1*-mutant plasmids in *trt1Δ taz1Δ* cells (H).

When combined with *tpz1-K75A*, however, nearly all Trt1 interface mutants, except for *N90A*, lost telomeres (Fig S8H), highlighting the importance of these residues for telomere maintenance. ChIP assays showed increased telomere occupancy for all tested Trt1 mutants (Fig 5A), consistent with the elevated telomerase association typically observed in short-telomere strains. Thus, these telomerase mutants reduce productive telomere extension per telomere-bound telomerase complex rather than telomerase recruitment.

To test how these Trt1 residues interact functionally with the Tpz1 TEL-patch, we measured binding of the same Trt1 mutants in a *tpz1-K75A* background before eventual telomere loss. We found that *trt1 tpz1-K75A* double mutants no longer showed increased Trt1 binding despite carrying short telomeres (Fig 5B), suggesting that the Ssb1/2-Trt1 interactions and the Tpz1 TEL-patch provide partially redundant contributions that together stabilize or enhance Trt1 association with short telomeres, while they are not required for telomerase recruitment per se.

We next tested genetic redundancy among the Ssb1–Trt1, Ssb2–Trt1, and Ssb2–Tpz1 interfaces. When *ssb2-E69R-E76R,* affecting the Ssb2–Tpz1 interface, was combined with *trt1-E86A* or *trt1-D157A* of the Ssb2–Trt1 interface or with *trt1-E116A-R119A* of the Ssb1–Trt1 interface, all double mutants exhibited synergistic telomere loss (Fig 5C). Likewise, combining *ssb2-T93A-T94A* at the Ssb2–Trt1 interface with *trt1-E116A-R119A* resulted in synergistic telomere loss (Fig 5D). By contrast, when *ssb2-T93A-T94A* was combined with *trt1-E86A* or *trt1-D157A*, which are all predicted to affect the Ssb2–Trt1 interaction, double-mutants maintained telomeres even after extensive restreaking, although *E86A* showed additional telomere shortening in this background (Fig 5D). These results, together with the synergistic telomere loss observed when *tpz1-K75A* (Trt1–Tpz1 interface) was combined with mutants affecting either the Ssb2–Tpz1 or Ssb2–Trt1 interfaces (Fig S8B-C), support the view that the Ssb1–Trt1, Ssb2–Trt1, Ssb2–Tpz1, and Trt1–Tpz1 interfaces make partially redundant but non-overlapping contributions that collectively promote productive telomere extension by telomerase.

Notably, *ssb1-W523C*, *ssb2-W112A,* and *tpz1-R81E* differ from other interface mutants in that each causes complete telomere loss on its own. This suggests that those mutations may affect not only the specific interfaces predicted by the model, but also additional aspects of telomere maintenance, such as telomerase recruitment (i.e., *ssb2-W112A*), substrate positioning, or coordination of telomerase action with replisome dynamics at telomeres.

### Shelterin mutations allow telomerase-dependent maintenance of telomeres in *ssb1-W523C*, but not *tpz1-R81E*

The severe telomere-loss phenotypes of *ssb1-W523C* and *tpz1-R81E* raised the question of why telomerase, even when efficiently recruited, fails to maintain telomeres in these mutants. To gain further mechanistic insight, we investigated whether shelterin-related mutations that cause elongated telomeres could suppress telomere loss in these mutant backgrounds.

We found that combining *ssb1-W523C* with *tpz1-K242R*, *tpz1-AWAAA*, *poz1Δ*, or *rap1Δ* enabled *W523C* cells to maintain stable telomeres in a telomerase-dependent manner (Fig S9A-C). However, telomeres remained very short, indicating that telomerase was still less efficient than in wt cells. SUMOylation of Tpz1-K242 promotes recruitment of Stn1–Ten1–Polα [10–12], and the SWSSS motif in Tpz1 contributes redundantly to Tpz1–Stn1 interaction while also playing an additional role in telomerase inhibition [12]. Likewise, both *poz1Δ* and *rap1Δ* substantially delay the recruitment of Stn1–Ten1–Polα to telomeres during late S/G_2_ [50]. Thus, the common feature of these *W523C* suppressors is prolonged and enhanced accumulation of ssDNA, RPA, and telomerase at telomeres, which may provide additional time for a partially impaired telomerase complex to extend telomeres.

In contrast, none of these telomerase-dependent suppressors rescued *tpz1-R81E* (Fig S9D-E), indicating that increased RPA accumulation is not sufficient to bypass the activation defect in *tpz1-R81E* cells. We did observe that *taz1Δ* allows both *ssb1-W523C* and *tpz1-R81E* cells to maintain stable telomeres (Fig S9F-G), but suppression also occurred in a *trt1Δ* background, suggesting that this rescue may reflect the homologous recombination (HR)-based mechanism of telomere maintenance, analogous to *trt1Δ taz1Δ* cells [35, 51, 52].

These results distinguish two types of defects: *ssb1-W523C*, which can be partially overcome by prolonging the time window for telomerase action, and *tpz1-R81E*, which appears to be more fundamentally defective in the telomerase activation step itself.

### RPA–Trt1, RPA–Tpz1, and Tpz1-Trt1 interactions are required for catalytically inactive telomerase to inhibit recombination-dependent telomere maintenance

The HR-dependent mechanism that maintains telomeres in *trt1Δ taz1Δ* cells requires Tel1^ATM^, Rad51, Rad52, Mre11–Rad50–Nbs1 (MRN) complex, and Rap1 [35, 51, 52]. Because Taz1 suppresses telomere recombination, reintroduction of a *taz1*^+^ plasmid causes telomere loss and chromosome circularization of *trt1Δ taz1Δ* survivors [51, 52]. Previous work further showed that plasmids expressing catalytically inactive Trt1 (*D743A* or *D590A*) or a Trt1-ΔT/RT truncation construct lacking the RT domain can likewise inhibit recombination and force telomere loss in *trt1Δ taz1Δ* cells [51, 52] (Fig 5E-F). Notably, these Trt1 mutant constructs all retain the wt TEN-domain, which is predicted to mediate Trt1 interactions with RPA and Tpz1.

We therefore asked whether the predicted RPA–Trt1–Tpz1 interaction interfaces are required for catalytically inactive Trt1 to inhibit recombination. When Trt1-D743A was introduced into *trt1Δ taz1Δ* cells carrying additional mutations that disrupt the Ssb1–Trt1 (*ssb1-W523C*), Trt1–Tpz1 (*tpz1-K75A*), or Ssb2–Tpz1 (*tpz1-R81E*) interfaces, the plasmid failed to cause telomere loss (Fig 5F-G). Importantly, these same mutations did not prevent telomere loss induced by reintroduction of Taz1, indicating that the defect is specific for telomerase-dependent inhibition of telomere recombination.

Similarly, introducing additional Trt1 mutations predicted to disrupt the Ssb1–Trt1 (E116A or R119A) or Ssb2–Trt1 (E86A or D157A) interfaces rendered the Trt1-D743A plasmid ineffective at forcing telomere loss in *trt1Δ taz1Δ* cells (Fig 5H). These results provide an independent functional test of the AlphaFold3 model and show that RPA–Trt1, RPA–Tpz1, and Trt1–Tpz1 interactions are required for catalytically inactive telomerase to inhibit recombination-dependent telomere maintenance. Thus, the same interaction network that promotes productive telomere extension in telomerase-positive cells also enables catalytically dead telomerase to block recombination-based telomere maintenance.

### RPA–Trt1–Tpz1 regulation of telomerase is likely conserved in budding yeast and humans

Because Ssb1-W523 is highly conserved across eukaryotes, we next asked whether the equivalent residue in budding yeast, Rfa1-W533, also contributes to telomere maintenance. Freshly generated *rfa1-W533C* cells showed progressive loss of cell viability, followed by the emergence of survivor cells that maintained short telomeres, in some cases with evidence of subtelomeric rearrangements (Fig 6A-B). Combining *rfa1-W533C* with *rad52Δ* caused much earlier and more severe loss of viability (Fig 6C), indicating that Rad52-dependent telomere recombination [53] improves survival of *rfa1-W533C* cells. ChIP assays further showed that the budding yeast telomerase regulatory subunit Est1 is recruited to telomeres at levels similar to those in wt in *rfa1-W533C* cells (Fig 6D). Thus, as in fission yeast, the primary defect is not telomerase recruitment but rather productive telomerase action after recruitment. Since early generation *rfa1-W533C* cells grew comparably to wt cells (Fig 6C) and did not show severe sensitivity to MMS, HU, or CPT (Fig S10), we believe that this mutation does not severely affect RPA’s general replisome function during telomere replication or global DNA damage responses.

**Fig 6.**
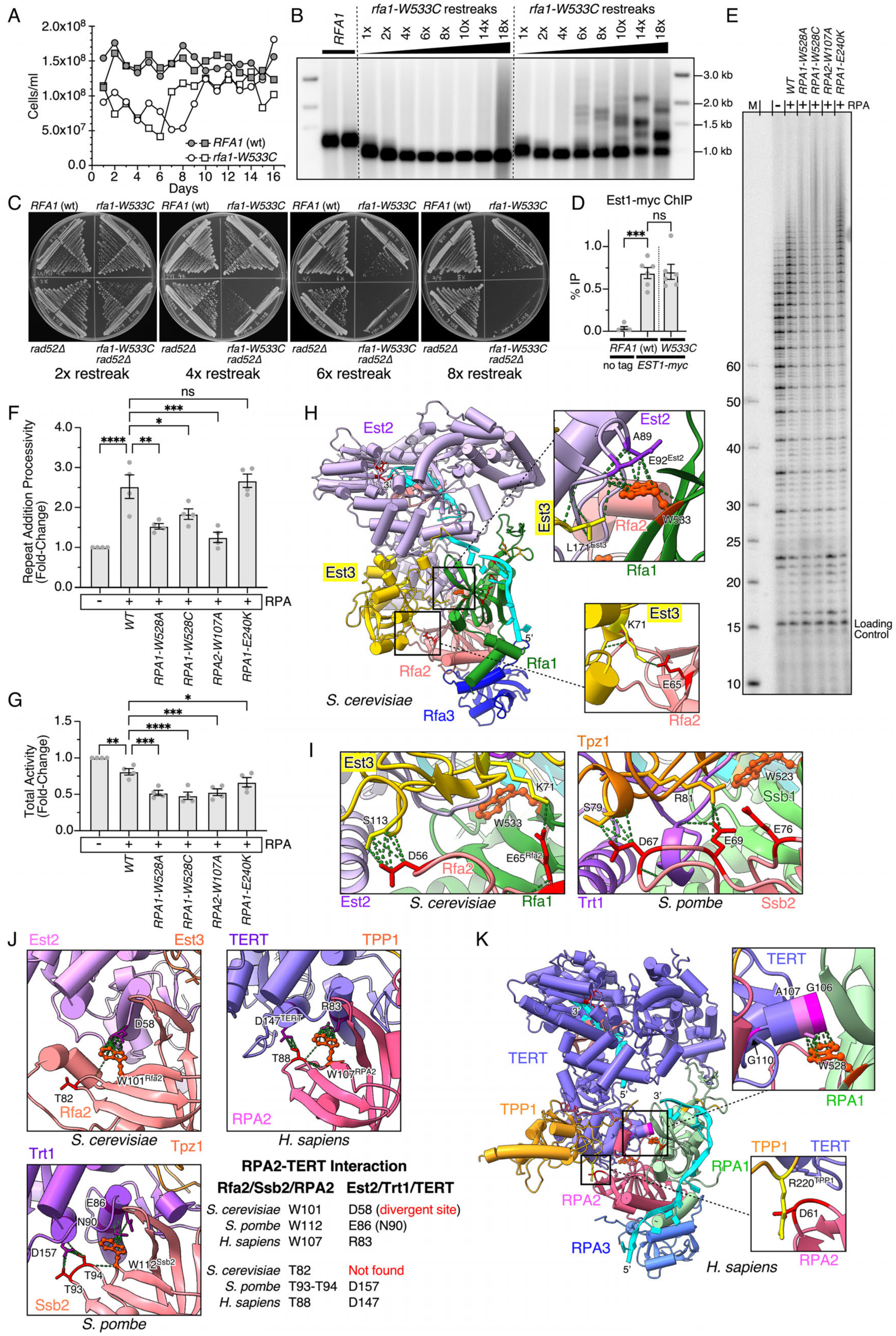
The Rfa1/RPA1 residue equivalent to Ssb1-W523 contributes to telomerase function in budding yeast and humans. **(A)** Growth curve analysis of budding yeast *rfa1-W533C* strains, carrying the equivalent mutation to fission yeast *ssb1-W523C*. **(B)** Telomere length analysis by Southern blot of two independent *rfa1-W533C* clones after successive restreaks. **(C)** Cell viability assay of budding yeast *rfa1-W533C rad52Δ* cells. YPDA plates from the indicated restreaks were photographed after incubation at 30°C for 2 days. **(D)** ChIP analysis of telomere binding by the budding yeast telomerase subunit Est1-myc in *RFA1* and *rfa1-W553C* strains. Statistical significance was assessed by ANOVA with Dunnett’s comparison test (ns stands for not significant; ***P ≤ 0.001). See S1 Data for individual % IP values and statistical analysis. **(E-G)** Direct telomerase assay showing effects of wt and mutant human RPA complexes on telomerase activity and processivity. The *in vitro* telomerase assay PAGE (E) is quantified to monitor changes in repeat addition processivity (F) and total activity (G). Previously established strongest RPA processivity mutant *RPA2-W107A,* as well as *RPA1-E240K*, which causes telomere shortening but does not affect processivity, were included as controls. For (F-G), Statistical significance was assessed by ANOVA with Dunnett’s comparison test (ns stands for not significant; *P ≤ 0.05; **P ≤ 0.01; ***P ≤ 0.001; ****P ≤ 0.0001). See S1 Data for individual values and additional statistical analysis. **(H)** AlphaFold3 model for budding yeast RPA–Est2–Est3 complex interactions. Rfa1 W533 and Est3 L171 interact with adjacent sites on Est2 (top inset), and Rfa2 E65 interacts with Est3 K71 (bottom inset). **(I)** Comparison of Ssb2/Rfa2 to Tpz1/Est3 interactions between fission and budding yeast. Showing K71^Est3^–E65^Rfa2^ and S113^Est3^–D56^Rfa2^ interactions in budding yeast (left panel), and R81^Tpz1^–E69^Ssb2^ and S79^Tpz1^–D67^Ssb2^ interactions in fission yeast (right panel). **(J)** Detailed structures of Rfa2-Est2-Est3 interface in budding yeast, fission yeast (Ssb2–Trt1–Tpz1), and human (RPA2–TERT–TPP1). Corresponding RPA2–TERT interactions are indicated. **(K)** AlphaFold3 model of human RPA–TERT–TPP1complex. W528^RPA1^–G106/A107^TERT^ (top panel) and D61^RPA2^–R220^TPP1^ (bottom panel) interactions are shown.

Human RPA functions as a telomerase processivity factor [26]. We therefore also tested whether the human equivalent residue, RPA1-W528, contributes to this RPA function. Direct telomerase assays revealed that purified RPA complexes containing *RPA1-W528A* or RPA1-*W528C* showed a substantially reduced stimulation of telomerase processivity, with the W528A mutation having a stronger effect than W528C and an effect comparable to that of *RPA2-W107A* [26] (Fig 6E-G). Together, these results indicate that RPA1-W528 contributes to RPA-mediated stimulation of telomerase processivity. However, future studies, such as the establishment of cell lines expressing RPA1-W528 mutant alleles, will be required to evaluate the impact of RPA1-W528 mutations on telomere maintenance in human cells.

Because RPA’s role in modulating telomerase activity after recruitment appears to be broadly conserved, we expanded our AlphaFold3 structural modeling analysis for RPA–Trt1–Tpz1-like complexes to budding yeast and humans. AlphaFold3 models of the budding yeast RPA-Est2^TERT^–Est3^TPP1^ complex (Figs 6H and S11A-D) suggested that Rfa1-W533 contacts Est2 at an interface analogous to the Ssb1–Trt1 interface identified in fission yeast (Fig 3B). Notably, the predicted RPA-Est2-Est3 structure was consistent with a recently published cryo-EM structure of the Est2-Est3 interface, and the predicted RPA-interacting surfaces are available to interact with RPA in the context of the holoenzyme complex, consisting of TLC1 RNA, Est1-3, and Pop1, 6, and 7 subunits [54] (Fig S11E). Interestingly, the budding yeast Est3-L171 residue, previously classified as a TELR residue [55], was predicted to contact either the Est2 region adjacent to Rfa1-W533 or the Rfa1-W533 itself, suggesting that Est3-L171 may collaborate with Rfa1-W533 in stabilizing the RPA-Est2-Est3 complex (Figs 6H and S11F).

The AlphaFold3 model also predicted that Est3-K71, another residue classified as TELR [55], interacts with Rfa2-E65 (Fig 6H), consistent with recent work showing reciprocal rescue between *est3-K71E* and *rfa2-E65K* mutations [22]. Although the residues linking the RPA2 subunit to the TPP1-like protein have diverged, this interaction appears functionally analogous to the R81^Tpz1^-E69^Ssb2^ interaction in fission yeast (Figs 6I and S3D).

The two Ssb2–Trt1 interactions identified in fission yeast, W112^Ssb2^–E86^Trt1^ and T93/94^Ssb2^–D157^Trt1^, appear to be well conserved in humans [26] (Fig 6J). In budding yeast, Rfa2-W101 is predicted to interact with Est2-D58 (Fig 6J), although this contact is shifted relative to the equivalent interface in fission yeast and humans. In contrast, Rfa2-T82, the equivalent of Ssb2-T93/T94 and human RPA2-T88, is not predicted to contact Est2 (Fig 6J). Although the budding yeast *rfa2-W101K* mutant exhibits substantial telomere shortening [22], the phenotype is much milder than the complete telomere loss caused by *ssb2-W112A* in fission yeast or the loss of telomerase stimulation activity caused by human *RPA2-W107A* mutant [26]. Likewise, mutations at Rfa2-W101 do not cause strong growth defects in budding yeast [56, 57], unlike *ssb2-W112A* in fission yeast, suggesting that this residue is less critical for overall RPA function in budding yeast.

Finally, AlphaFold3 modeling of the human RPA–TERT–TPP1 complex [26] (Figs 6K and S12) predicts that RPA1-W528 contacts G106 and A107 in TERT. Notably, the disease-associated *TERT-G106W* mutation causes a severe reduction in telomerase activity in patients with cryptic *Dyskeratosis Congenita* (cDKC) [58], and nearby TERT mutations, *G110A* and *V119M*, likewise cause severe telomere shortening [59]. Thus, these observations support the idea that the RPA1–TERT interface is also functionally important in humans.

Collectively, the experimental data and structural modeling presented in this study support a model in which coordinated RPA–TERT and RPA–Tpz1/TPP1 interactions enable telomerase to productively extend telomeric ssDNA in a wide variety of organisms.

## Discussion

### The RPA–Trt1–Tpz1 module links telomerase recruitment to productive telomere extension

Earlier work in budding yeast and fission yeast suggested that the RPA complex contributes to telomere maintenance and may collaborate with telomerase and telomere-associated proteins [21, 33]. However, a mechanistic explanation for these observations remained unclear. Evidence that an RPA-like complex can function as a core telomerase cofactor first emerged from studies of *Tetrahymena thermophila*, where cryo-EM structures revealed that the RPA-like TEB complex forms an integral component of the telomerase holoenzyme and promotes repeat-addition processivity by positioning telomeric ssDNA for nucleotide addition [28, 46, 48]. Whether canonical RPA performs an analogous function in other eukaryotes remained uncertain until recent studies demonstrated that budding yeast RPA2 interacts with the TPP1 ortholog Est3 to regulate telomerase activity [22], and that human RPA stimulates telomerase processivity through a defined RPA2-TERT interaction [26].

In this work, we identify a ternary RPA–Trt1–Tpz1 complex in fission yeast that couples telomerase recruitment to productive telomere extension. Genetic dissection of multiple interaction interfaces, combined with AlphaFold3-guided structural modeling, supports a model in which RPA directly contributes to telomerase activation at chromosome ends. Most notably, the *ssb1-W523C* mutation uncouples telomerase recruitment from productive telomere extension, as *ssb1-W523C* cells accumulate robust levels of RPA, Rad26^ATRIP^, and Trt1^TERT^ at progressively shortening telomeres, yet ultimately lose all telomeric repeats and circularize chromosomes (Fig 2A-D). Thus, in this mutant background, telomerase is recruited but unable to extend telomeric ends. This separation-of-function phenotype demonstrates that RPA is not merely a passive ssDNA-binding factor but plays an active role in telomerase activation.

Our AlphaFold3-guided genetic analysis supports the existence of four functionally important interaction surfaces, Ssb1–Trt1, Ssb2–Trt1, Ssb2–Tpz1, and the TEL-patch mediated Trt1–Tpz1 interaction, that together define a regulatory network linking RPA to telomerase activation. These interactions likely enable assembly of an RPA–Trt1–Tpz1 complex that promotes productive engagement of the telomeric 3’ end with the telomerase catalytic center (Figs 3 and 7A). Consistent with this model, the strongest telomere-maintenance defects observed in this study map to residues within the DBD-C domain of Ssb1 and the DBD-D domain of Ssb2, domains positioned to coordinate ssDNA binding with interaction surfaces involving Trt1 and Tpz1. In particular, *ssb1-W523C*, *ssb2-W112A*, and *tpz1-R81E* each disrupt residues predicted to participate in these interfaces and each cause complete telomere loss in fission yeast, but the underlying defects are not identical.

**Fig 7.**
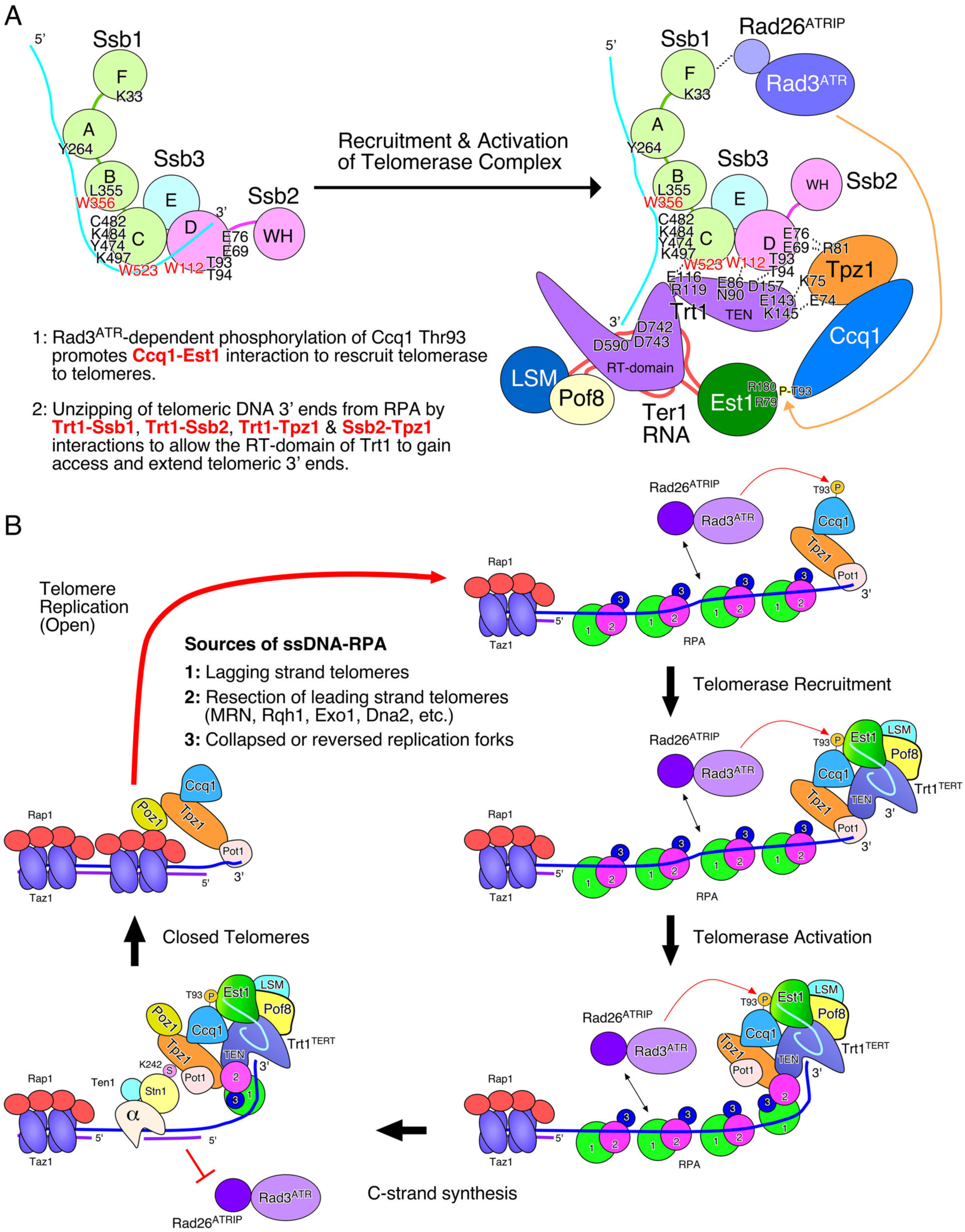
Working model for fission yeast telomere regulation by RPA and shelterin. **(A)** Model illustrating the change in interactions when RPA is bound to telomeric ssDNA (left) versus when RPA engages Trt1 and Tpz1 (right). Residues identified in this study as contributing to telomere maintenance and protein-protein interactions are indicated. **(B)** A working model for how RPA-coated telomeric ssDNA 3’ overhangs promote telomerase recruitment and activation. RPA bound to ssDNA recruits Rad3^ATR^–Rad26^ATRIP^, facilitating Ccq1-T93 phosphorylation and subsequent telomerase recruitment. Direct protein-protein interactions among RPA, Trt1, and Tpz1 promote the release of the 3’ ssDNA end into the telomerase catalytic site, enabling telomere extension. Recruitment of Pot1 and Stn1–Ten1–Polα promotes C-strand synthesis and completion of the replication cycle.

ChIP assay showed that *ssb2-W112A* exhibits reduced accumulation of Ssb2 at short telomeres and severely diminished Trt1 association in early-generation cells (Fig 4H-I), indicating that this mutation compromises telomerase recruitment. Thus, *ssb2-W112A* is not a clean separation-of-function activation mutant, but rather a particularly severe allele that disrupts both RPA association with telomeres and the predicted Ssb2–Trt1 interface. In contrast, *ssb1-W523C* and *tpz1-R81E* are better interpreted as post-recruitment mutants, because both retain increased Trt1 binding to shortening telomeres before eventual telomere loss (Figs 2D and 4B), indicating that telomerase recruitment occurs, but productive telomere extension is impaired. The distinct suppression profiles of *ssb1-W523C* and *tpz1-R81E* indicate that the Ssb1–Trt1 and Ssb2–Tpz1 interfaces make qualitatively different contributions within the ternary complex. Suppression of *ssb1-W523C* by mutations that prolong the persistence of RPA-coated ssDNA suggests that the Ssb1–Trt1 interaction primarily enhances the efficiency of telomerase within a limited temporal window, allowing telomerase to overcome inhibitory effects imposed by shelterin and Stn1–Ten1. In contrast, the inability of these suppressors to rescue *tpz1-R81E*, except through restoration of the predicted electrostatic interaction with Ssb2, suggests that the Tpz1-Ssb2 interface plays a more fundamental role in enabling productive engagement of telomeric ssDNA by telomerase.

These interactions function together with the previously described TEL-patch-mediated Trt1–Tpz1 interaction [8, 9] to promote productive extension of telomeric ssDNA with telomerase following recruitment through the Rad3^ATR^/Tel1^ATM^–Ccq1–Est1 pathway [6]. Comparative analysis of AlphaFold3-predicted structures further suggests that analogous interaction networks operate in budding yeast and humans (Figs 6 and S13). Taken together, these observations support a model in which RPA acts as a conserved architectural component of telomerase regulatory assemblies, coordinating interactions between TERT and TPP1-like factors to enable efficient telomere elongation.

### A model for RPA- and Tpz1-mediated telomerase regulation during DNA replication

Integrating our findings with prior work, we propose a working model describing how RPA accumulation at telomeres is regulated during DNA replication and how elevated RPA levels facilitate telomerase-mediated telomere extension (Fig 7B). In fission yeast, telomeric RPA-coated ssDNA arises from multiple replication-coupled processes during S-phase. At lagging-strand replicated telomeres, ssDNA accumulates because lagging-strand synthesis is delayed relative to the arrival of the replicative CMG helicase and the leading-strand DNA polymerase Polε [13, 50]. At leading-strand replicated telomeres, the MRN complex, together with the RecQ helicase Rqh1, Exo1, and Dna2, generates an RPA-coated 3’ ssDNA overhang [60–62]. Replication fork stalling, reversal, or collapse near chromosome ends may further contribute to the accumulation of RPA-coated ssDNA at telomeres [63].

Accumulation of RPA-coated ssDNA increases dramatically as telomeres shorten (Fig 2), promoting Rad3^ATR^–Rad26^ATRIP^ association and phosphorylation of Ccq1-T93, which in turn stimulates the Ccq1-Est1 interaction and preferential recruitment of telomerase to shorter telomeres [6, 42, 50]. Several mechanisms likely contribute to this length-dependent feedback. Shorter telomeres contain fewer Taz1 binding sites, leading to altered replication timing and increased exposure of ssDNA [50, 63, 64]. In addition, shorter telomeres may also undergo increased MRN-dependent resection due to reduced Ccq1 binding (Fig 2G) [65]. However, this positive feedback has a lower limit: when telomeres become critically short, shelterin can no longer accumulate efficiently, leading to loss of both Ccq1 and telomerase binding despite persistent RPA and ATR association (Figs 2D, 2E, 2G and 4B).

RPA likely promotes telomere extension through several distinct mechanisms. First, it maintains the telomeric 3’ overhang in an accessible single-stranded conformation by resolving secondary structures, such as G-quadruplexes [34, 66, 67]. Second, it facilitates telomerase recruitment by promoting assembly of Rad3^ATR^–Rad26^ATRIP^ and the subsequent Ccq1–Est1 interaction [42, 68]. Third, and most central to this study, formation of the RPA–Trt1–Tpz1 complex likely promotes transfer of the telomeric 3’ end from the canonical RPA-bound state to the telomerase catalytic site (Figs 3 and 7). In this model, coordinated interactions among RPA, Tpz1, and Trt1 position the substrate for productive extension while limiting inappropriate access of recombination factors such as Rad52.

As telomerase extends the 3’ end, the RPA–Trt1–Tpz1 complex may translocate along telomeric DNA, leaving newly synthesized ssDNA behind. The fate of this ssDNA likely determines whether telomere extension continues or terminates. Binding by additional RPA and ATR complexes may reinforce signaling to Ccq1 and telomerase retention, whereas binding by Pot1 promotes transition to a protected, telomerase-inaccessible state. This transition is coordinated with C-strand synthesis by the Stn1–Ten1–Polα complex. As C-strand synthesis proceeds, RPA is displaced, the overhang is converted to dsDNA, and Taz1 binding is restored, promoting Rap1-Poz1 assembly and Tpz1 SUMOylation, which further enhances Stn1–Ten1–Polα recruitment [12, 18]. Ultimately, the remaining G-rich overhang is bound by Pot1, restoring a fully protected telomere state (Fig 7B).

Replication-coupled accumulation of telomeric RPA-coated ssDNA is conserved across eukaryotes, and S-phase-specific accumulation of telomeric overhangs has been well documented [69–71]. In these systems, lagging-strand telomeres accumulate ssDNA due to delayed fill-in synthesis, whereas leading-strand telomeres undergo MRN-mediated resection [71–74]. This conservation suggests that the RPA-dependent telomerase activation mechanism described here may operate broadly across eukaryotes.

Proteomic analyses of replicating human telomeres show enrichment of RPA, TPP1, and hTERT, but reduced levels of POT1 [75]. In addition, POT1 appears to be dispensable for telomerase-mediated telomere extension in mouse embryonic stem cells [76]. Together with evidence that POT1 primarily restrains telomerase by promoting CST-mediated C-strand synthesis [71], these findings support a model in which RPA–TERT–TPP1 assemblies play a central role during active telomere extension, whereas POT1 primarily terminates extension and restores the protected state.

### Evolutionary conservation of the RPA-TERT-TPP1 complex

All four interaction interfaces identified in the fission yeast RPA–Trt1–Tpz1 complex have predicted counterparts in budding yeast and humans (Fig S13). Although direct structural validation remains limited, AlphaFold3 modeling provides a framework for rationalizing and re-evaluating decades of prior genetic and biochemical observations across these systems.

Despite extensive sequence divergence among Tpz1, Est3, and TPP1, structural modeling suggests that these proteins share a conserved fold and overall architecture. A key feature emerging from these structure model-based comparisons is that, while the global organization of the ternary complex appears conserved, specific residue-level interactions have been extensively rewired. Equivalent residues are often predicted to engage different partners, or distinct regions of the same partner, thereby generating organism-specific interaction modes within a conserved functional module (Fig S3 and S13C).

Several selected examples illustrate this principle. The TEL-patch residues E74 and K75 of Tpz1 occupy positions structurally equivalent to E168 and E169 on human TPP1, yet are predicted to contact different regions of their respective telomerase catalytic subunits, namely K145 and E143 on fission yeast Trt1 (Fig S3C-D and S6D) versus R774 and Y772 on human TERT (Fig S13B) [77, 78]. The budding yeast Est3 TEL-patch residue E114 interacts with Est2-K62 [54] (Fig S13A), whereas the structurally analogous Tpz1-R81 residue in fission yeast does not contact Trt1 but instead reaches across to Ssb2-E69 (Fig 4A and S3D).

Against this backdrop of interface reshuffling, one class of contacts appears to have remained particularly stable: residues equivalent to Ssb1-W523 and Ssb2-W112 are consistently predicted to participate in RPA–TERT interactions across all three organisms. Importantly, these predicted contacts are structurally incompatible with the simultaneous binding of these residues to ssDNA, and studies supported their positive contribution to telomere maintenance in budding yeast and humans (Fig 6) [22, 26]. These tryptophan residues of RPA, therefore, appear to define the most deeply conserved elements of the RPA–TERT interfaces.

The AlphaFold3 model of the budding yeast RPA-Est2-Est3 complex also offers a useful framework for interpreting earlier Rfa2 and Est3 mutagenesis data. For example, *rfa2*-*D56K* causes severe telomere shortening and replicative senescence [22], and AlphaFold3 predicts that Rfa2-D56 interacts with Est3-S113, a contact that appears to be conserved in fission yeast, where the equivalent Ssb2-D67 contacts Tpz1-S79 (Fig 6I). Interestingly, despite being predicted to interact with Rfa2-D56, *est3-S113R* shows no telomere defects [55], likely because this mutation retains favorable ionic interactions with Rfa2-D56. It will therefore be interesting to test whether *est3-S113D* causes a stronger telomere phenotype and whether it can suppress the senescence phenotype of *rfa2-D56K*. Analogous experiments could test the functional relevance of the predicted equivalent interaction in the fission yeast ternary complex.

Remarkably, AlphaFold3 predicts that human RPA2-D61 (equivalent to Ssb2-D67 and Rfa2-D56) also contacts TPP1-R220 within the RPA–TERT–TPP1 complex (Fig 6K and S12D), suggesting unusually strong evolutionary conservation of this RPA2-centered interaction node (Fig S13C). TPP1-R220 lies within the TELR region (R218-V221), originally identified by structural comparison with budding yeast Est3 and thought to be functionally analogous to the Est3-L171 region [79]. Mutations in the TPP1 TELR region impair TPP1-mediated telomerase stimulation and disrupt telomere maintenance in human cells without affecting telomerase recruitment [79], supporting the relevance of the predicted RPA2–TPP1 interface. At the same time, Est3 L171 is predicted to support the W533^Rfa1^–Est2 interface rather than contacting Rfa2 (Figs 6H and S9E). Thus, conserved TELR sites may contribute to telomere maintenance through distinct mechanistic routes across organisms.

Further expanding on prior experimental and modeling work on the human RPA–TERT–TPP1 complex [26], AlphaFold3 also suggested that TERT-R132 and TERT-E113 may interact, potentially stabilizing the TERT region predicted to interact with RPA1-W528 (Fig S12E). TERT mutations near the predicted RPA1–TERT interface impair telomerase function in human cells [58, 59]. In addition, the disease-associated *TERT*-*R132D* mutation disrupts telomerase recruitment [80] and likely telomerase activation as well, because forced targeting of TERT to telomeres does not rescue its telomere maintenance defect [81]. Interestingly, whereas cryo-EM structures of the TERT–TPP1 complex place TPP1-E215 near both TERT-K78 and R132 [77], AlphaFold3 ternary complex models predict an interaction with K78 only, potentially freeing TERT-R132 to engage TERT-E113. The region of TERT predicted to contact RPA1-W528 is currently unresolved in available TERT–TPP1 cryo-EM structures [77, 78], raising the possibility that including RPA in future structural analyses might stabilize this region and enable direct visualization of the proposed RPA1–TERT interaction interface.

### Limitations of the study

The consistency of our findings in fission yeast with prior studies in budding yeast and humans suggests that RPA plays a conserved role in telomerase regulation across eukaryotes. However, several limitations should be noted. Although data from current and previous studies support the proposed interaction network, high-resolution structural determinations of RPA–TERT–TPP1/Tpz1/Est3 complexes remain lacking. In addition, key mechanistic steps, such as transfer of the telomeric 3’ end from RPA to the telomerase catalytic center, are inferred rather than directly observed. Future structural and biochemical studies will therefore be required to test the validity of these predicted assemblies. Nevertheless, the convergence of genetic, biochemical, and comparative structural modeling evidence supports a model in which canonical RPA functions not only as an ssDNA-binding factor and checkpoint platform, but also as a conserved architectural component of telomerase-regulatory complexes that couple telomerase recruitment to productive chromosome-end extension while limiting recombination-based telomere maintenance.

## Materials and methods

### Yeast strains, plasmids, and primers

Yeast strains used in this study were constructed using standard methods [82–85]. *S. pombe* cells were cultured in YES (Yeast Extract with Supplements) medium at 32°C. For most fission yeast strains, mutations or epitope tags in RPA or Trt1 were introduced by transforming a haploid strain. For *ssb2-W112A* strains, the mutation (with or without an epitope tag) was introduced into diploid strains; haploid progenies were obtained by sporulation in supplemented SPA medium (SPAS) followed by tetrad dissection using a Zeiss dissection microscope. To combine mutations or epitope-tagged alleles, genetic crosses between opposite mating-type strains were performed in SPAS medium, and haploid progenies with desired genotypes were identified. Newly generated strains were repeatedly streaked on YES agar plates to obtain terminal telomere phenotypes.

*S. cerevisiae* strains were grown at 30°C in YPDA (Yeast Peptone Dextrose Adenine) medium. The *rfa1-W533C* mutation was introduced into haploid or diploid budding yeast strains by transformation. Double mutants (*rfa1-W533C rad52Δ* or *rfa1-W533C Est1-myc*) were generated by transforming a diploid *rfa1-W533C/RFA1* strain with the corresponding constructs, followed by sporulation and selection of haploid progenies with desired genotypes.

Point mutations and truncations were generated with Q5 PCR (NEB, M0491L), Gibson Assembly (NEB, E5510S), or site-directed mutagenesis kits Q5 (NEB, E0554S) or QuikChange Lightning (Agilent, 210519) on plasmids carrying the gene of interest. All constructs were verified by whole-plasmid sequencing (Plasmidsaurus, Louisville, KY). pRIT2-ssb1-mutant plasmids were generated and sequence verified by Genscript (Piscataway, NJ). To integrate various mutant alleles, mutant genes were either excised from plasmids or amplified by PCR and transformed into fission yeast or budding yeast cells. Correct genomic integration was verified by PCR and DNA sequencing.

Fission yeast and budding yeast strain genotypes are listed in Table S1 and Table S2, respectively. Plasmids used in this study are listed in Table S3. DNA primers used in this study are listed in Table S4. The sources of previously published mutants or epitope-tagged constructs used in this study and additional details on the construction of various newly generated alleles from this study are listed in Table S5.

### Construction of elongated telomere strains

Isogenic fission yeast strains with progressively shortening telomere lengths were generated by crossing *rap1Δ* with *poz1Δ* strains carrying epitope-tagged and/or mutant alleles, followed by dissection and selection of *rap1^+^ poz1^+^* progenies (Fig S5A). Because *rap1Δ* and *poz1Δ* strains carry highly elongated telomeres [5, 86], the resulting strains initially carry elongated telomeres. Strains were then restreaked successively, and progressive telomere shortening was monitored by telomere-length Southern blot analysis (See Fig S5B for an example of *trt1-D473A-myc* restreaks.) For each restreak, cells were processed to make a glycerol stock at -80°C to allow recovery of these strains with defined telomere length for later experimental use.

To allow telomere elongation in *rap1Δ* or *poz1Δ* strains with *trt1-D743A* or *trt1Δ*, a plasmid that expresses wt Trt1 (pREP-trt1^+^) was utilized (Fig S5B-C), while a plasmid that expresses wt Tpz1 (pREP-tpz1^+^) was utilized for *poz1Δ* strains carrying *tpz1-R81E* or *tpz1-K75A* (Fig S5D). For *rap1Δ* or *poz1Δ* strains carrying *ssb1-W523C*, a plasmid that expresses wt Ssb1 (pIRT2-ssb1^+^) was utilized to elongate telomeres (Fig S5E).

### Growth curve analysis

Budding yeast cells were grown in YPDA at 30°C, diluted to 5×10^5^ cells/mL in fresh medium, and cell density was measured every 24 hours over multiple days, as described previously [87].

### Chromatin immunoprecipitation (ChIP)

ChIP assays were performed as described previously [88]. Exponentially growing cells were crosslinked with 1% formaldehyde by addition of 1/10 volume of fixation solution (11% formaldehyde, 100 mM NaCl, 1mM EDTA pH 8.0, 0.5 mM EGTA pH 8.0, 50 mM Tris-HCl pH 8.0) for 20 min at room temperature, incubated for an additional 5 min after addition of Glycine at a final concentration of 125 mM. Cells were washed 3× with TBS (20 mM Tris-HCl, pH 7.6, 150 mM NaCl), pelleted, and frozen in liquid nitrogen.

Cells were lysed in lysis buffer (50 mM Hepes-KOH pH 7.5, 140 mM NaCl, 1mM EDTA, 1% Triton X-100, 0.1% sodium deoxycholate, Roche complete protease inhibitor cocktail) with glass beads using FastPrep-24 5G (MP Biomedicals), followed by sonication for 3x 30sec (Bioruptor Pico, Diagenode). Cleared lysates were incubated with either monoclonal anti-myc (9B11, Cell Signaling, 58730), anti-FLAG (M2 F1804, Sigma), or anti-GFP (Sigma, 11814460001) antibodies, followed by capture with Dynabeads Protein G (ThermoFisher, 10009D).

ChIP DNA samples were analyzed by qPCR (BioRad CFX Duet) with PerfeCTa SYBR Green SuperMix (Quantabio, 95054-500), amplified with either fission yeast primers 637 and 638, or budding yeast primers 2371 and 2372 (Table S4). Values were normalized to input DNA samples and expressed as %IP. Error bars represent SEM.

### Southern blot analysis

Genomic DNA was prepared by resuspending cell pellets in equal volumes of DNA buffer (100 mM Tris pH 8.0, 1 mM EDTA pH 8.0, 100 mM NaCl, 1% SDS) and Phenol:Chloroform:Isoamyl alcohol (25:24:1), and lysing cells with glass beads using FastPrep-24 5G (MP Biomedicals). Genomic DNA (1 μg) was digested overnight with EcoRI (fission yeast) or XhoI (budding yeast), separated on 1% agarose gel, and transferred to Nytran SPC membranes (Cytiva).

Fission yeast telomeres were detected using a ^32^P-labeled telomeric repeat probe generated by random priming of a ApaI-SacI telomere repeat DNA fragment, derived from pTELO plasmid (Table S3). Budding yeast telomeres were detected using ^32^P-labeled Y’ probe generated by random priming of a PCR product amplified from budding yeast genomic DNA with primers 2220 and 2221 (Table S4). Blots were imaged with Amersham Typhoon Phosphorimager (Cytiva).

### Western blot analysis

Western blot analysis was performed using monoclonal anti-myc (9B11, Cell Signaling 58730), anti-FLAG (M2 F1804, Sigma), or anti-Ssb1 [30], and monoclonal anti-α-tubulin (clone B-5-1-2 T5168, Sigma) as primary antibodies. As secondary antibodies, either HRP-conjugated (goat) anti-mouse (ThermoFisher, 31430) or IRDye 800CW Goat anti-Mouse IgG (Licor, 926-32210) were used. Blots were imaged with either an Azure 400 imager for HRP detection or an LI-COR Odyssey CLx imager for fluorescent detection.

### Fluorescence microscopy

For live-cell microscopy, cells expressing Ssb3-GFP were cultured to <0.8 OD_600_ in minimal media (PMG +HUAL) at 25°C in three independent experiments. Cells were then concentrated and spotted onto clear glass slides.

15-20 Z-stacks/frame (Z-step size 0.2-0.27µm) were acquired using a Yokogawa Spinning Disk confocal Leica DM8i inverted microscope with Leica CS apochromatic 100× oil objective 1.43 NA. Prime 95B Back-illuminated sCMOS camera and Leica LAS-X software version 3.7.x were used. Maximum intensity Z-stack projections were analyzed in ImageJ and the percentages of nuclei with distinct GFP-foci were quantified for >200 individual cells on average per experiment.

### AlphaFold3 modeling

AlphaFold3 models were generated using a web server (https://alphafoldserver.com) for the indicated proteins and analyzed with UCSF ChimeraX [89]. For *S. pombe* complex modeling, full-length Trt1, full-length Ssb1, Ssb2, and Ssb3, and Tpz1 corresponding to amino acids 1-219 were used. For *S. cerevisiae* complex modeling, all proteins used were full-length. For human complex modeling, full-length TERT, TPP1 corresponding to amino acids 87-266 (or 1-180 with a more recently revised TPP1 amino acid residue number) [90], full-length RPA1 and RPA3, and RPA2 corresponding to amino acids 1-175. The resulting representative AlphaFold3 server results are shown in Fig S14 to help evaluate the confidence of these models based on pIDDT values. For PDB accession numbers for previously published structures used in comparative analyses to AlphaFold3 models are: 4GNX (*U. maydis* RPA-ssDNA) [38], 7UY6 (*Tetrahymena* telomerase complex) [47], 6I52 (*S. cerevisiae* RPA-ssDNA) [91], 9SWN (*S. cerevisiae* telomerase complex) [54], and 7QXA (human telomerase-TPP1 complex) [77].

### Expression and purification of human telomerase

HEK293T cells (ATCC) were co-transfected with a modified pVan107-hTERT plasmid [92], in which TERT carried an N-terminal 3xFLAG-Twin-Strep tag, and pCDNA-U3-TR-HDV (a gift from Dr. Kelly Nguyen) at a 1:3 mass ratio using JetPrime reagent (Polyplus, France). Cells were harvested 48 hours after transfection and lysed in CHAPS lysis buffer (10 mM HEPES, pH 7.5, 1 mM MgCl2, 1 mM EGTA, 0.5% CHAPS, 10% glycerol, 5 mM TCEP, 1 mM PMSF). Lysates were clarified by centrifugation at 17,000×g for 20 min, and the supernatant was incubated with anti-DYKDDDDK G1 affinity resin (GenScript, USA) for 2 hours at 4°C with rotation. Beads were washed with 15 column volumes of telomerase buffer (50 mM Tris-HCl, pH 8.0, 50 mM KCl, 1 mM MgCl2, 1 mM TCEP, 30% glycerol), and bound material was eluted with 2 column volumes of the same buffer supplemented with 0.5 mg/mL 3xFLAG peptide (APExBIO, USA). Eluted telomerase was concentrated using Amicon Ultra 30 kDa MWCO centrifugal filters (Millipore), snap-frozen in liquid nitrogen in small aliquots, and stored at -80°C. TERT in purified telomerase was detected and quantified by Western blot. Samples were resolved by SDS-PAGE alongside a 3xFLAG-tagged protein standard, transferred to nitrocellulose using a semi-dry transfer module, and blocked with StartingBlock Blocking Buffer (Thermo Fisher Scientific, USA) for 1 hour at room temperature. Membranes were incubated overnight at 4°C with monoclonal anti-FLAG M2-HRP antibody (Sigma-Aldrich, A8592) at the manufacturer-recommended concentration, washed 3× with 1× PBST containing 0.5% Tween-20, and developed using SuperSignal West Pico PLUS chemiluminescent substrate (Thermo Fisher Scientific, USA). Human TR (hTR) in purified telomerase was detected and quantified by dot blot. Telomerase aliquots were incubated with Proteinase K and RNaseOUT (Invitrogen, USA) for 1 h at 37°C, mixed with an equal volume of formamide loading dye, heated to 95°C for 10 min, and spotted onto Hybond N+ membrane (Cytiva, USA) together with *in vitro*-transcribed hTR standards. Membranes were UV-crosslinked, pre-hybridized for 1 h at 42°C in ULTRAhyb Ultrasensitive Hybridization Buffer (Invitrogen, USA), and hybridized overnight at 42°C with ^32^P-labeled hTR DNA probes as described [26]. Membranes were washed twice each for 15 min at 42°C with Buffer W1 (2× SSC, 0.1% SDS) and Buffer W2 (0.1× SSC, 0.1% SDS), exposed to a phosphor screen, and imaged using a Typhoon RGB imager (Cytiva, USA). hTR quantification was used to determine the final concentration of purified telomerase.

### Expression and purification of human RPA

Human RPA and its single amino acid mutants (W107A, W528A, W528C, E240K) were purified using a previously described protocol [93, 94]. In summary, the plasmid p11d-tRPA (Catalog #102613, Addgene, USA) and its respective mutant plasmids were transformed into *E. coli* BL21(DE3) cells (New England Biolabs, USA). A single colony was picked and grown overnight in 4 L of LB medium at 37°C without shaking. They were grown the next day and induced at O.D. (600) of 0.6 by adding 0.3 mM IPTG. The cells were grown overnight at 16°C and collected the next day. The cells were resuspended in Buffer A (25 mM HEPES, 1 M NaCl, 10% Glycerol, 1 mM EDTA, 1 mM DTT, 0.5 mg/ml lysozyme, 1X Protease Inhibitor cocktail (Roche) and sonicated for lysis. The lysate was centrifuged at 35,000 x g for 45 min. The supernatant was buffer exchanged into a buffer containing 500 mM NaCl and incubated with 4 ml of HiTrap Blue beads for 1 hour at 4°C with constant rotation. It was then transferred to a gravity column and sequentially washed with 10 CV of Buffer A containing 500 mM NaCl, followed by 5 CV of Buffer A containing 800 mM NaCl. The RPA was eluted with 8 CV of Buffer A containing 2.5 M NaCl + 30% Ethylene Glycol. The eluate was diluted into a buffer containing 500 mM NaCl and incubated with 1 ml of hydrated ssDNA-cellulose (Cell Systems) for 1 hour at 4 °C with constant rotation. It was then transferred to a gravity column and subsequently washed with 10 CV Buffer A containing 500 mM NaCl. The RPA was eluted with 8 CV of Buffer A containing 1.5 M NaCl + 50% Ethylene Glycol. The elution was then diluted into a buffer containing 150 mM NaCl. The elute was subsequently loaded into a HiTrap Q column. The Q column was washed with wash buffers containing 150 mM NaCl. The protein was eluted using a gradient from 150 to 600 mM NaCl. SDS-PAGE was used to analyze protein bands, and the pure fractions were pooled and concentrated using Amicon Ultra 10 kDa MWCO centrifugal tubes (Millipore, USA) to obtain the final RPA protein complexes.

### Direct telomerase assay and quantification

A standard base reaction (20 μl) consisted of dNTPs, 200 nM 5’ Phos-TTAGGGTTAGGGTTAGGG DNA primer (Integrated DNA Technologies, USA), and approximately 0.5 nM telomerase (quantified using TR dot blot) in the assay buffer containing 50 mM HEPES–NaOH (pH 7.5), 100 mM KCl, 5 mM MgCl₂, and 2 mM DTT. Recombinant protein complexes and proteins were added to the reactions at concentrations specified in each figure legend. The reaction mixtures were pre-incubated at room temperature for 30 min, after which the enzymatic reactions were initiated by adding dNTPs. The dNTP mixture used was: 10 μM dGTP, 10 μM dTTP, 2.42 μM dATP, and 0.084 μM [α-³²P]dATP. The reactions were incubated at 37°C for 1 hour, then quenched with 5 M ammonium acetate and 200 μg/mL glycogen (Roche) for DNA precipitation. A radiolabeled 15-nt (TTAGGGTTAGGGTTA) oligonucleotide was added as a loading control in this step. After DNA precipitation, each sample was dissolved in 5 μl ultrapure water (Catalog #10977015, Invitrogen, USA) and 5 μl 2x formamide loading dye. 9 μL of each sample was loaded into each well of a 10% or 12% 1x TBE 7 M urea PAGE gel. Electrophoresis was performed at a constant 45 W until the bromophenol blue reached a third of the way up the gel. The gels were vacuum-dried at 80°C for 1 hour, then exposed to a storage phosphor screen. After exposure, the gels were imaged on an Amersham Typhoon gel imager (Cytiva, USA). Enzyme activity and repeat-addition processivity were quantified using GelAnalyzer software (version 23.11; www.gelanalyzer.com). Telomerase activity was determined by summing the signal intensities of all extension products and normalizing the sum to the loading control. Repeat addition processivity was calculated as the ratio of the cumulative signal from longer extension products (>+8 repeats) to that from shorter extension products (≤+8 repeats), as described previously [26].

## Supporting information

S1 Data: datasets for graphs

S2 Data: un-cropped images files

UCSF Chimera files for AF3 models of RPA-TERT-TPP1-like complexes

## Supporting information

**S1 Fig. Characterization of telomere maintenance defects in *ssb1* mutants isolated from a genetic screen. (A)** Schematic diagram of the fission yeast telomere region. For Southern blot analysis, genomic DNA was digested with EcoRI and hybridized with a telomere repeat probe. The EcoRI site lies ∼750 bp from the end of the telomeric repeat tract. Primers used in telomere ChIP qPCR (Table S4) are also indicated. **(B, C)** Telomere length analysis by Southern blot of the indicated *ssb1/rpa1* mutant strains. Mutant allele numbers are indicated (#), and relative Ssb1 protein expression levels, previously quantified by western blot [30], are shown. Mutants expressing less than 50% of wt Ssb1 are highlighted in red. **(D)** Plasmid-based complementation analysis: *ssb1-W523C* cells were transformed with plasmids carrying individual ssb1 point mutations and analyzed by Southern blot for telomere length. The original mutations addressed in this assay are indicated. Mutant plasmids that caused very short telomeres are highlighted in red. **(E)** Plasmid assay as in (D), using *ssb1* mutant plasmids carrying mutations equivalent to those known to cause telomere shortening in human cells, indicated by an asterisk (*).

**S2 Fig. Additional data for *ssb1* mutants with short telomeres. (A)** Schematic representation of the fission yeast *ssb1* locus, indicating the positions of the mutations. For integration selection purposes, the nmt1 3’ UTR and the kanamycin resistance marker kanMX were placed downstream of *ssb1*. **(B)** Telomere length analysis by Southern blot of the indicated *ssb1* mutant strains. All strains were extensively restreaked to obtain terminal telomere phenotypes. These strains represent independently generated second-mutant clones of the mutants shown in Fig 1C. **(C)** Expression levels of single *ssb1* mutant strains assessed by western blot analysis. Ssb1 was detected using an anti-Ssb1 antibody. A Ponceau-stained membrane is shown as loading control. Quantifications of relative Ssb1 protein expression levels to wt Ssb1 (set to 100%) are shown below. **(D)** Telomere length analysis by Southern blot of the indicated multiple independently obtained *ssb1* mutant strains. All strains were extensively restreaked to obtain terminal telomere phenotypes. **(E)** ChIP analysis of Trt1-myc binding to telomeres in the indicated *ssb1* mutant strains. Statistical significance was assessed by ANOVA with Dunnett’s comparison test (*P ≤ 0.05; **P ≤ 0.01). See S1 Data for individual % IP values and additional statistical analysis. **(F-H)** Telomere length analysis by Southern blot of the indicated *ssb1* mutants in *tpz1^+^* (wt) or *tpz1-K75A* backgrounds after extensive restreaks. Multiple independent *ssb1 tpz1-K75A* double-mutant strains were analyzed.

**S3 Fig. Sequence alignments. (A-C)** Sequence alignments with MUSCLE [95], generated using SnapGene for Ssb1/Rfa1/RPA1 (A), Ssb2/Rfa2/RPA2 (B), and Trt1/Est2/TERT (C). Residues discussed in this study are indicated. **(D)** Sequence alignment for the N-terminal domain of Tpz1/Est3/TPP1, adjusted based on AlphaFold3 structure models. TEL-patch residues, TELR residues, and their proposed interactions with other complex components within the RPA–TERT–TPP1-like complexes are indicated.

**S4 Fig. Further characterization of *ssb1-W523C* strains. (A)** Five-fold serial dilutions of the indicated strains were plated onto YES medium with the indicated concentrations of MMS, HU, or CPT. Pictures were taken after 3 days at 32 °C. Earlier-generation *ssb1-W523C* strains with linear chromosomes were significantly less sensitive to drugs than the *ssb1-W523C* strain that has lost telomeres. Control strains sensitive to drugs (*trt1Δ* with circular chromosomes, *rad3Δ*, and *ssb1-D223Y*) were also included. **(B)** Pictures of fluorescence Ssb3-GFP foci for the indicated strains overlayed with fission yeast cells. Scale bar represents 10 μm. **(C)** Quantification of the fraction of cells with Ssb3-GFP foci for the indicated strains. Statistical significance was assessed by ANOVA with Dunnett’s comparison test (**P ≤ 0.01; ***P ≤ 0.001; ****P ≤ 0.0001). See S1 Data for individual % nuclei with foci values and additional statistical analysis.

**S5 Fig. Strategies for generating elongated telomere strain series. (A)** Schematic diagram of the strategy used to generate elongated isogenic telomere series. For more details, see the Materials and Methods section. **(B)** Representative Southern blot analysis of a telomere-elongated strain series for *trt1-D743C-myc.* **(C-E)** Southern blots for parental strains used in *rap1Δ* x *poz1Δ* crosses to generate telomere-elongated strains.

**S6 Fig. AlphaFold3 prediction of the RPA–Trt1–Tpz1 complex. (A, B)** Overlay of top-scoring AlphaFold3 models for the RPA–Trt1–Tpz1 complex, shown from two different viewpoints. (A) Overlay of models containing full-length RPA subunits (Ssb1, Ssb2, and Ssb3), full-length Trt1, and Tpz1(aa1-219). (B) Overlay of models containing trimmed RPA subunits (Ssb1 and Ssb2), full-length Ssb3 and Trt1, and Tpz1(aa1-155). **(C)** Superposition of the fission yeast AlphaFold3 model (left) with the *Tetrahymena* telomerase cryo-EM structure (center), shown individually and overlaid (right) in two orientations. **(D)** Detailed model of the interaction between Tpz1 TEL-patch residues K75 and E74 and Trt1-TEN-domain residues E143 and K145, respectively. Residues Tpz1-E116 and Trt1-S85, which may further stabilize this interface, are also indicated. **(E)** Detailed structure of the Ssb1–Trt1 interface.

**S7 Fig. Analysis of *ssb2* and *ssb3* mutants, and comparison between the RPA structure within the AlphaFold3 RPA–Trt1–Tpz1 model and the X-ray structure of the RPA–ssDNA complex. (A)** Schematic representations of fission yeast Ssb2 truncation mutants lacking the WH domain and of Ssb3. Regions represented in the *U. maydis* X-ray structure are indicated. **(B, C)** Telomere length analysis by Southern blot of *ssb2* truncation mutants (B) and *ssb3Δ* cells (C). **(D)** X-ray structure of *U. maydis* RPA bound to ssDNA [38], labeled using fission yeast nomenclature. DBD-A, -B, and -C of Ssb1, DBD-D of Ssb2, and OB-E of Ssb3 are indicated. Residues discussed in this study are marked. **(E, F)** Corresponding structures from the fission yeast AlphaFold3 model and *U. maydis* X-ray structure are shown individually and superimposed.

**S8 Fig. Additional data related to *ssb2* and *trt1* mutants. (A)** Telomere length analysis by Southern blot of *tpz1-R81E ssb2* double mutants in the presence (*trt1^+^*) or absence (*trt1Δ*) of telomerase. Terminal telomere phenotypes were determined after extensive restreaking. **(B, C)** Telomere length analysis by Southern blot of the indicated *ssb2* mutants affecting the Ssb2–Tpz1 (B) or the Ssb2–Trt1 (C) interfaces, each in combination with *tpz1-K75A*. Terminal telomere phenotypes were determined after extensive restreaking. **(D)** Western blot analysis of mutant *ssb2-FLAG* strains to verify expression levels. Ssb2 was detected with an anti-FLAG antibody, and tubulin served as a loading control. **(E)** ChIP analysis of Trt1-myc binding to telomeres in the indicated *ssb2* mutant strains, with corresponding telomere length analysis of the strains used for ChIP. Statistical significance was assessed by ANOVA with Dunnett’s comparison test (*P ≤ 0.05). See S1 Data for individual % IP values and additional statistical analysis. **(F)** Telomere length analysis by Southern blot of the indicated *trt1* mutant strains after extensive restreaking. Two independently derived clones were analyzed for each mutant. **(G)** Western blot analysis for mutant *trt1-myc* strains to verify expression levels. Trt1 was detected with an anti-myc antibody, and tubulin served as a loading control. **(H)** Telomere length analysis by Southern blot of the indicated *trt1* mutants in *tpz1^+^* (wt) or *tpz1-K75A* backgrounds. Two independent *tpz1-K75A* double-mutant strains were analyzed. Terminal telomere phenotypes were determined after extensive restreaking.

**S9 Fig. Suppressor analysis for *ssb1-W523C* and *tpz1-R81E*. (A-G)** Telomere length analysis by Southern blot of the indicated fission yeast mutant strains.

**S10 Fig. Analysis of DNA damage sensitivity for *rfa1-W533C* strains.** Ten-fold serial dilutions of the indicated strains were plated onto YPDA medium with the indicated concentrations of MMS, HU, or CPT. Damage-sensitive *rad52Δ* strain served as a control. Pictures were taken after 2 days at 30 °C.

**S11 Fig. AlphaFold3 prediction of the RPA–Est2–Est3 complex. (A)** Overlay of top-scoring AlphaFold3 models for the budding yeast RPA–Est2–Est3 complex, shown from two different viewpoints. **(B)** Superposition of the fission yeast RPA–Trt1–Tpz1 complex with the budding yeast RPA-Est2-Est3 complex. **(C)** Superposition of the *Tetrahymena* Cryo-EM structure with the budding yeast RPA–Est2–Est3 complex. **(D)** Superposition of the budding yeast X-ray structure RPA–ssDNA with the budding yeast RPA–Est2–Est3 complex. **(E)** Superposition of the budding yeast cryo-EM structure of telomerase complex with the budding yeast RPA–Est2–Est3 complex. **(F)** Detailed structures of the Rfa1–Est2 interface. Five top-scoring AlphaFold3 models are shown individually (light frames) or superimposed (bold frame).

**S12 Fig. AlphaFold3 prediction of the RPA–TERT–TPP1 complex. (A)** Overlay of the top-scoring AlphaFold3 models for the human RPA–TERT–TPP1 complex, shown from two different viewpoints. **(B, C)** Human RPA–TERT–TPP1 model from AlphaFold3, and Human TERT–TPP1 cryo-EM structure, and RPA–ssDNA X-ray structure from *U.* maydis, shown individually (B) or superimposed (C). **(D)** Detailed view of the predicted human D61^RPA2^–R220^TPP1^ interaction, supported by neighboring residues, together with K78^TERT^–E215^TTP1^ interactions. **(E)** Detailed view of the predicted TERT-G106/A107 interaction with RPA1-W528, stabilized by intramolecular TERT interactions involving E113, R132, and K78, as well as the TPP1-E215 interaction. TPP1 is omitted from this panel to help visualize the relevant interactions.

**S13 Fig. Comparison to RPA–TERT–TPP1-like complex models from budding yeast and humans. (A, B)** Functionally important protein-protein interactions identified in the fission yeast RPA–Trt1–Tpz1 complex are conserved in budding yeast (A) and humans (B). Rfa1–Est2/RPA1–TERT interactions are shown in pink, Rfa2–Est2/RPA2–TERT in yellow, Rfa2–Est3/RPA2–TPP1 in blue, and Est2–Est3/TERT–TPP1 in green. Interacting sites are indicated. **(C)** Sequence alignments for Ssb2/Rfa2/RPA2 (left) and Tpz1/Est3/TPP1 (right). Interactions of individual residues with other complex partners are indicated.

**S14 Fig. Outputs from the AlphaFold3 server. (A)** *S. pombe* Trt1-RPA-Tpz1. **(B)** *S. cerevisiae* Est2-RPA-Est3. **(C)** *H. sapiens* TERT-RPA-TPP1.

**S1 Table. Fission yeast strains used in this study.**

**S2 Table. Budding yeast strains used in this study.**

**S3 Table. Plasmids used in this study.**

**S4 Table. DNA primers used in this study.**

**S5 Table. Source of various mutated and tagged alleles for fission yeast strains used in this study.**

**S1 Data. Source data and statistical analysis for graphs in main and supplemental figures.**

**S2 Data. Uncropped images for main and supplemental figures.**

**S3 Data. UCSF Chimera X files for RPA-TERT-TPP1-like complexes from *S. pombe*, *S. cerevisiae*, and humans.**

## Acknowledgements

This work was supported by NIH/National Institute of General Medical Sciences grants R01GM143316 to TMN, R01GM153806 to CJL, and R35GM144307 to YjX. We thank Drs. Paul Russell (The Scripps Research Institute), Tamás Fischer (Australian National University), Virginia A. Zakian (Princeton University), Julia P. Cooper (University of Colorado Anschutz), Katherine L. Friedman (Vanderbilt University), Fuyuki Ishikawa (Kyoto University), and Junko Kanoh (University of Tokyo) for sharing published fission yeast and budding yeast strains, Kelly Nguyen (MRC Laboratory of Molecular Biology) for the pCDNA-U3-TR-HDV plasmid, and Bianca Linde and Dayo Akinsiku for their initial efforts in generating reagents used in this study. We also thank Yaneris Alvarado-Cartagena (University of Illinois Chicago) for help with fluorescence microscopy. While the manuscript is entirely written and edited by the authors, we consulted Grammarly and ChatGPT to improve its grammar and readability. The authors take full responsibility for all the contents of the manuscript.

