## Supplemental Figures S1-S14 for "Fission yeast RPA–TERT–Tpz1^TPP1^ complex promotes telomere extension and suppresses telomere recombination"

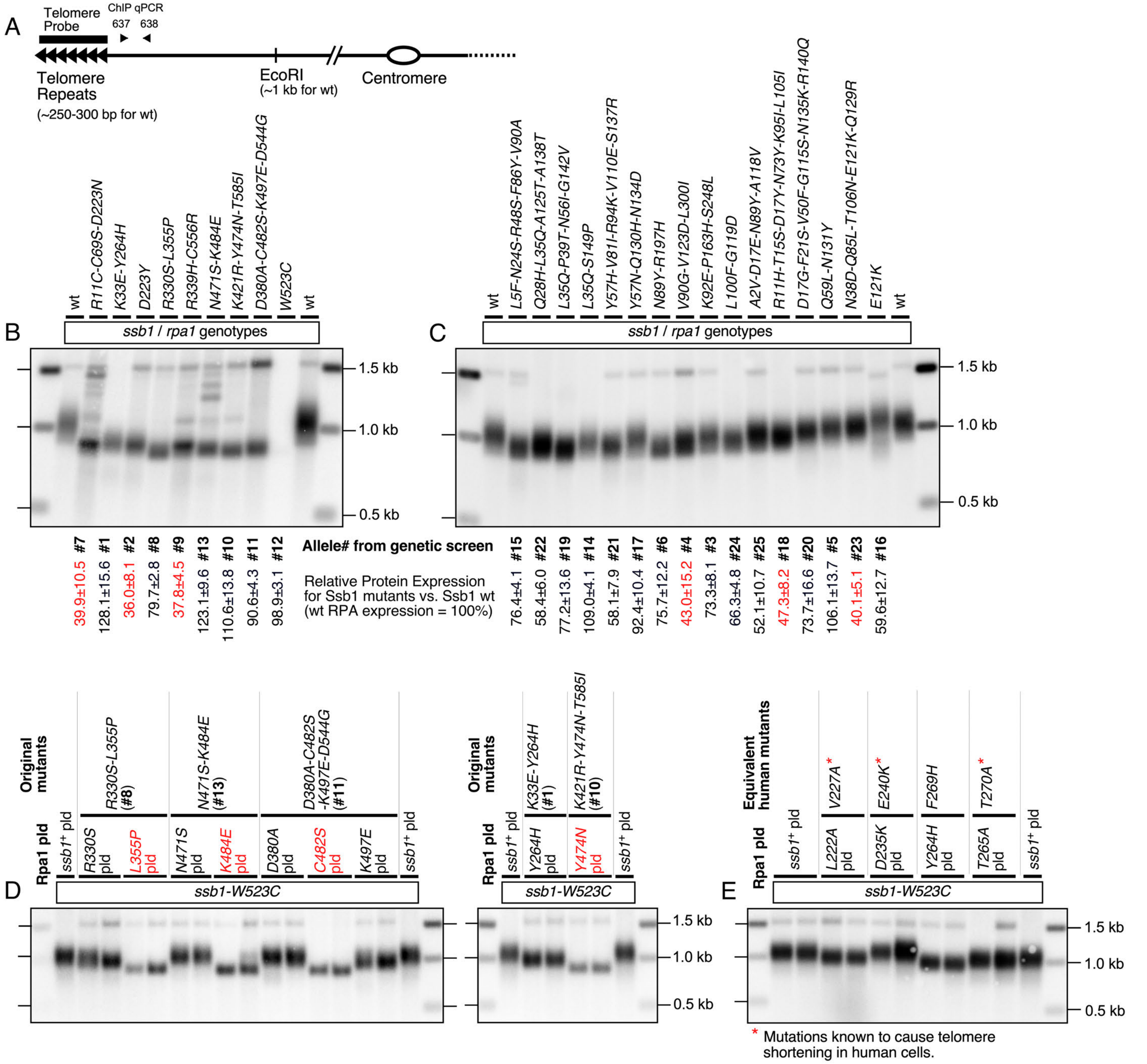


**S1 Fig. Characterization of telomere maintenance defects in *ssb1* mutants isolated from a genetic screen. (A)** Schematic diagram of the fission yeast telomere region. For Southern blot analysis, genomic DNA was digested with EcoRI and hybridized with a telomere repeat probe. The EcoRI site lies ~750 bp from the end of the telomeric repeat tract. Primers used in telomere ChIP qPCR (Table S4) are also indicated. **(B, C)** Telomere length analysis by Southern blot of the indicated *ssb1/rpa1* mutant strains. Mutant allele numbers are indicated (#), and relative Ssb1 protein expression levels, previously quantified by western blot [30], are shown. Mutants expressing less than 50% of wt Ssb1 are highlighted in red. **(D)** Plasmid-based complementation analysis: *ssb1-W523C* cells were transformed with plasmids carrying individual ssb1 point mutations and analyzed by Southern blot for telomere length. The original mutations addressed in this assay are indicated. Mutant plasmids that caused very short telomeres are highlighted in red. **(E)** Plasmid assay as in (D), using *ssb1* mutant plasmids carrying mutations equivalent to those known to cause telomere shortening in human cells, indicated by an asterisk (*).

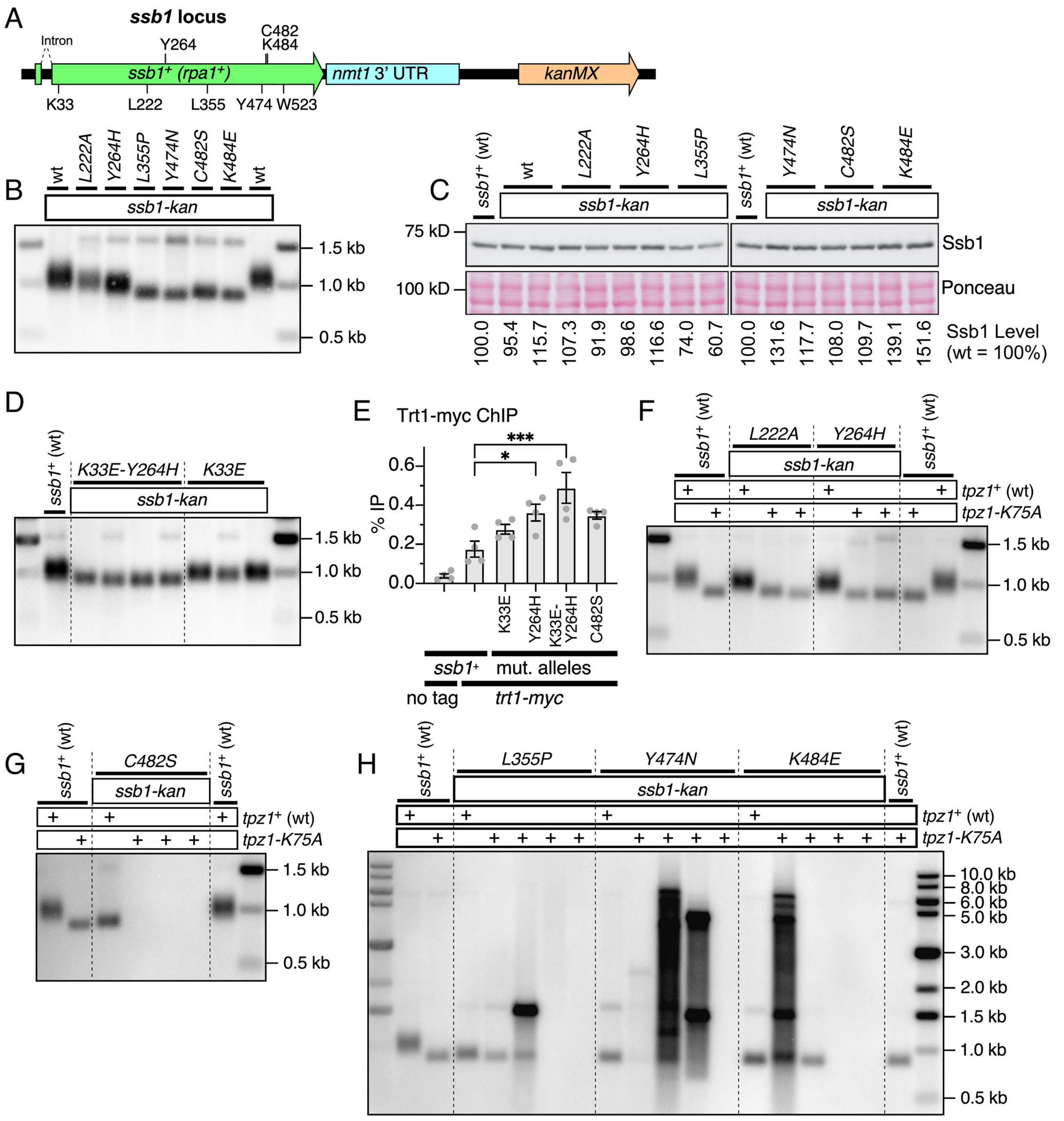
A
ssb1 locus
Intron
Y264
C482
K484
ssb1+ (rpa1+)
nmt1 3' UTR
kanMX
K33
L222
L355
Y474
W523
B
wt
L222A
Y264H
L355P
Y474N
C482S
K484E
wt
ssb1-kan
1.5 kb
1.0 kb
0.5 kb
C
ssb1+ (wt)
wt
L222A
Y264H
L355P
ssb1+ (wt)
Y474N
C482S
K484E
ssb1-kan
ssb1-kan
75 kD
100 kD
Ssb1
Ponceau
100.0 95.4 115.7 107.3 91.9 98.6 116.6 74.0 60.7 100.0 131.6 117.7 108.0 109.7 139.1 151.6
Ssb1 Level
(wt = 100%)
D
ssb1+ (wt)
K33E-Y264H
K33E
ssb1-kan
1.5 kb
1.0 kb
0.5 kb
E
Trt1-myc ChIP
% IP
0.6
0.4
0.2
0.0
***
*
K33E
Y264H
K33E-Y264H
C482S
ssb1+
mut. alleles
no tag
trt1-myc
F
ssb1+ (wt)
L222A
Y264H
ssb1+ (wt)
ssb1-kan
tpz1+ (wt)
tpz1-K75A
1.5 kb
1.0 kb
0.5 kb
G
ssb1+ (wt)
C482S
ssb1-kan
ssb1+ (wt)
tpz1+ (wt)
tpz1-K75A
1.5 kb
1.0 kb
0.5 kb
H
ssb1+ (wt)
L355P
Y474N
K484E
ssb1-kan
ssb1+ (wt)
tpz1+ (wt)
tpz1-K75A
10.0 kb
8.0 kb
6.0 kb
5.0 kb
3.0 kb
2.0 kb
1.5 kb
1.0 kb
0.5 kb

**S2 Fig. Additional data for *ssb1* mutants with short telomeres. (A)** Schematic representation of the fission yeast *ssb1* locus, indicating the positions of the mutations. For integration selection purposes, the nmt1 3' UTR and the kanamycin resistance marker kanMX were placed downstream of *ssb1*. **(B)** Telomere length analysis by Southern blot of the indicated *ssb1* mutant strains. All strains were extensively restreaked to obtain terminal telomere phenotypes. These strains represent independently generated second-mutant clones of the mutants shown in Fig 1C. **(C)** Expression levels of single *ssb1* mutant strains assessed by western blot analysis. Ssb1 was detected using an anti-Ssb1 antibody. A Ponceau-stained membrane is shown as loading control. Quantifications of relative Ssb1 protein expression levels to wt Ssb1 (set to 100%) are shown below. **(D)** Telomere length analysis by Southern blot of the indicated multiple independently obtained *ssb1* mutant strains. All strains were extensively restreaked to obtain terminal telomere phenotypes. **(E)** ChIP analysis of Trt1-myc binding to telomeres in the indicated *ssb1* mutant strains. Statistical significance was assessed by ANOVA with Dunnett's comparison test (*$P \leq 0.05$; ***$P \leq 0.001$). See S1 Data for individual % IP values and additional statistical analysis. **(F-H)** Telomere length analysis by Southern blot of the indicated *ssb1* mutants in *tpz1*$^{+}$ (wt) or *tpz1-K75A* backgrounds after extensive restreaks. Multiple independent *ssb1 tpz1-K75A* double-mutant strains were analyzed.

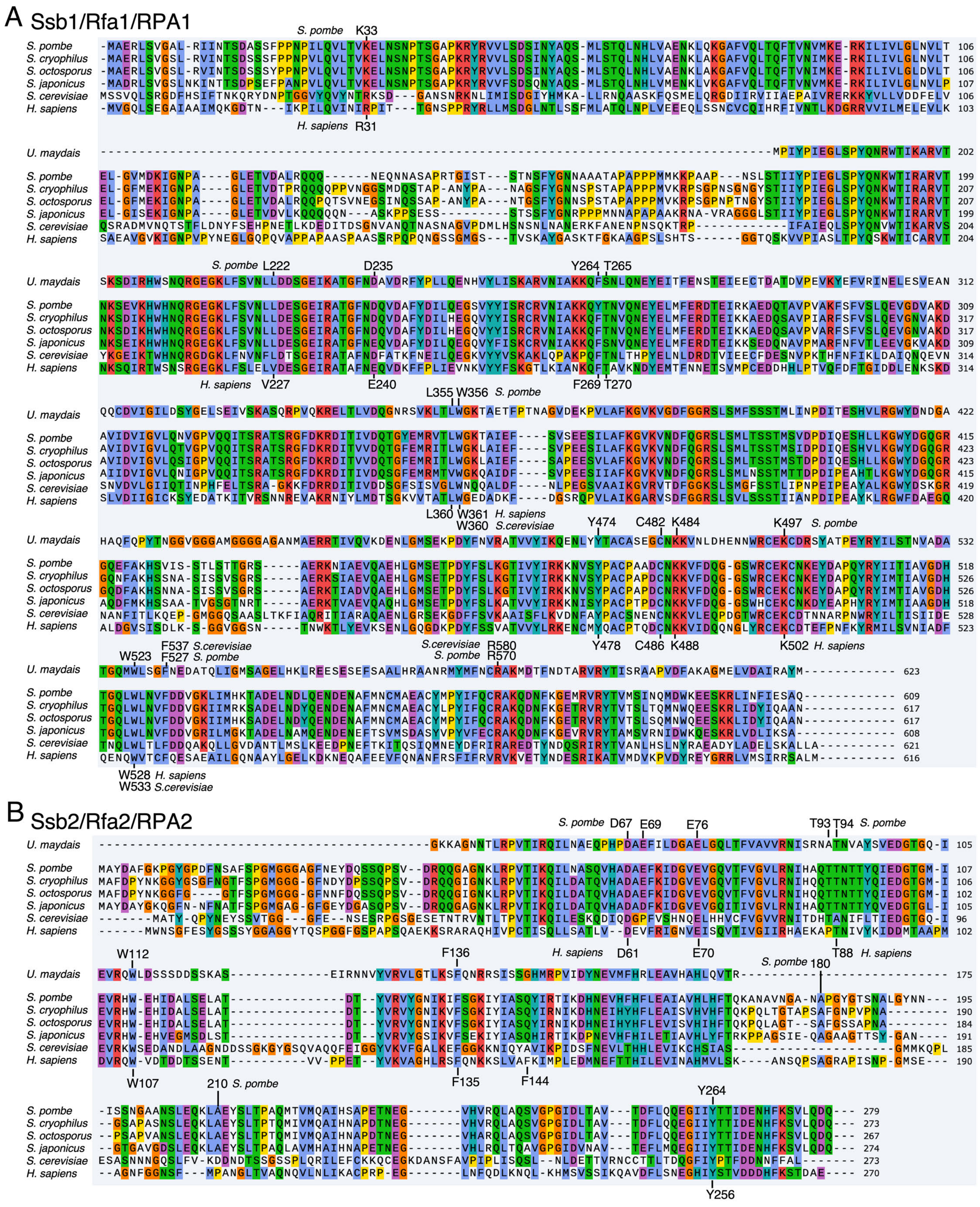
A Ssb1/Rfa1/RPA1
B Ssb2/Rfa2/RPA2
U. maydais
S. pombe
S. cryophilus
S. octosporus
S. japonicus
S. cerevisiae
H. sapiens
S. pombe K33
H. sapiens R31
S. pombe L222
D235
Y264 T265
H. sapiens V227
E240
F269 T270
L355 W356 S. pombe
L360 W361 H. sapiens
W360 S.cerevisiae
Y474
C482 K484
K497 S. pombe
Y478
C486 K488
K502 H. sapiens
W523
F537 S.cerevisiae
F527 S. pombe
S.cerevisiae R580
S. pombe R570
W528 H. sapiens
W533 S.cerevisiae
S. pombe D67
E69
E76
T93 T94 S. pombe
H. sapiens D61
E70
T88 H. sapiens
W112
F136
S. pombe 180
W107
F135
F144
210 S. pombe
Y264
Y256
106
106
106
107
106
103
202
199
207
207
199
204
204
312
309
317
317
309
314
314
422
415
423
423
415
419
420
532
518
526
526
518
528
523
623
609
617
617
608
621
616
105
107
106
102
105
96
102
175
195
190
184
191
185
190
279
273
267
274
273
270

**C** Trt1/Est2/TERT

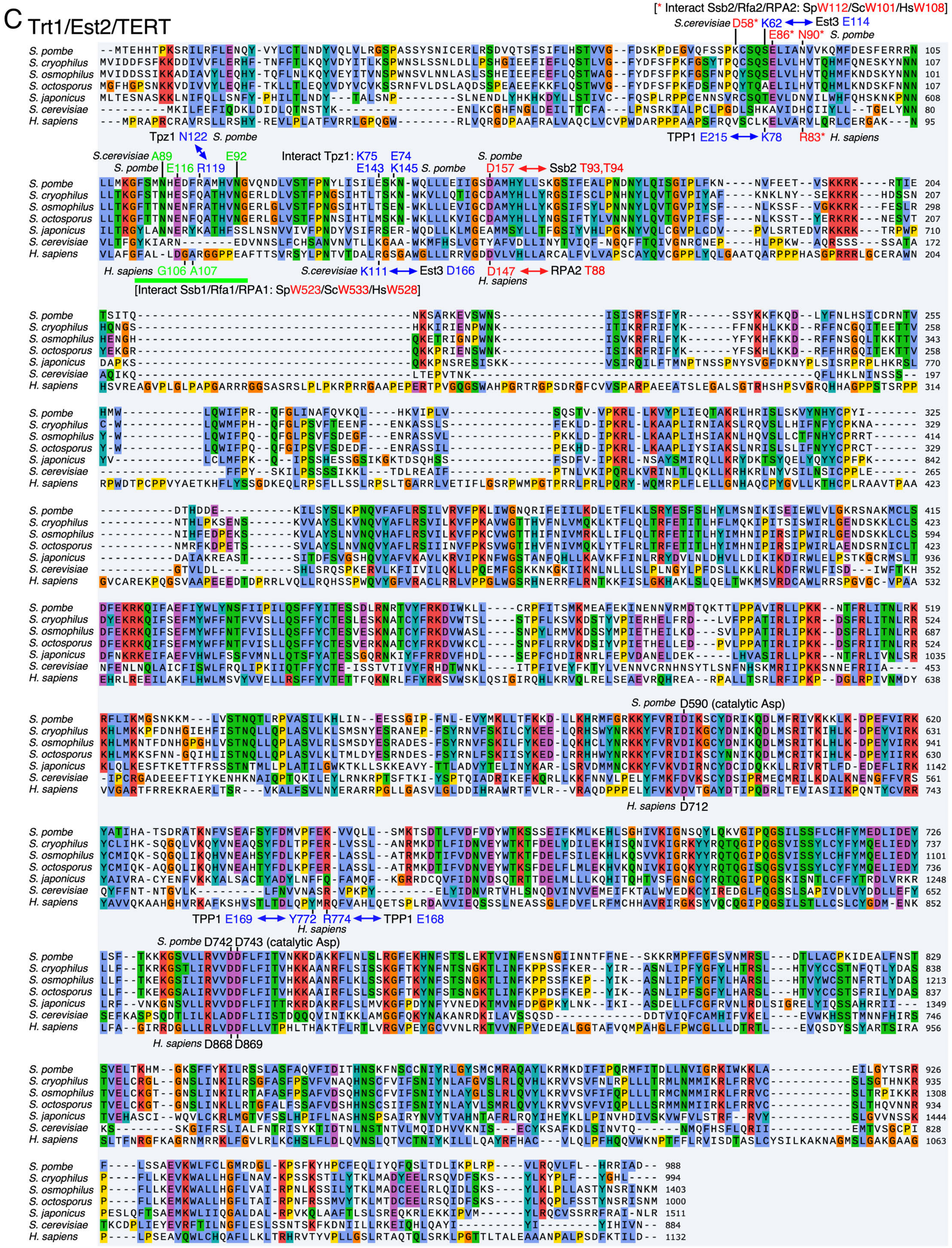

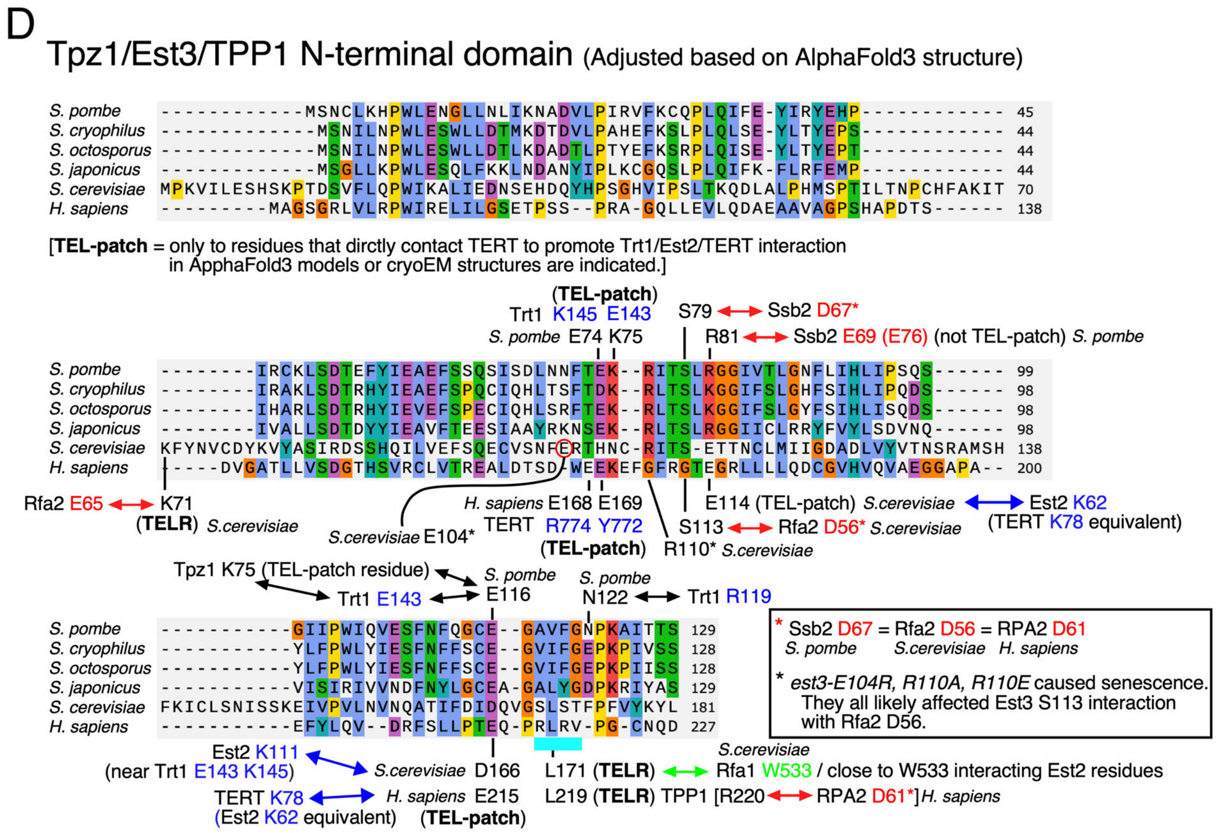


**S3 Fig. Sequence alignments. (A-C)** Sequence alignments with MUSCLE [95], generated using SnapGene for Ssb1/Rfa1/RPA1 (A), Ssb2/Rfa2/RPA2 (B), and Trt1/Est2/TERT (C). Residues discussed in this study are indicated. **(D)** Sequence alignment for the N-terminal domain of Tpz1/Est3/TPP1, adjusted based on AlphaFold3 structure models. TEL-patch residues, TELR residues, and their proposed interactions with other complex components within the RPA–TERT–TPP1-like complexes are indicated.

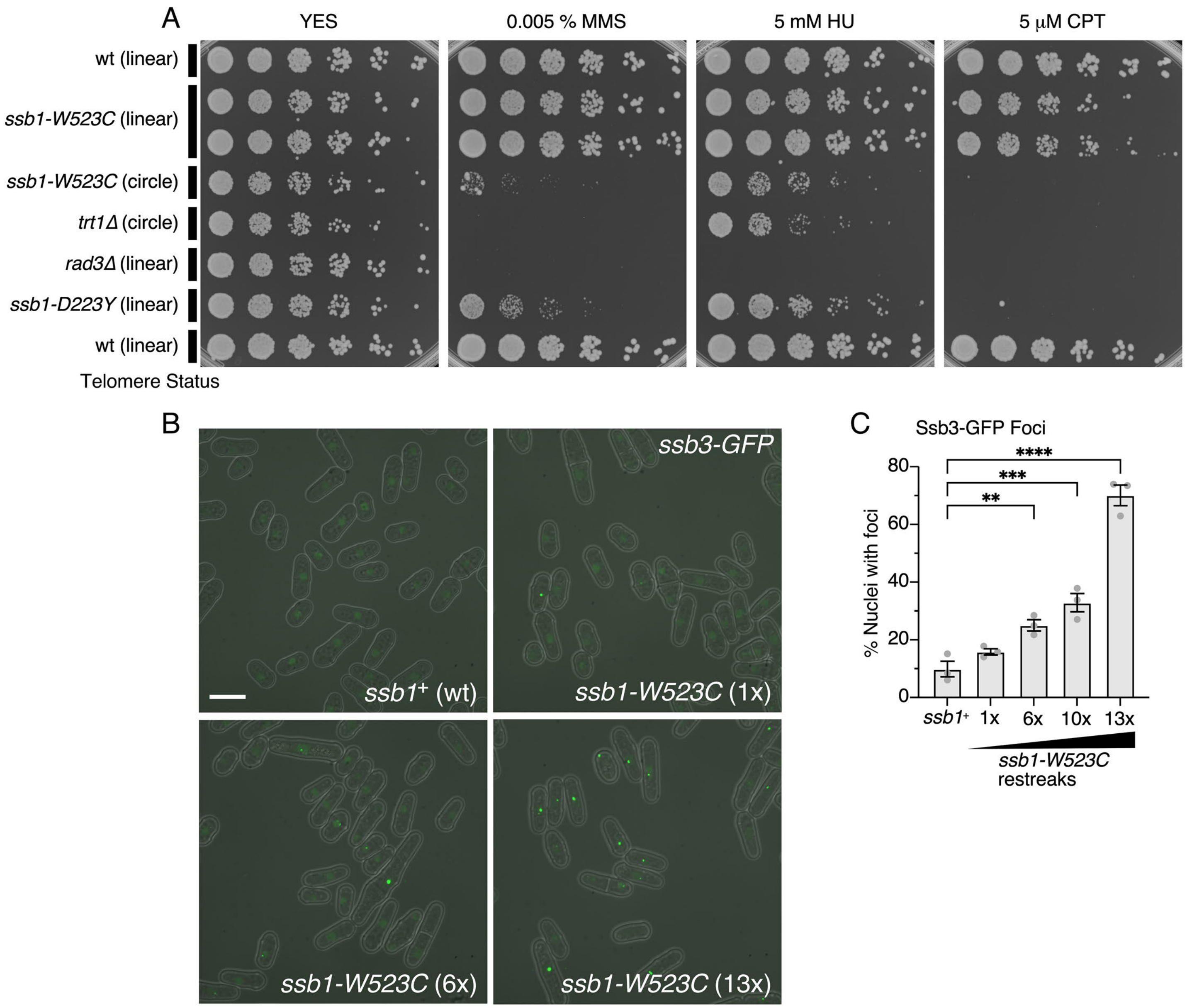


**S4 Fig. Further characterization of *ssb1-W523C* strains. (A)** Five-fold serial dilutions of the indicated strains were plated onto YES medium with the indicated concentrations of MMS, HU, or CPT. Pictures were taken after 3 days at 32 °C. Earlier-generation *ssb1-W523C* strains with linear chromosomes were significantly less sensitive to drugs than the *ssb1-W523C* strain that has lost telomeres. Control strains sensitive to drugs (*trt1Δ* with circular chromosomes, *rad3Δ*, and *ssb1-D223Y*) were also included. **(B)** Pictures of fluorescence Ssb3-GFP foci for the indicated strains overlayed with fission yeast cells. Scale bar represents 10 μm. **(C)** Quantification of the fraction of cells with Ssb3-GFP foci for the indicated strains. Statistical significance was assessed by ANOVA with Dunnett's comparison test (**P ≤ 0.01; ***P ≤ 0.001; ****P ≤ 0.0001). See S1 Data for individual % nuclei with foci values and additional statistical analysis.

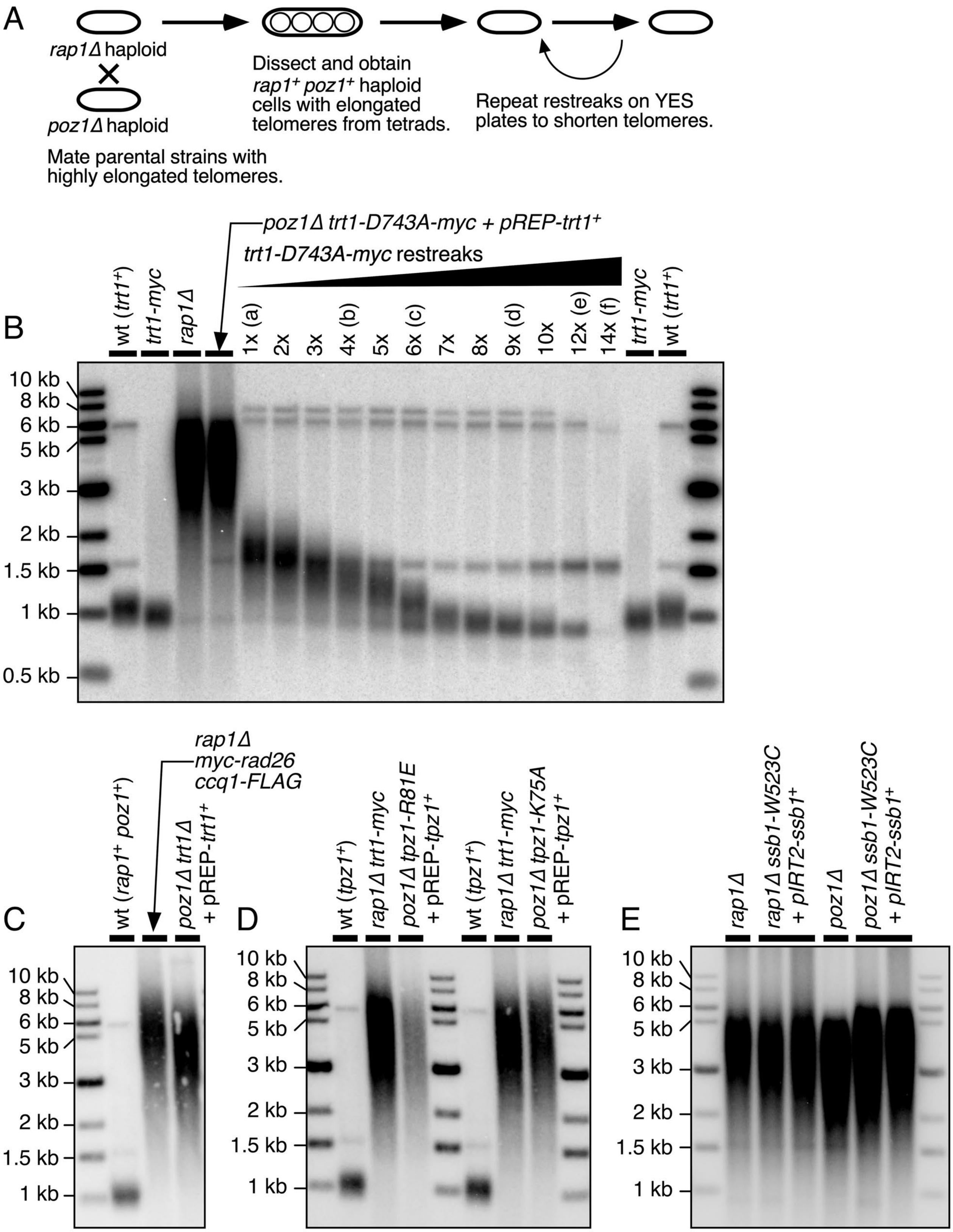


**S5 Fig. Strategies for generating elongated telomere strain series. (A)** Schematic diagram of the strategy used to generate elongated isogenic telomere series. For more details, see the Materials and Methods section. **(B)** Representative Southern blot analysis of a telomere-elongated strain series for *trt1-D743C-myc.* **(C-E)** Southern blots for parental strains used in *rap1Δ* x *poz1Δ* crosses to generate telomere-elongated strains.

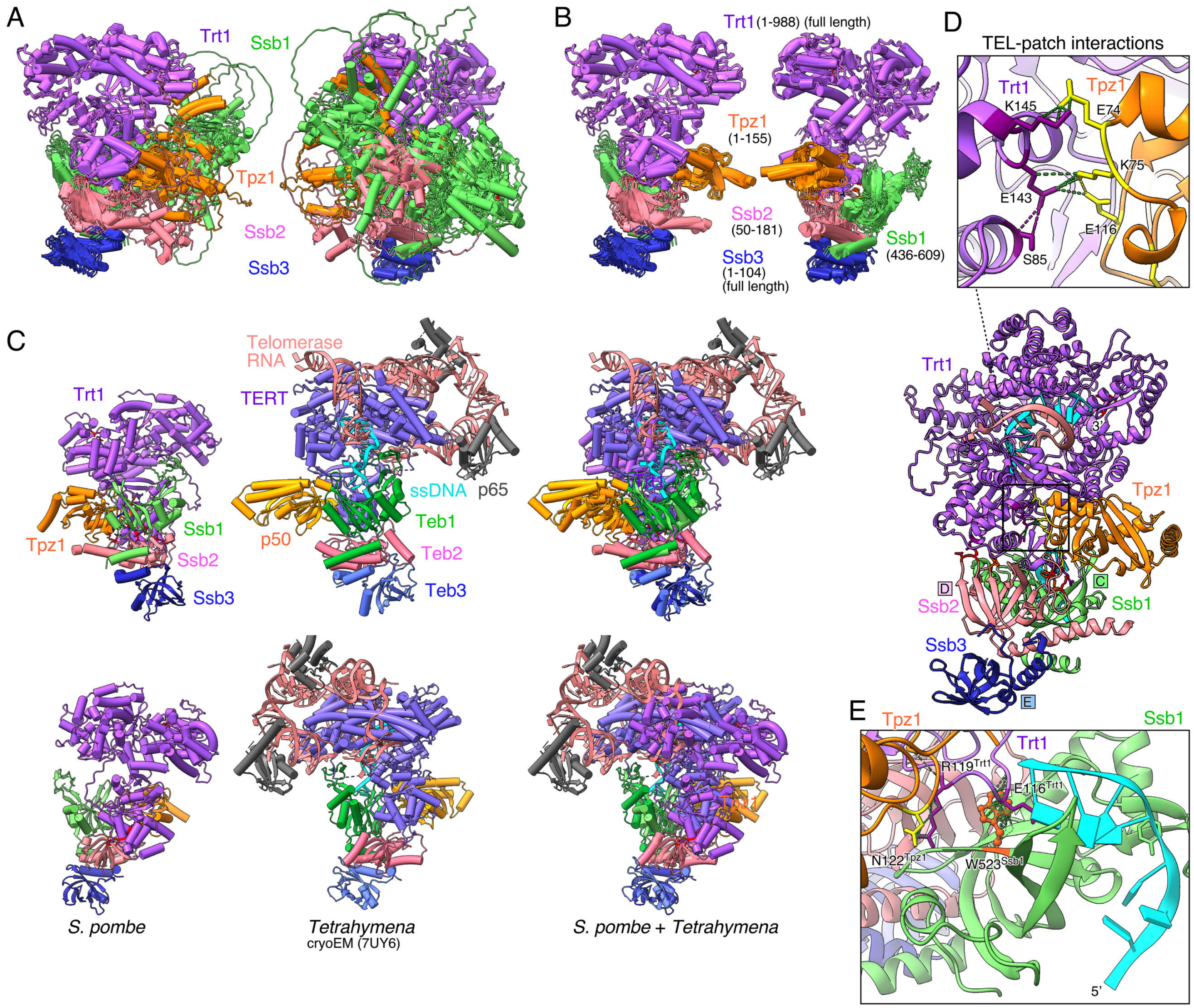


**S6 Fig. AlphaFold3 prediction of the RPA–Trt1–Tpz1 complex. (A, B)** Overlay of top-scoring AlphaFold3 models for the RPA–Trt1–Tpz1 complex, shown from two different viewpoints. (A) Overlay of models containing full-length RPA subunits (Ssb1, Ssb2, and Ssb3), full-length Trt1, and Tpz1(aa1-219). (B) Overlay of models containing trimmed RPA subunits (Ssb1 and Ssb2), full-length Ssb3 and Trt1, and Tpz1(aa1-155). **(C)** Superposition of the fission yeast AlphaFold3 model (left) with the *Tetrahymena* telomerase cryo-EM structure (center), shown individually and overlaid (right) in two orientations. **(D)** Detailed model of the interaction between Tpz1 TEL-patch residues K75 and E74 and Trt1-TEN-domain residues E143 and K145, respectively. Residues Tpz1-E116 and Trt1-S85, which may further stabilize this interface, are also indicated. **(E)** Detailed structure of the Ssb1–Trt1 interface.

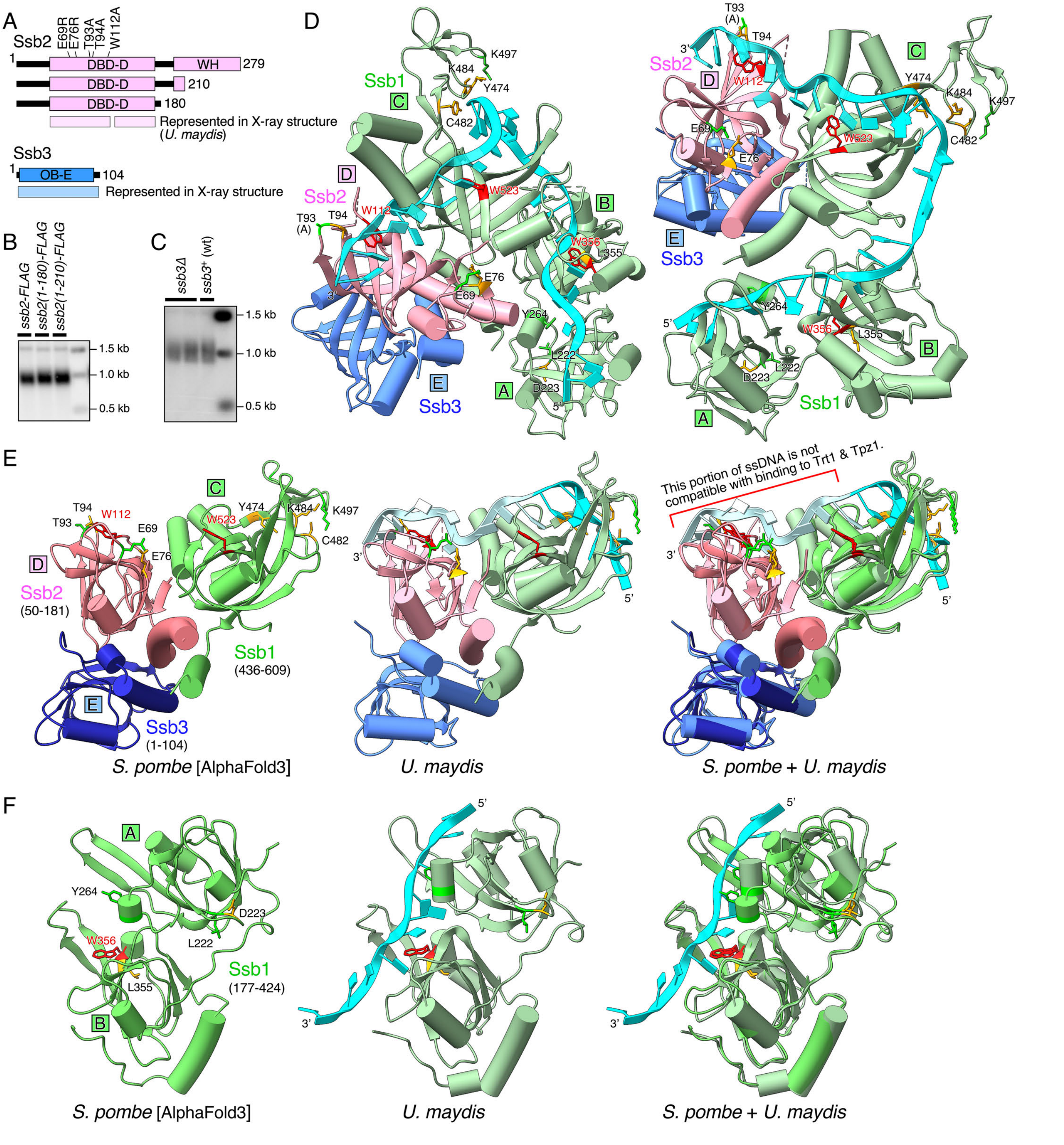


**S7 Fig. Analysis of *ssb2* and *ssb3* mutants, and comparison between the RPA structure within the AlphaFold3 RPA–Trt1–Tpz1 model and the X-ray structure of the RPA–ssDNA complex. (A)** Schematic representations of fission yeast Ssb2 truncation mutants lacking the WH domain and of Ssb3. Regions represented in the *U. maydis* X-ray structure are indicated. **(B, C)** Telomere length analysis by Southern blot of *ssb2* truncation mutants (B) and *ssb3Δ* cells (C). **(D)** X-ray structure of *U. maydis* RPA bound to ssDNA [38], labeled using fission yeast nomenclature. DBD-A, -B, and -C of Ssb1, DBD-D of Ssb2, and OB-E of Ssb3 are indicated. Residues discussed in this study are marked. **(E, F)** Corresponding structures from the fission yeast AlphaFold3 model and *U. maydis* X-ray structure are shown individually and superimposed.

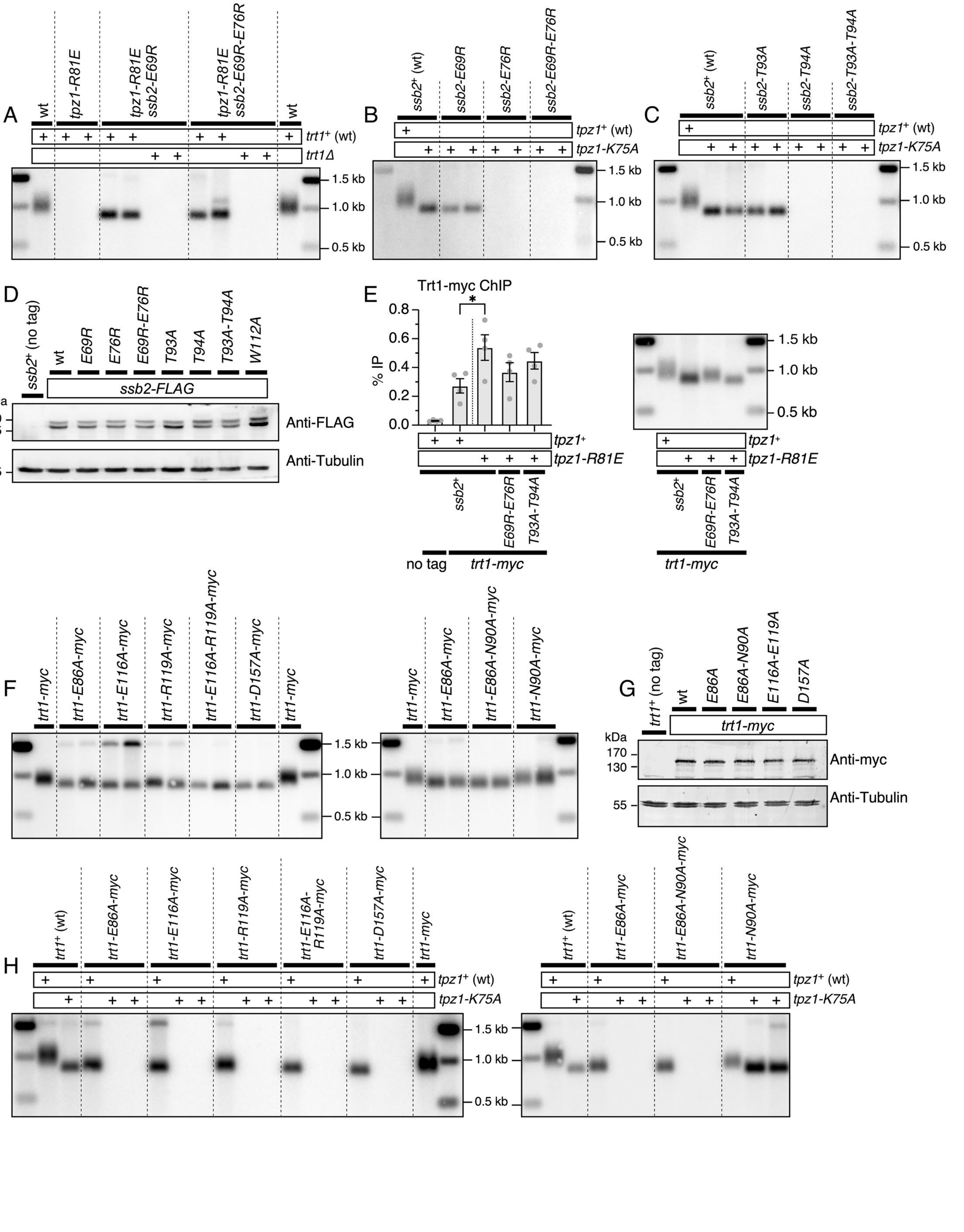
A
wt
tpz1-R81E
tpz1-R81E ssb2-E69R
tpz1-R81E ssb2-E69R-E76R
trt1+ (wt)
trt1Δ
1.5 kb
1.0 kb
0.5 kb
B
ssb2+ (wt)
ssb2-E69R
ssb2-E76R
ssb2-E69R-E76R
tpz1+ (wt)
tpz1-K75A
C
ssb2-T93A
ssb2-T94A
ssb2-T93A-T94A
D
ssb2+ (no tag)
wt
E69R
E76R
E69R-E76R
T93A
T94A
T93A-T94A
W112A
ssb2-FLAG
kDa
40
35
55
Anti-FLAG
Anti-Tubulin
E
Trt1-myc ChIP
*
% IP
0.8
0.6
0.4
0.2
0.0
tpz1+
tpz1-R81E
ssb2+
E69R-E76R
T93A-T94A
no tag
trt1-myc
F
trt1-myc
trt1-E86A-myc
trt1-E116A-myc
trt1-R119A-myc
trt1-E116A-R119A-myc
trt1-D157A-myc
trt1-E86A-N90A-myc
trt1-N90A-myc
G
trt1+ (no tag)
E86A
E86A-N90A
E116A-E119A
D157A
170
130
Anti-myc
H
trt1+ (wt)
trt1-E116A-R119A-myc

**S8 Fig. Additional data related to *ssb2* and *trt1* mutants. (A)** Telomere length analysis by Southern blot of *tpz1-R81E ssb2* double mutants in the presence (*trt1*$^{+}$) or absence (*trt1Δ*) of telomerase. Terminal telomere phenotypes were determined after extensive restreaking. **(B, C)** Telomere length analysis by Southern blot of the indicated *ssb2* mutants affecting the Ssb2–Tpz1 (B) or the Ssb2–Trt1 (C) interfaces, each in combination with *tpz1-K75A*. Terminal telomere phenotypes were determined after extensive restreaking. **(D)** Western blot analysis of mutant *ssb2-FLAG* strains to verify expression levels. Ssb2 was detected with an anti-FLAG antibody, and tubulin served as a loading control. **(E)** ChIP analysis of Trt1-myc binding to telomeres in the indicated *ssb2* mutant strains, with corresponding telomere length analysis of the strains used for ChIP. Statistical significance was assessed by ANOVA with Dunnett's comparison test (*$P \leq 0.05$). See S1 Data for individual % IP values and additional statistical analysis. **(F)** Telomere length analysis by Southern blot of the indicated *trt1* mutant strains after extensive restreaking. Two independently derived clones were analyzed for each mutant. **(G)** Western blot analysis for mutant *trt1-myc* strains to verify expression levels. Trt1 was detected with an anti-myc antibody, and tubulin served as a loading control. **(H)** Telomere length analysis by Southern blot of the indicated *trt1* mutants in *tpz1*$^{+}$ (wt) or *tpz1-K75A* backgrounds. Two independent *tpz1-K75A* double-mutant strains were analyzed. Terminal telomere phenotypes were determined after extensive restreaking.

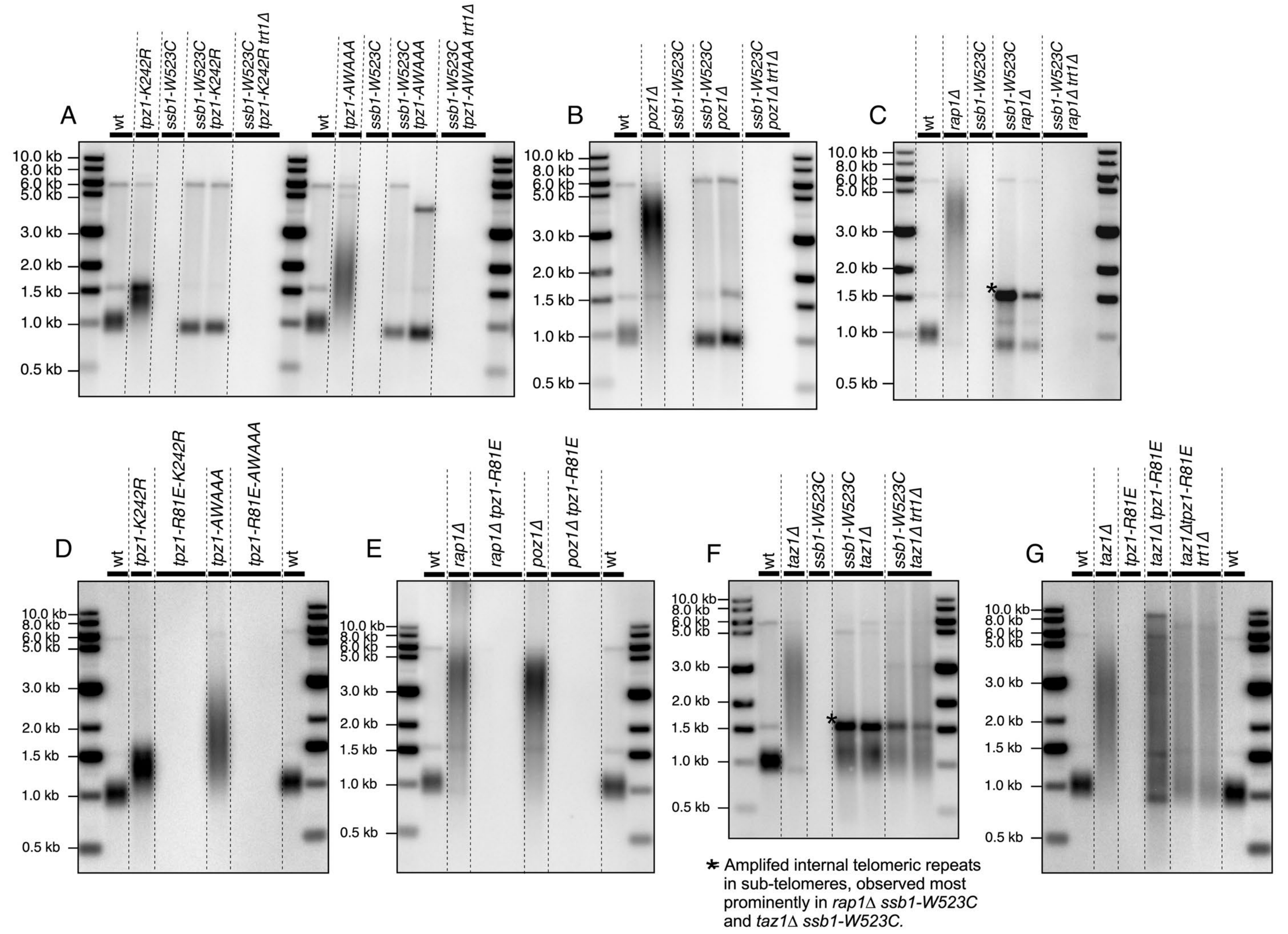


**S9 Fig. Suppressor analysis for *ssb1-W523C* and *tpz1-R81E*. (A-G)** Telomere length analysis by Southern blot of the indicated fission yeast mutant strains.

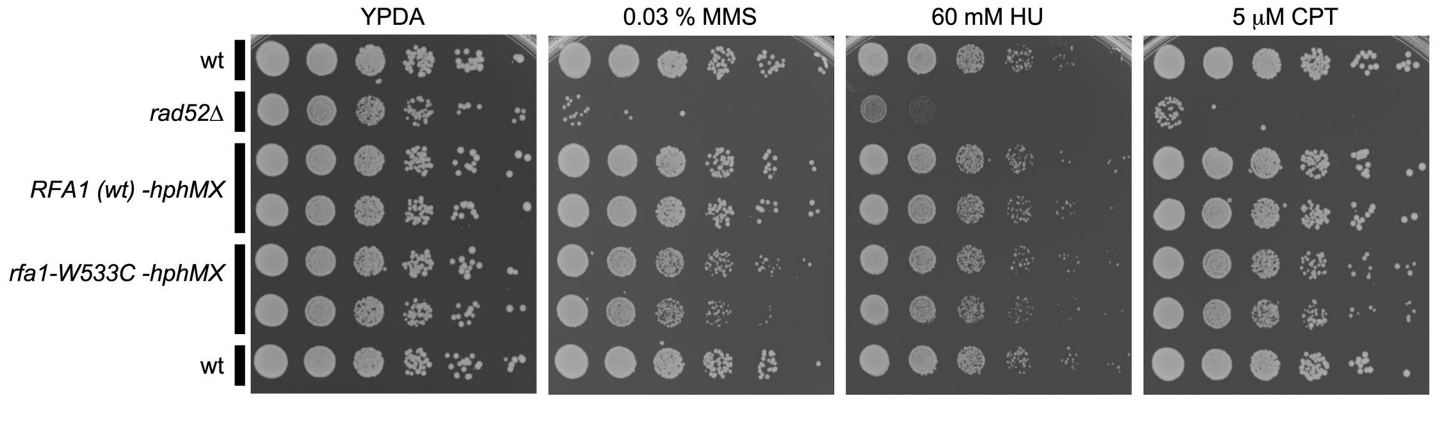


**S10 Fig. Analysis of DNA damage sensitivity for *rfa1-W533C* strains.** Ten-fold serial dilutions of the indicated strains were plated onto YPDA medium with the indicated concentrations of MMS, HU, or CPT. Damage-sensitive *rad52Δ* strain served as a control. Pictures were taken after 2 days at 30 °C.

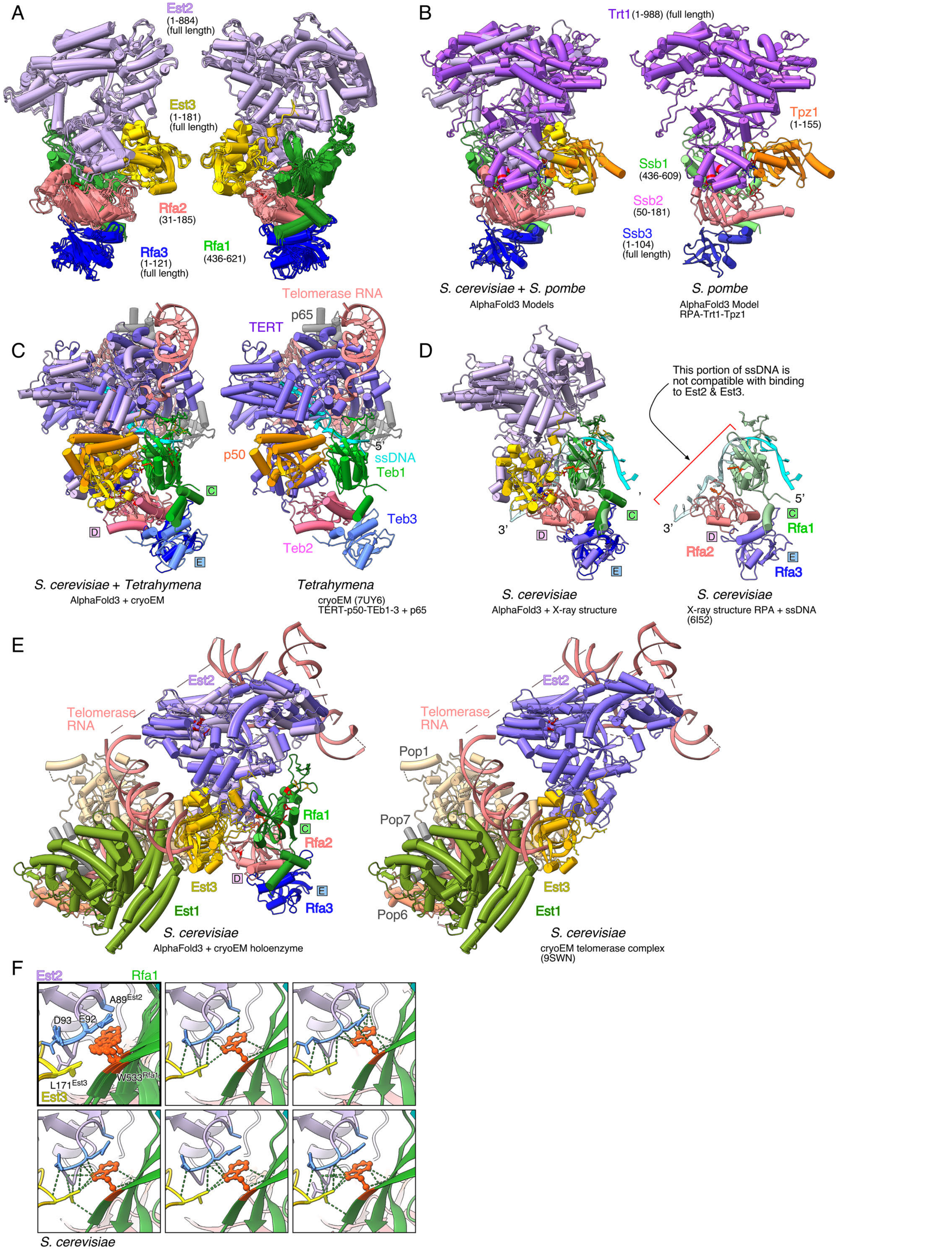
A
Est2
(1-884)
(full length)
Est3
(1-181)
(full length)
Rfa2
(31-185)
Rfa3
(1-121)
(full length)
Rfa1
(436-621)
B
Trt1 (1-988) (full length)
Tpz1
(1-155)
Ssb1
(436-609)
Ssb2
(50-181)
Ssb3
(1-104)
(full length)
S. cerevisiae + S. pombe
AlphaFold3 Models
S. pombe
AlphaFold3 Model
RPA-Trt1-Tpz1
C
Telomerase RNA
p65
TERT
p50
5'
ssDNA
Teb1
Teb3
Teb2
S. cerevisiae + Tetrahymena
AlphaFold3 + cryoEM
Tetrahymena
cryoEM (7UY6)
TERT-p50-TEb1-3 + p65
D
This portion of ssDNA is
not compatible with binding
to Est2 & Est3.
3'
5'
Rfa1
Rfa2
Rfa3
S. cerevisiae
AlphaFold3 + X-ray structure
S. cerevisiae
X-ray structure RPA + ssDNA
(6I52)
E
Est2
Telomerase
RNA
Rfa1
Rfa2
Est3
Est1
Rfa3
S. cerevisiae
AlphaFold3 + cryoEM holoenzyme
Pop1
Pop7
Pop6
S. cerevisiae
cryoEM telomerase complex
(9SWN)
F
Est2
Rfa1
A89Est2
D93
E92
L171Est3
W533Rfa1
Est3
S. cerevisiae

**S11 Fig. AlphaFold3 prediction of the RPA–Est2–Est3 complex. (A)** Overlay of top-scoring AlphaFold3 models for the budding yeast RPA–Est2–Est3 complex, shown from two different viewpoints. **(B)** Superposition of the fission yeast RPA–Trt1–Tpz1 complex with the budding yeast RPA-Est2-Est3 complex. **(C)** Superposition of the *Tetrahymena* Cryo-EM structure with the budding yeast RPA–Est2–Est3 complex. **(D)** Superposition of the budding yeast X-ray structure RPA–ssDNA with the budding yeast RPA–Est2–Est3 complex. **(E)** Superposition of the budding yeast cryo-EM structure of telomerase complex with the budding yeast RPA–Est2–Est3 complex. **(F)** Detailed structures of the Rfa1–Est2 interface. Five top-scoring AlphaFold3 models are shown individually (light frames) or superimposed (bold frame).

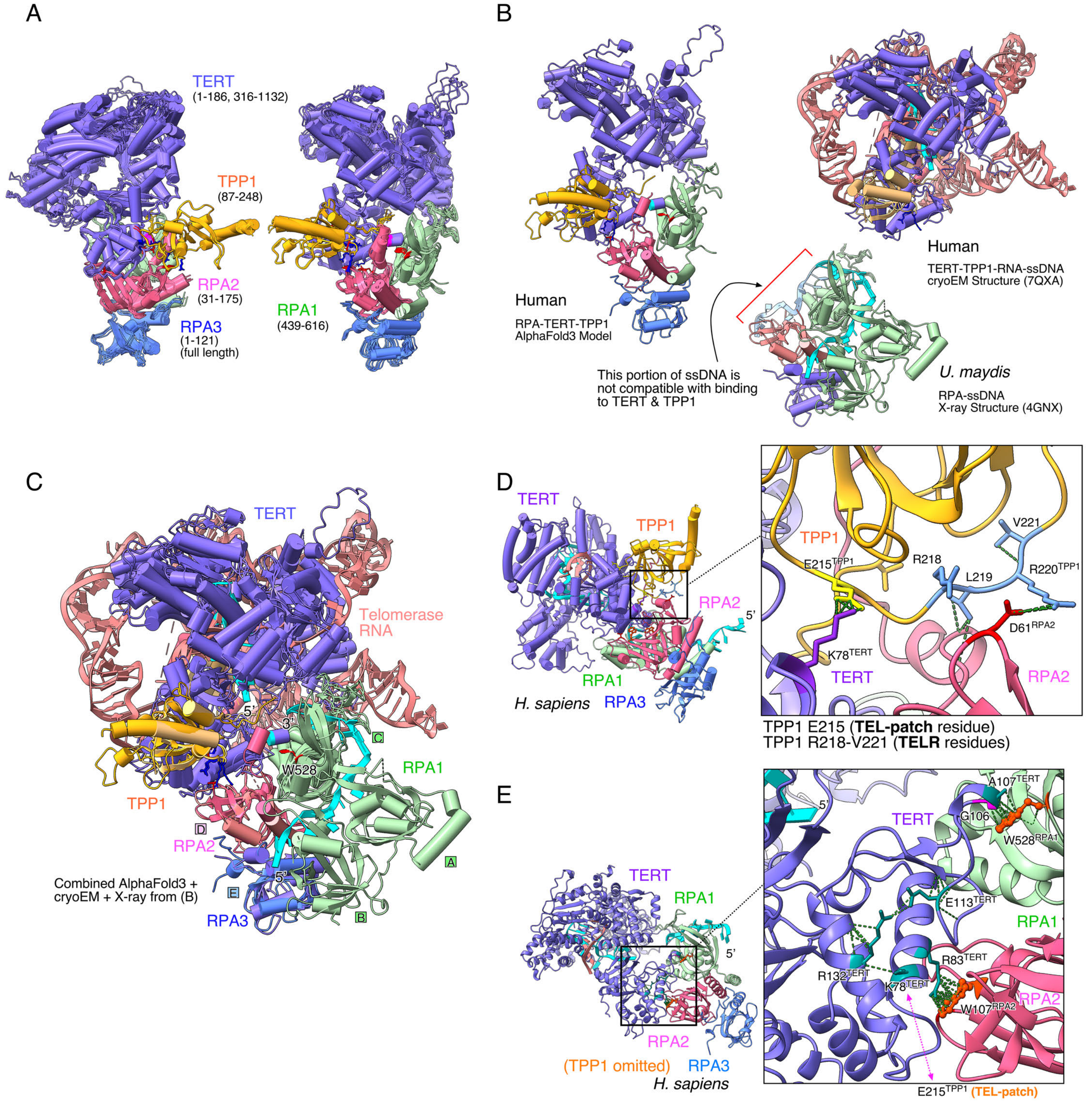


**S12 Fig. AlphaFold3 prediction of the RPA–TERT–TPP1 complex. (A)** Overlay of the top-scoring AlphaFold3 models for the human RPA–TERT–TPP1 complex, shown from two different viewpoints. **(B, C)** Human RPA–TERT–TPP1 model from AlphaFold3, and Human TERT–TPP1 cryo-EM structure, and RPA–ssDNA X-ray structure from *U.* maydis, shown individually (B) or superimposed (C). **(D)** Detailed view of the predicted human D61$^{RPA2}$–R220$^{TPP1}$ interaction, supported by neighboring residues, together with K78$^{TERT}$–E215$^{TTP1}$ interactions. **(E)** Detailed view of the predicted TERT-G106/A107 interaction with RPA1-W528, stabilized by intramolecular TERT interactions involving E113, R132, and K78, as well as the TPP1-E215 interaction. TPP1 is omitted from this panel to help visualize the relevant interactions.

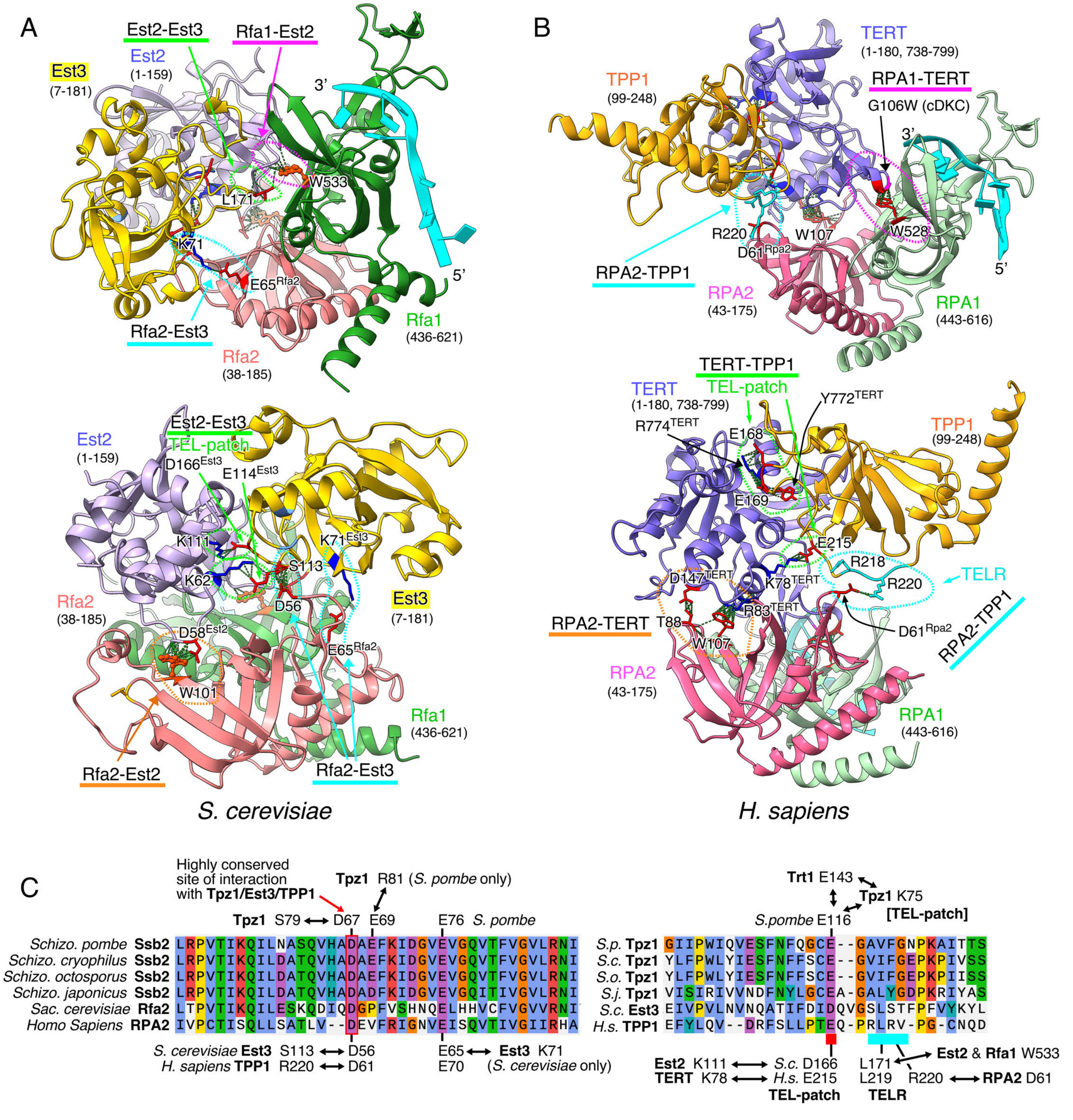


**S13 Fig. Comparison to RPA–TERT–TPP1-like complex models from budding yeast and humans. (A, B)** Functionally important protein-protein interactions identified in the fission yeast RPA–Trt1–Tpz1 complex are conserved in budding yeast (A) and humans (B). Rfa1–Est2/RPA1–TERT interactions are shown in pink, Rfa2–Est2/RPA2–TERT in yellow, Rfa2–Est3/RPA2–TPP1 in blue, and Est2–Est3/TERT–TPP1 in green. Interacting sites are indicated. **(C)** Sequence alignments for Ssb2/Rfa2/RPA2 (left) and Tpz1/Est3/TPP1 (right). Interactions of individual residues with other complex partners are indicated.

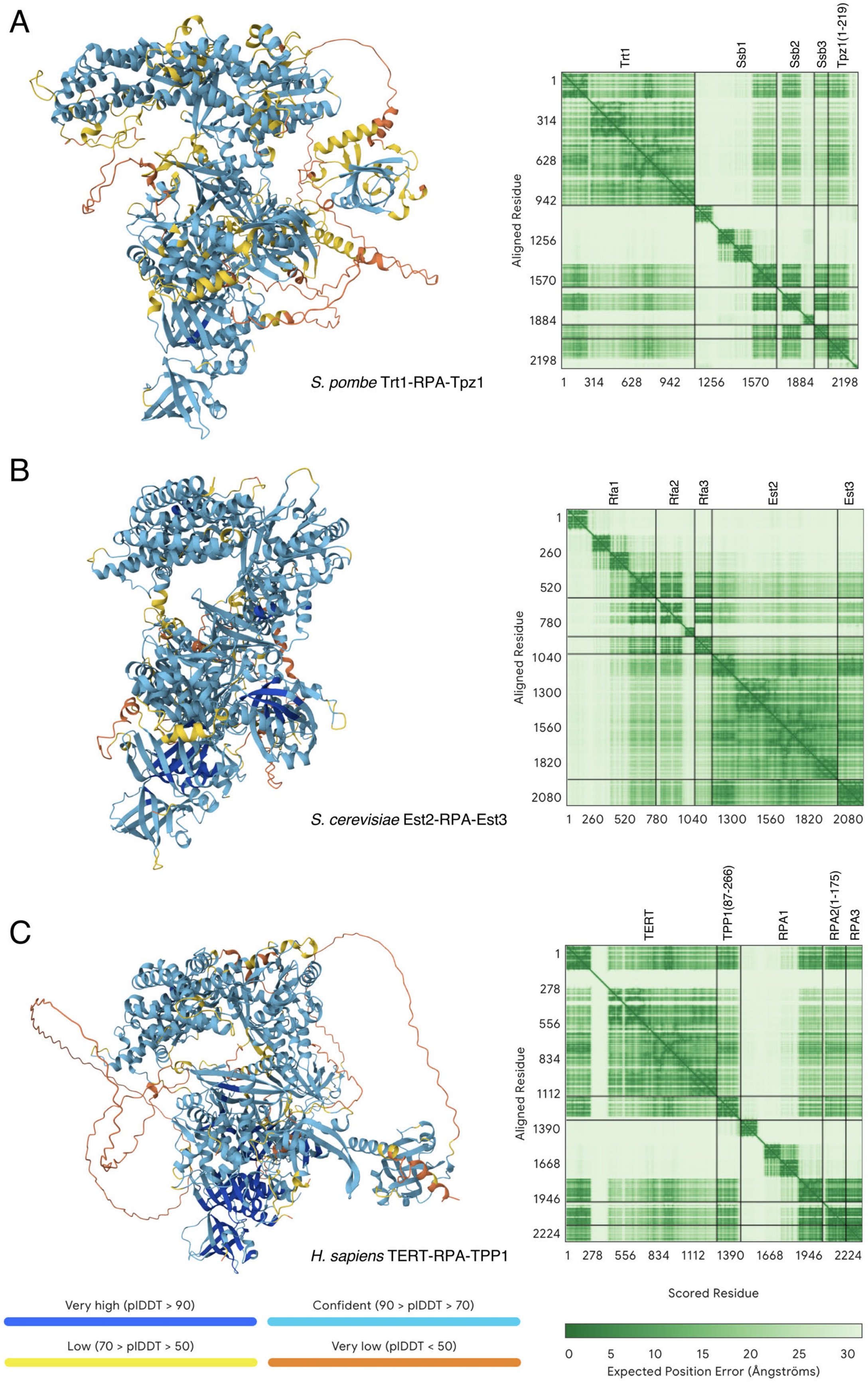


**S14 Fig. Outputs from the AlphaFold3 server. (A)** *S. pombe* Trt1-RPA-Tpz1. **(B)** *S. cerevisiae* Est2-RPA-Est3. **(C)** *H. sapiens* TERT-RPA-TPP1.
