## Supplementary material for "Fission yeast RPA–TERT–Tpz1^TPP1^ complex promotes telomere extension and suppresses telomere recombination": S2 Data: un-cropped images files

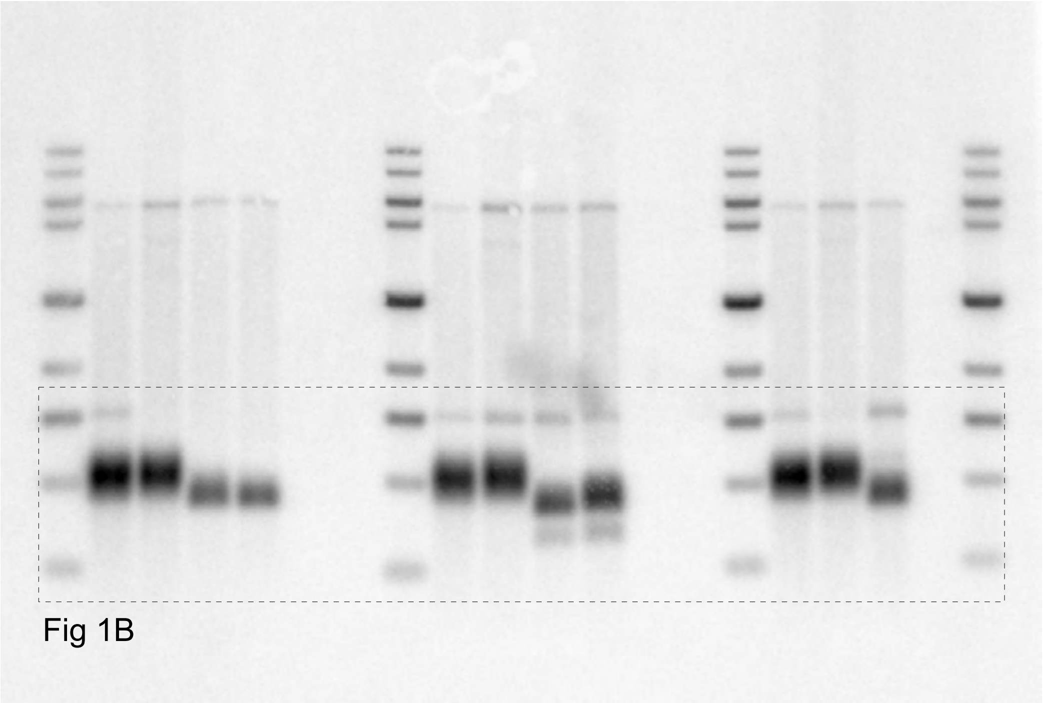
Fig 1B

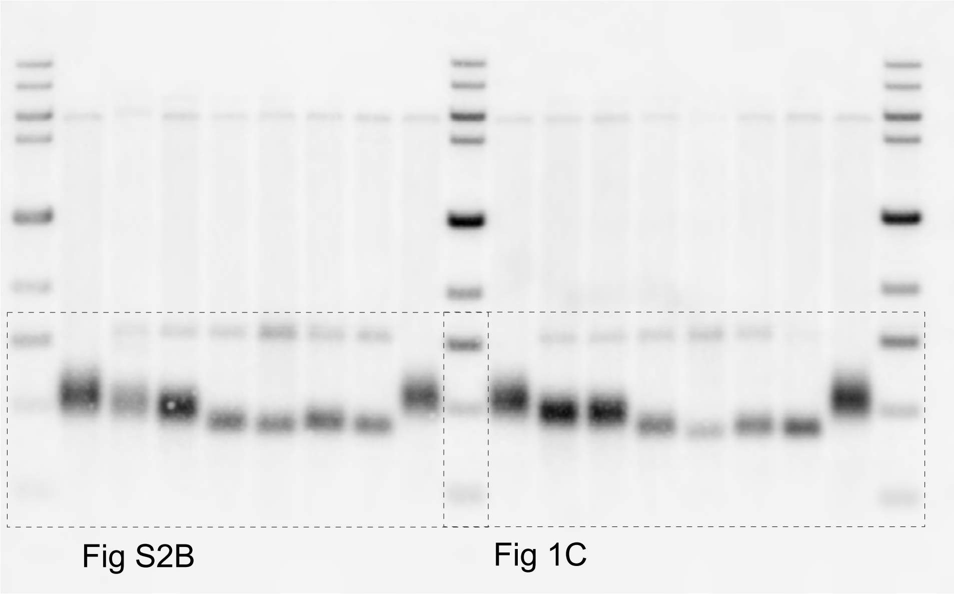
Fig S2B
Fig 1C

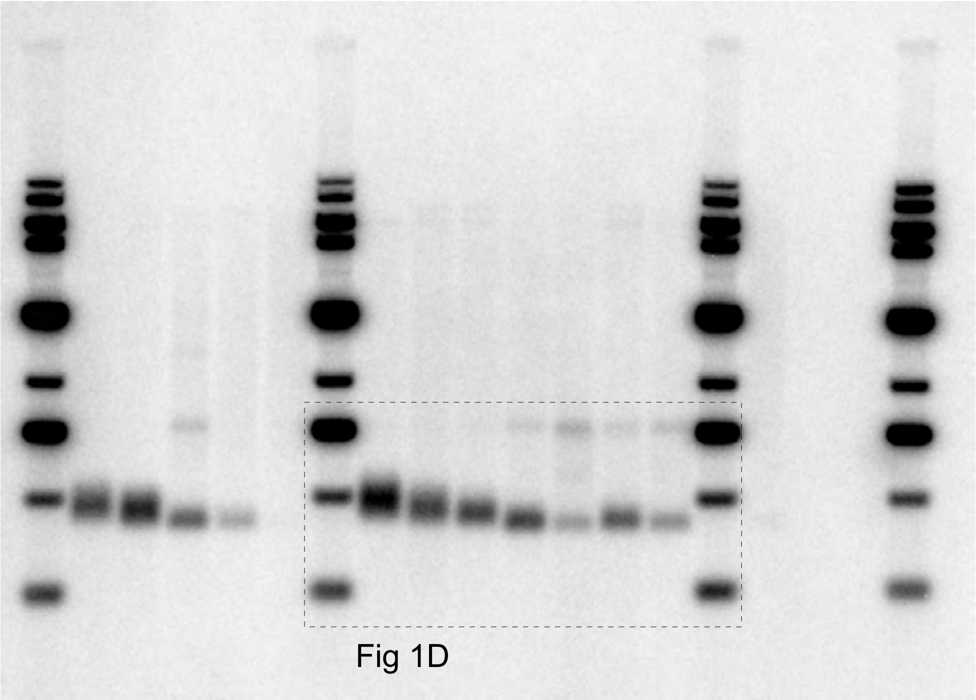
Fig 1D

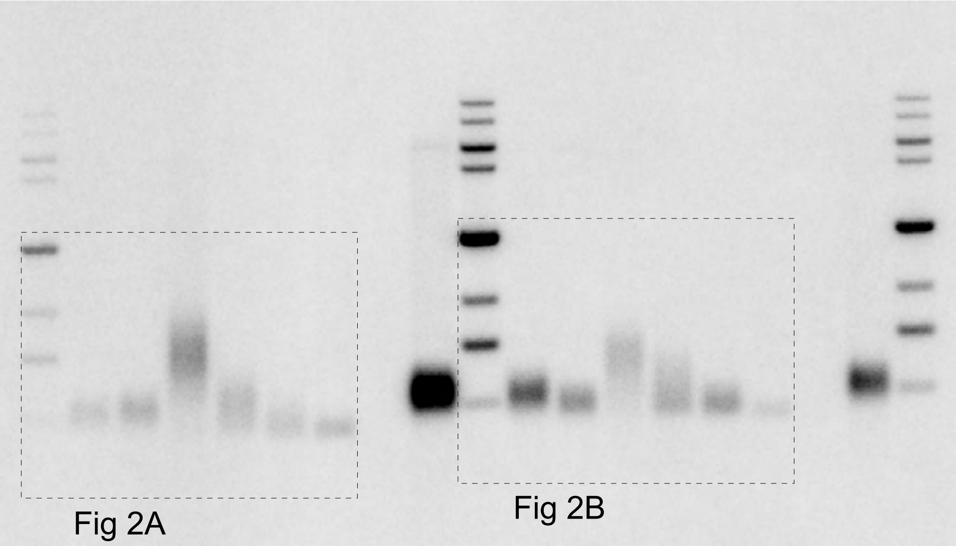
Fig 2A
Fig 2B

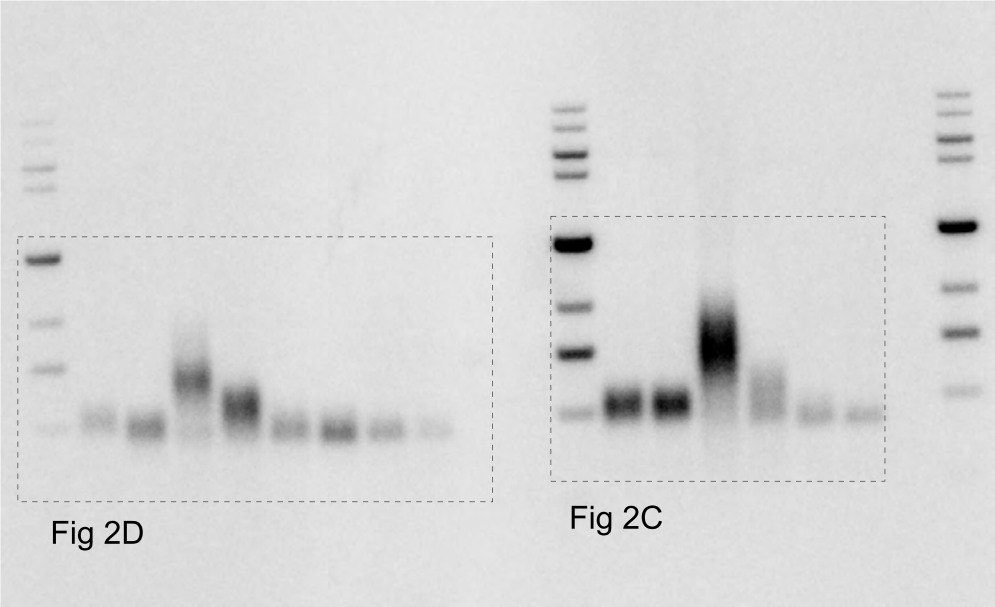
Fig 2D
Fig 2C

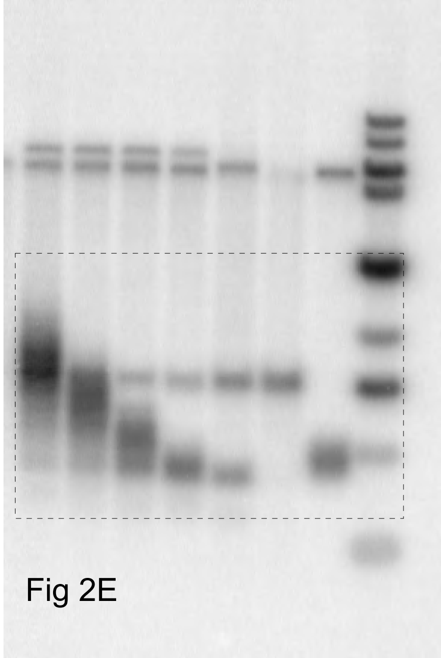
Fig 2E

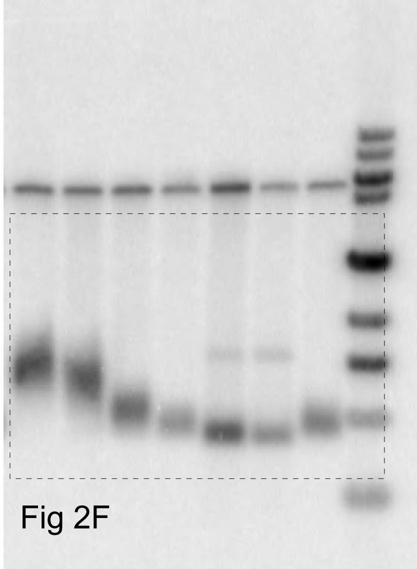
Fig 2F

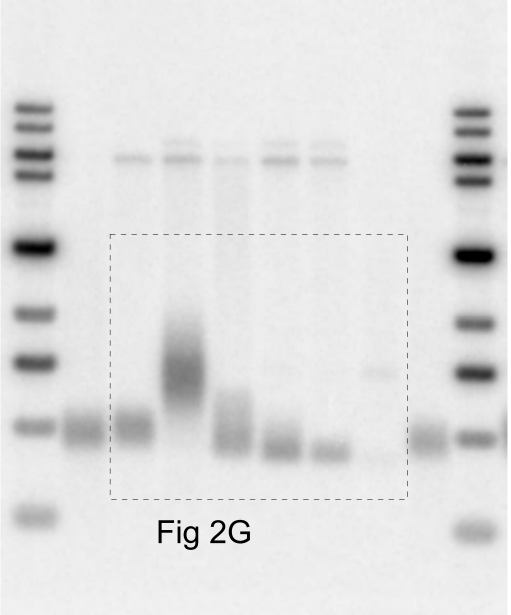
Fig 2G

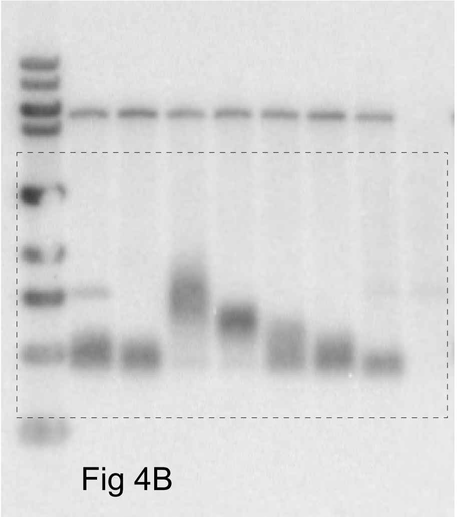
Fig 4B

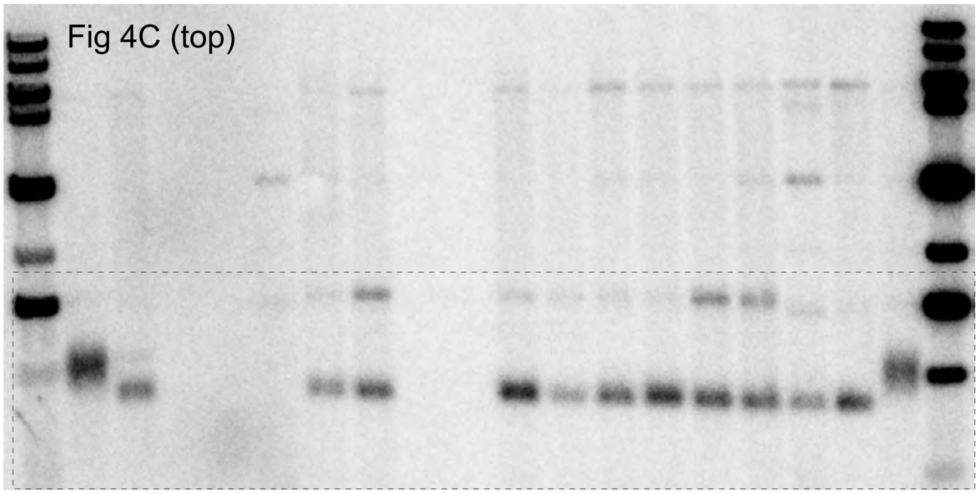
Fig 4C (top)

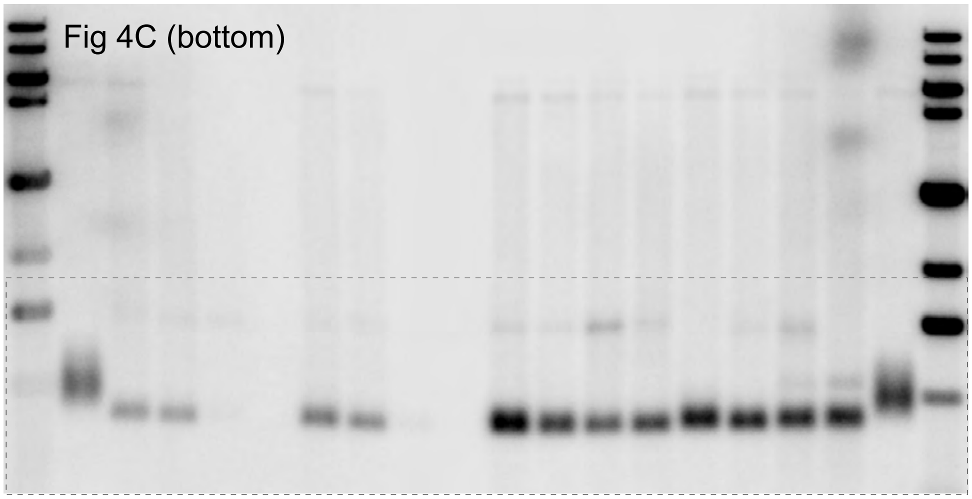
Fig 4C (bottom)

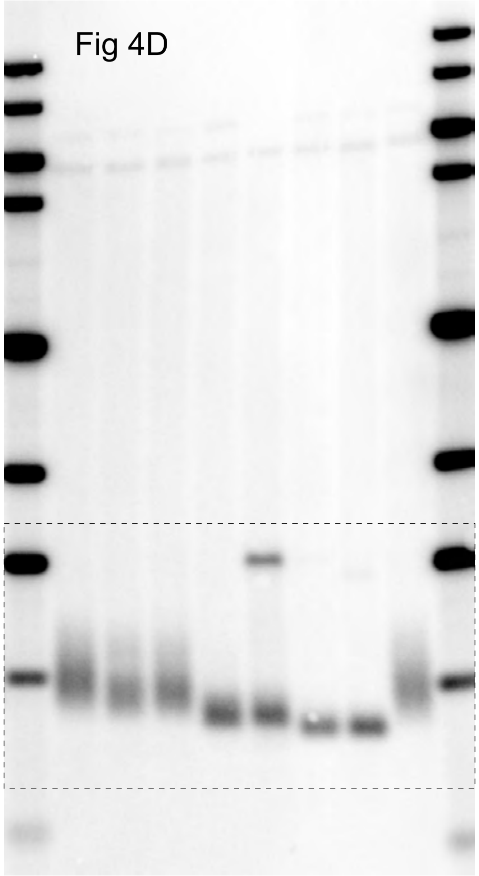
Fig 4D

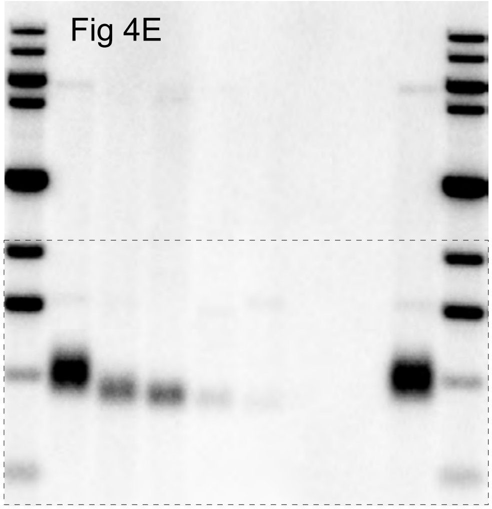
Fig 4E

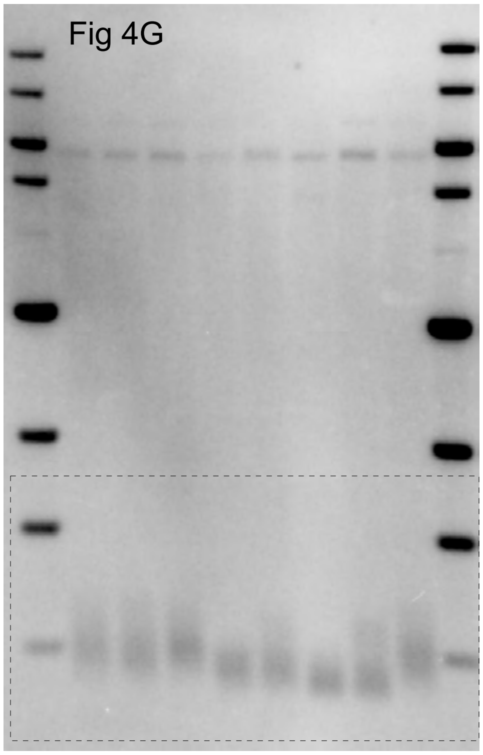
Fig 4G

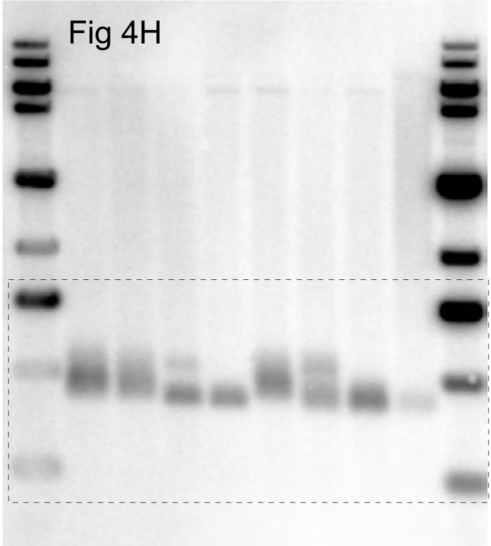
Fig 4H

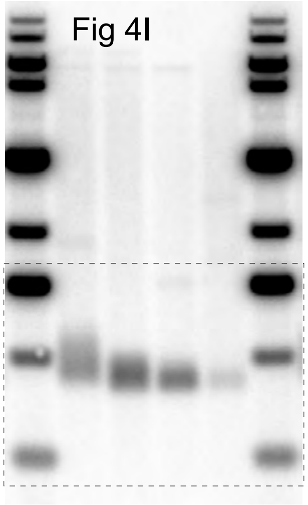
Fig 4I

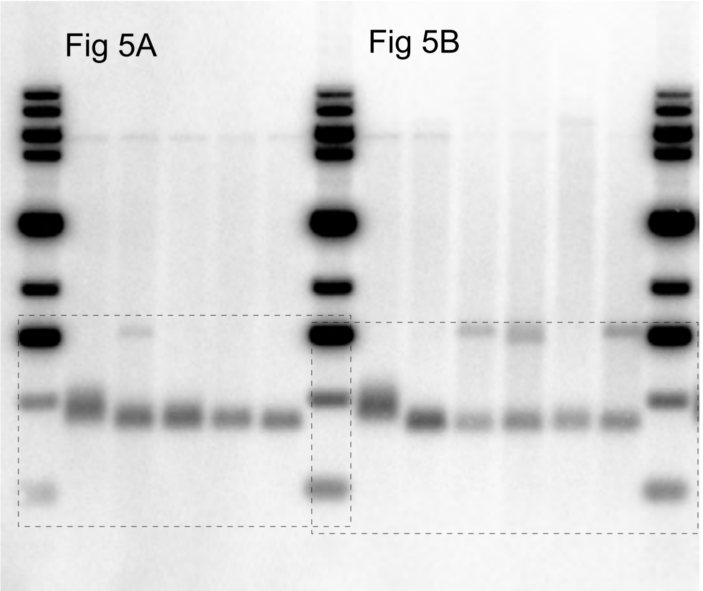
Fig 5A
Fig 5B

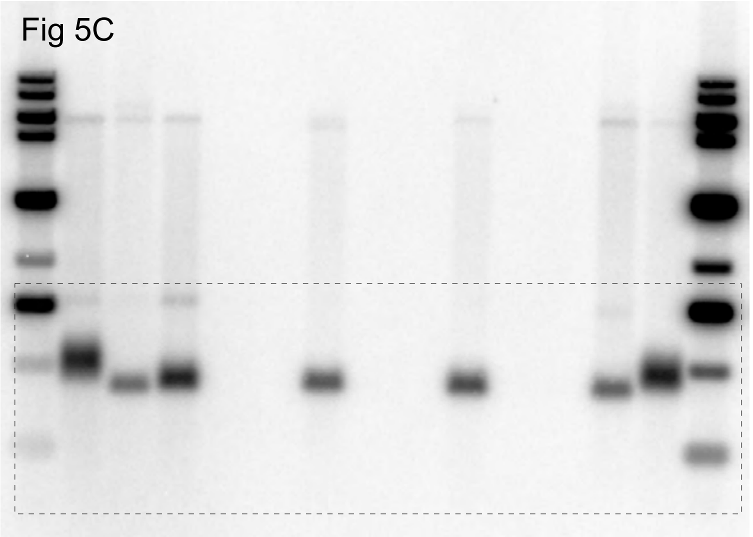
Fig 5C

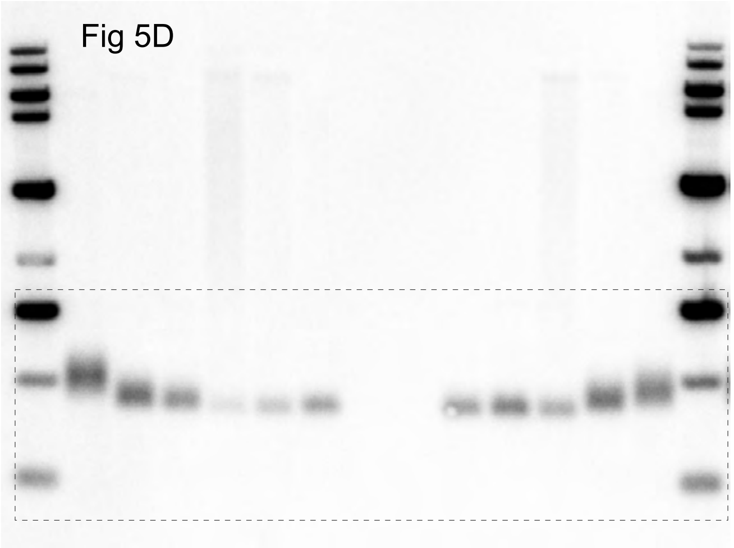
Fig 5D

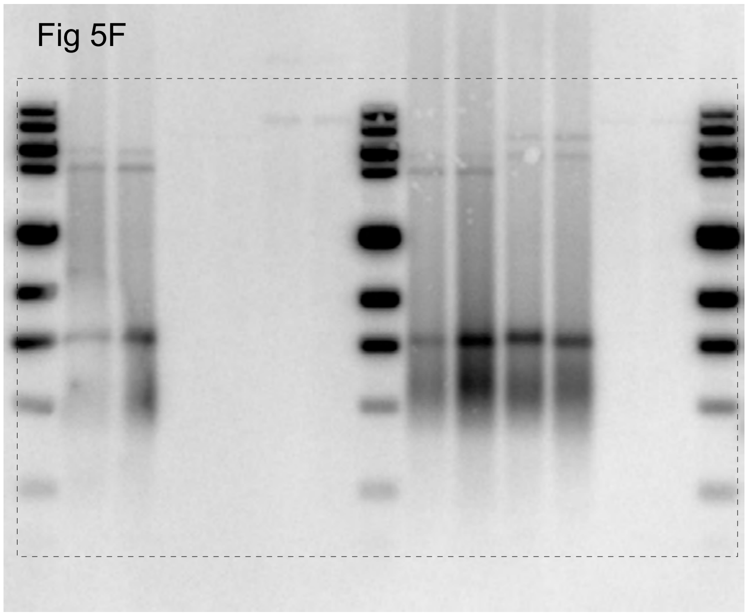
Fig 5F

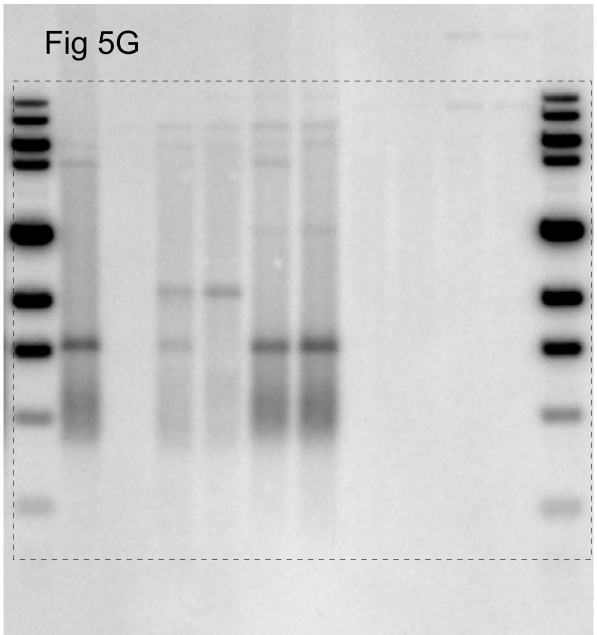
Fig 5G

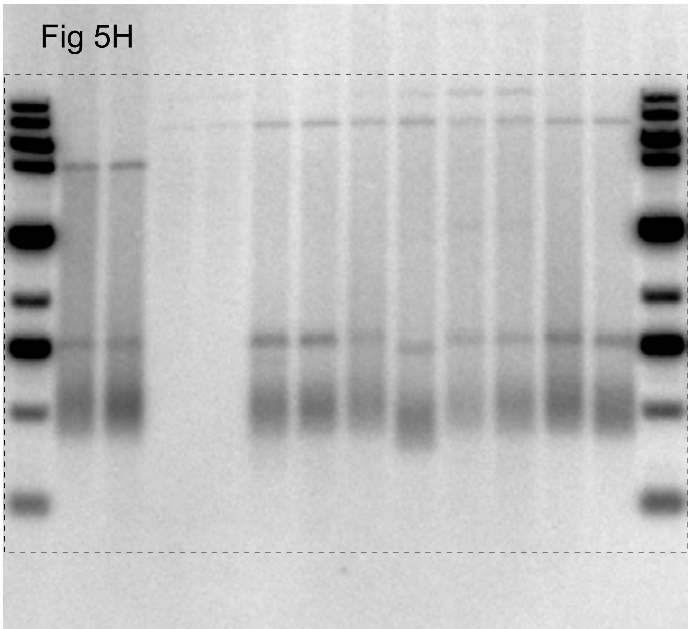
Fig 5H

Fig 6B

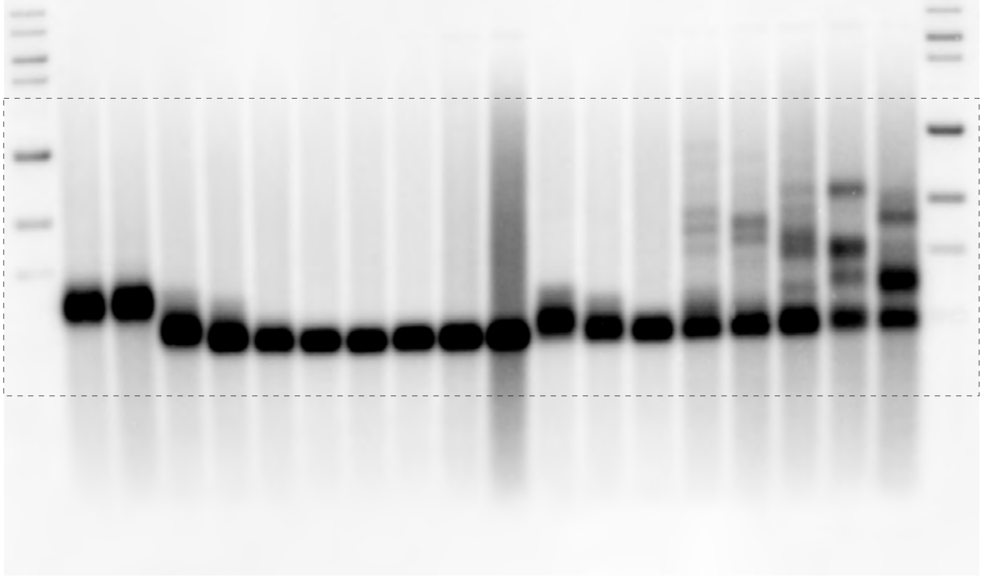

Fig 6C

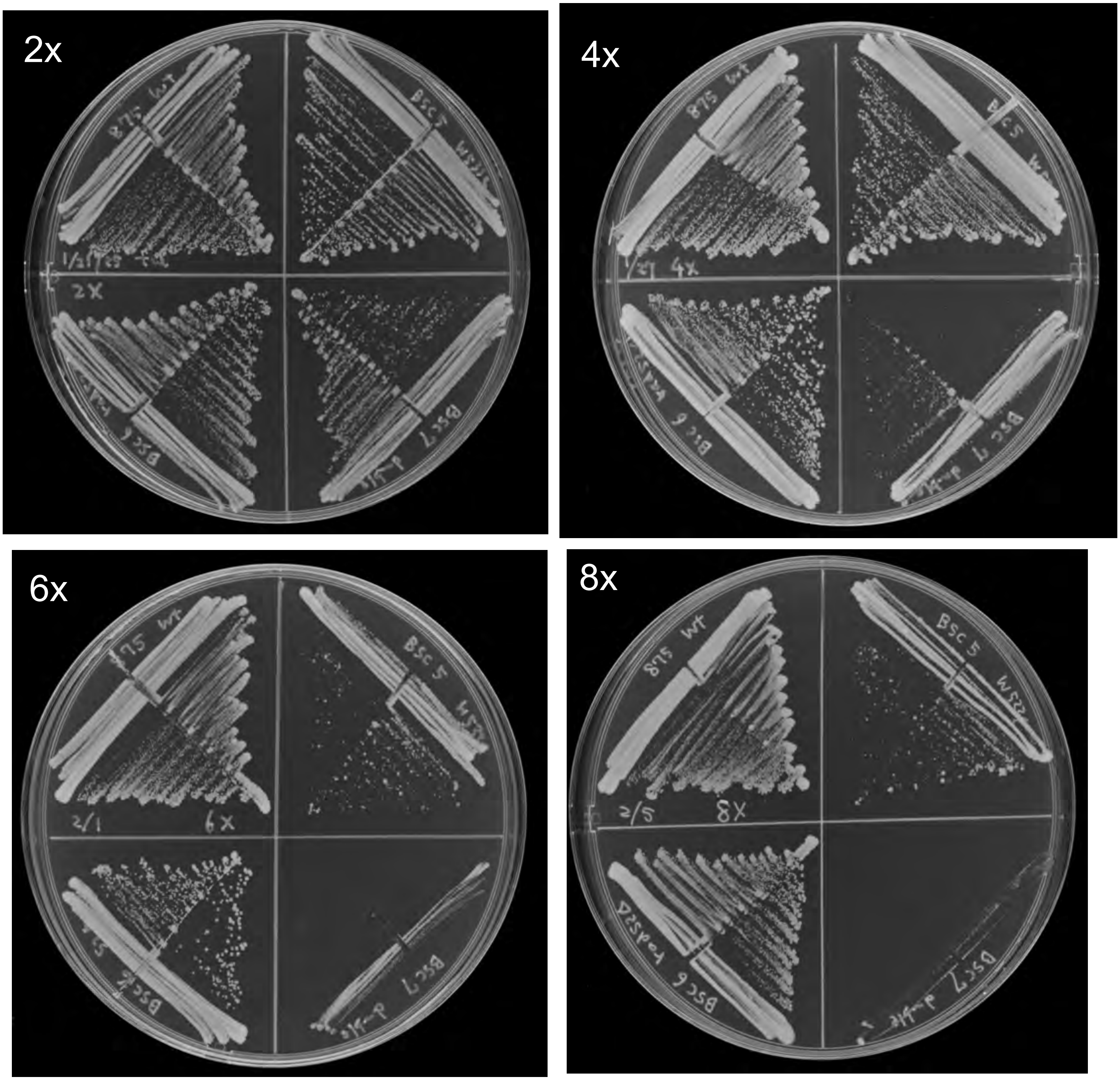

Fig 6E

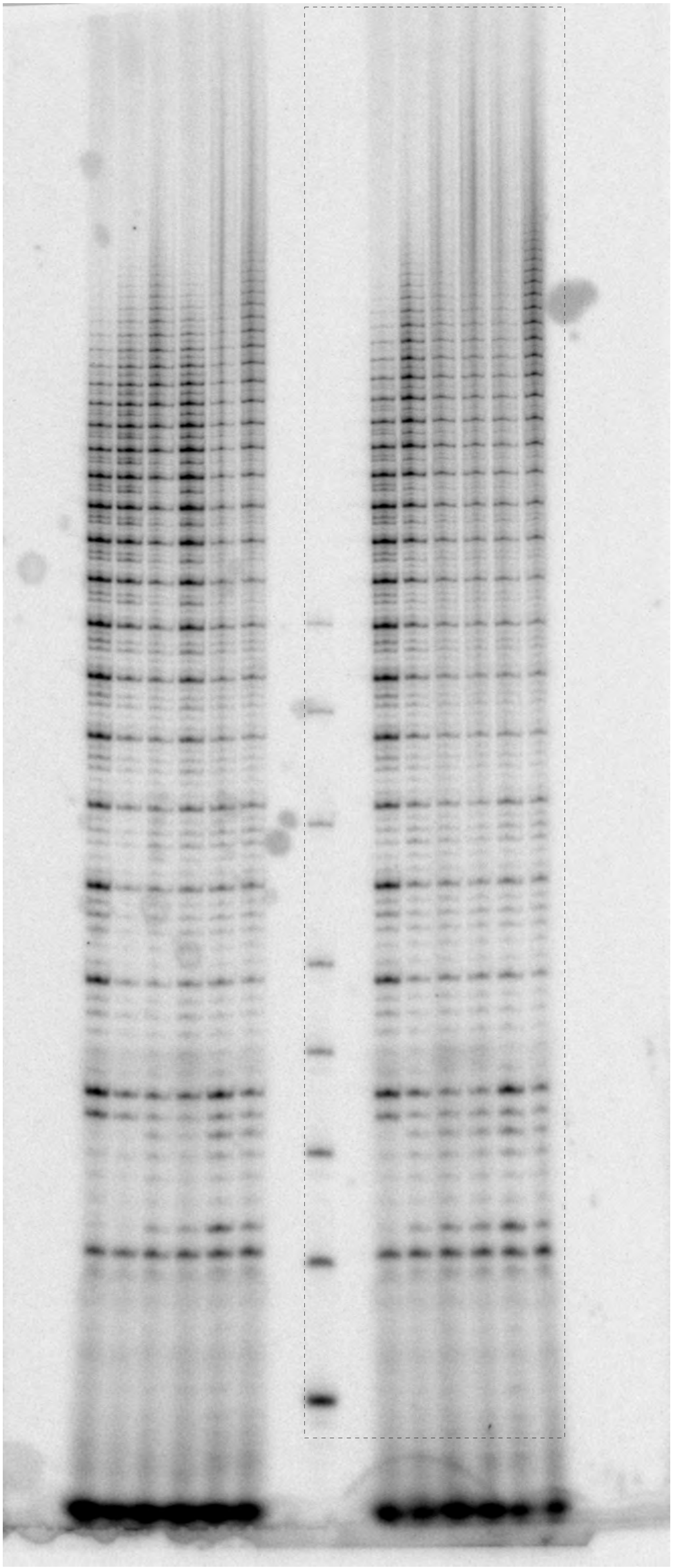

Fig S1B

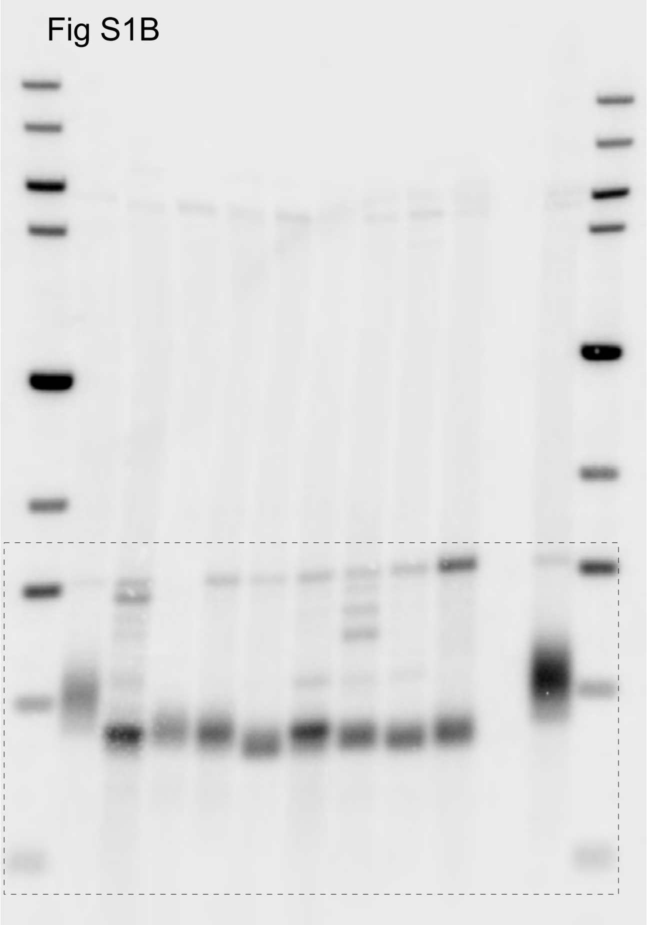

Fig S1D

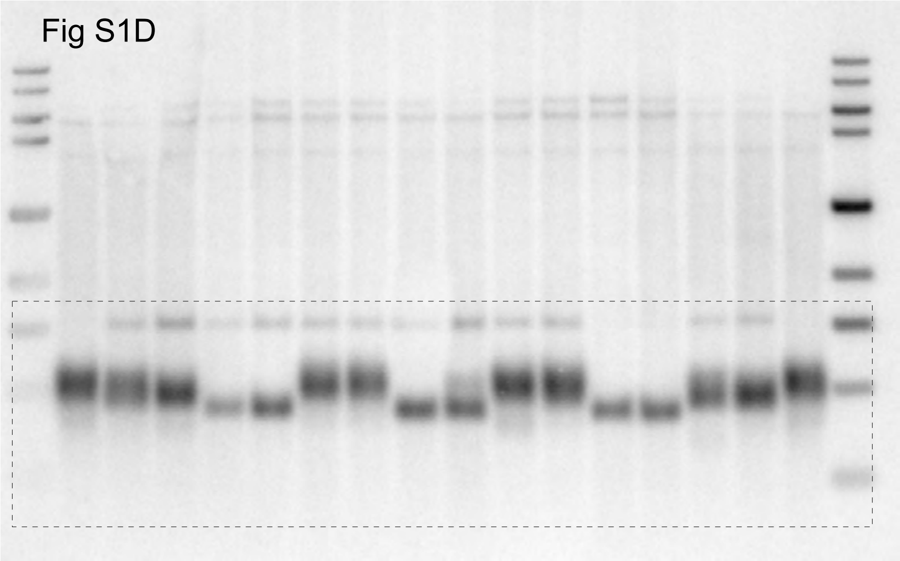

Fig S1D

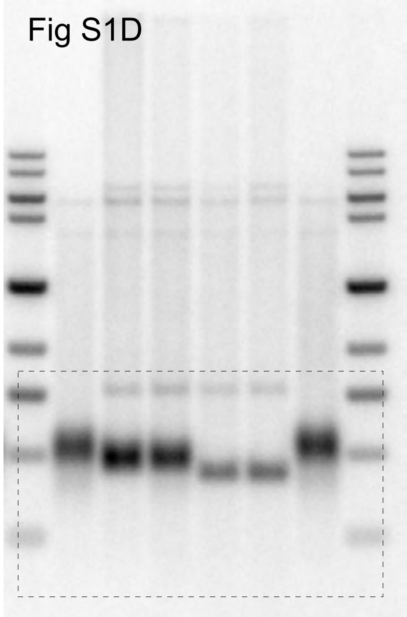

Fig S1C

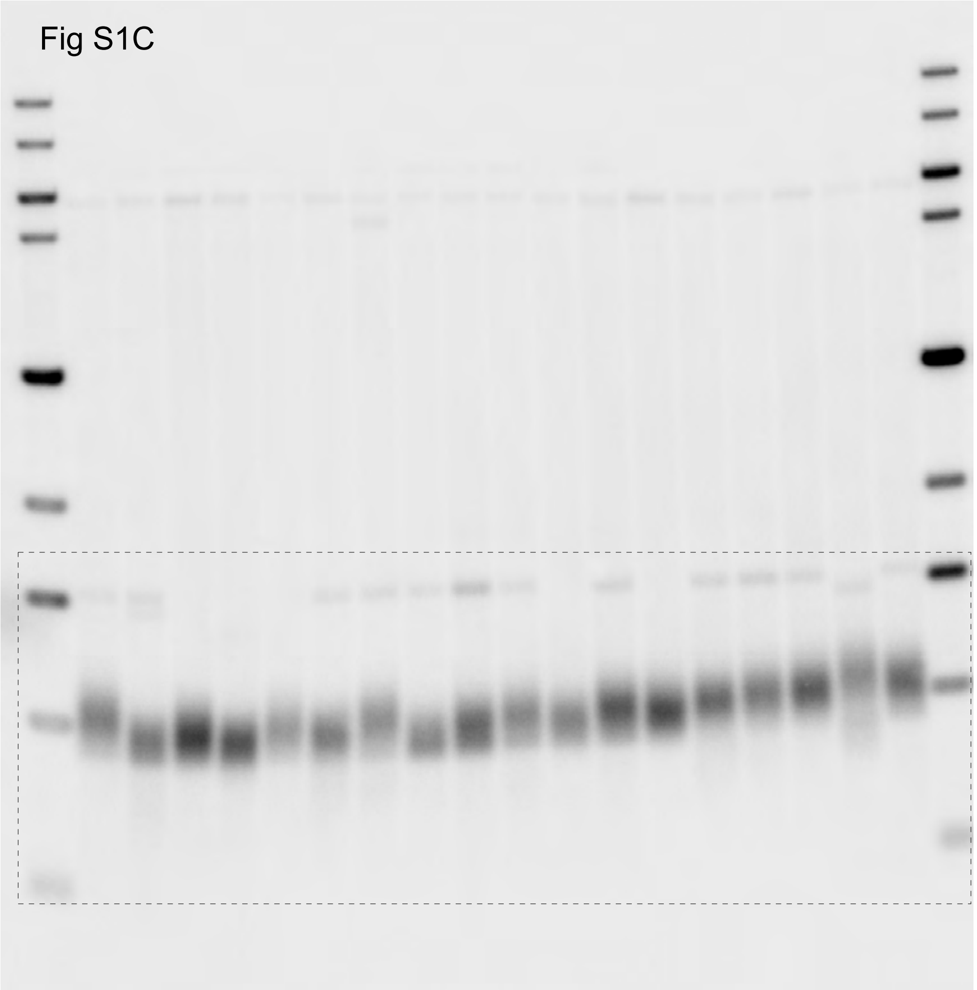

Fig S1E

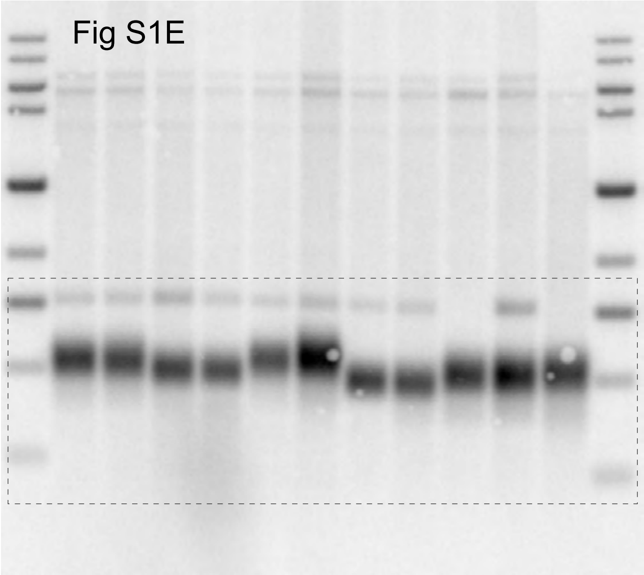

Fig S2D

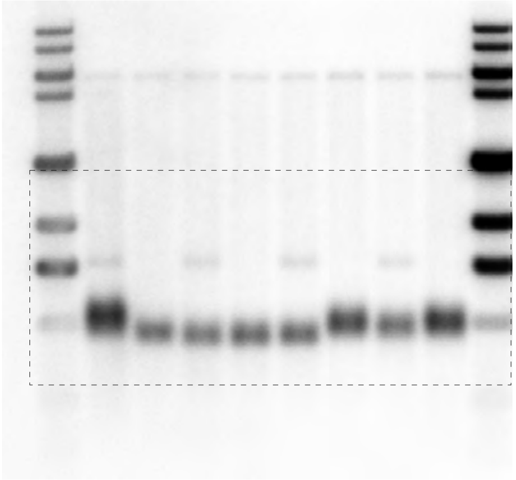

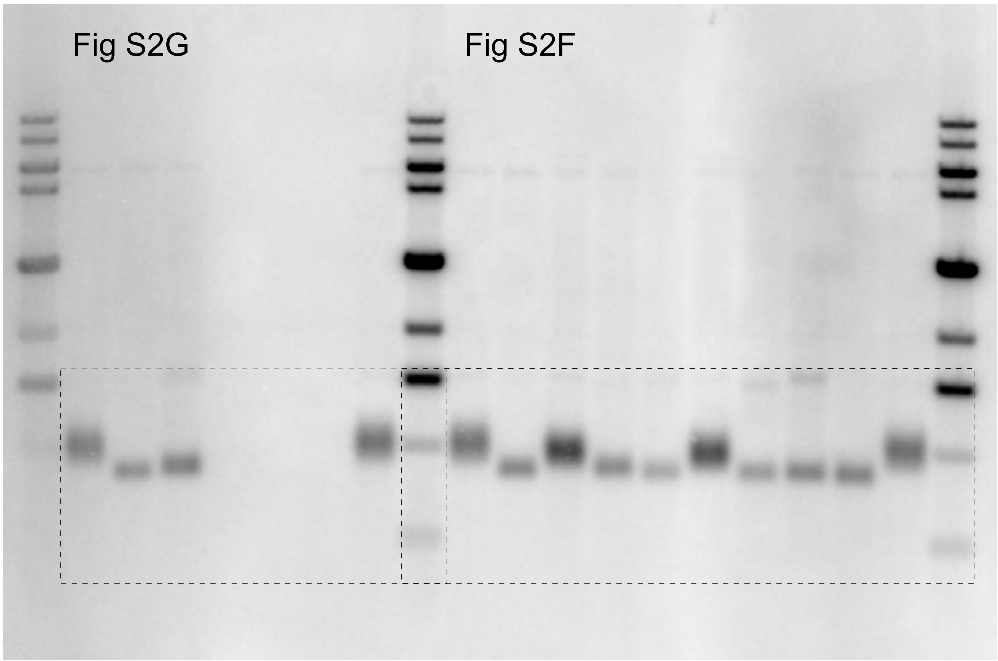


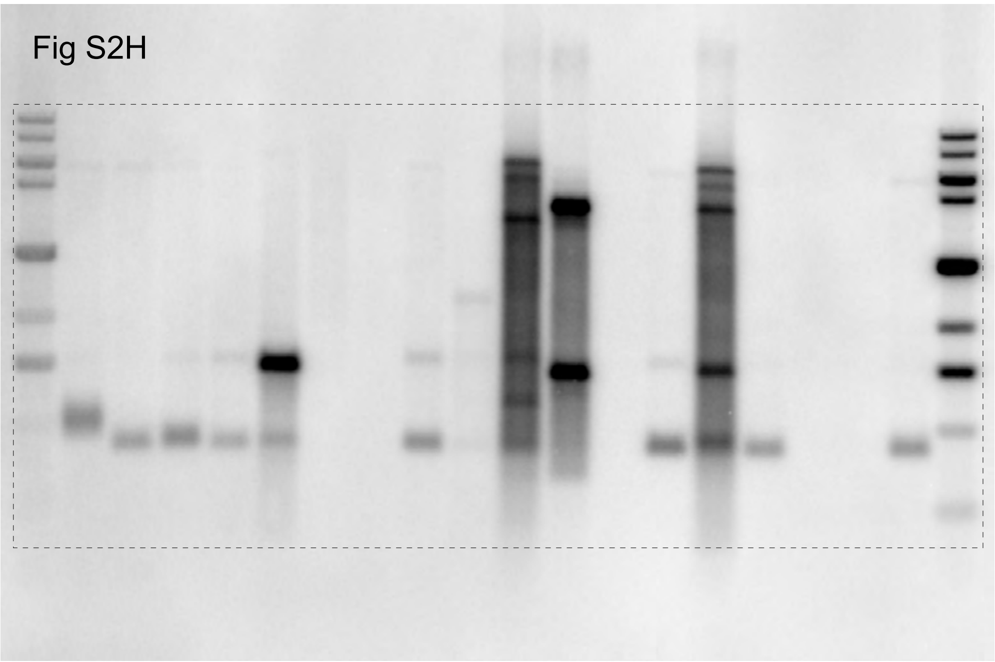

Fig S2C

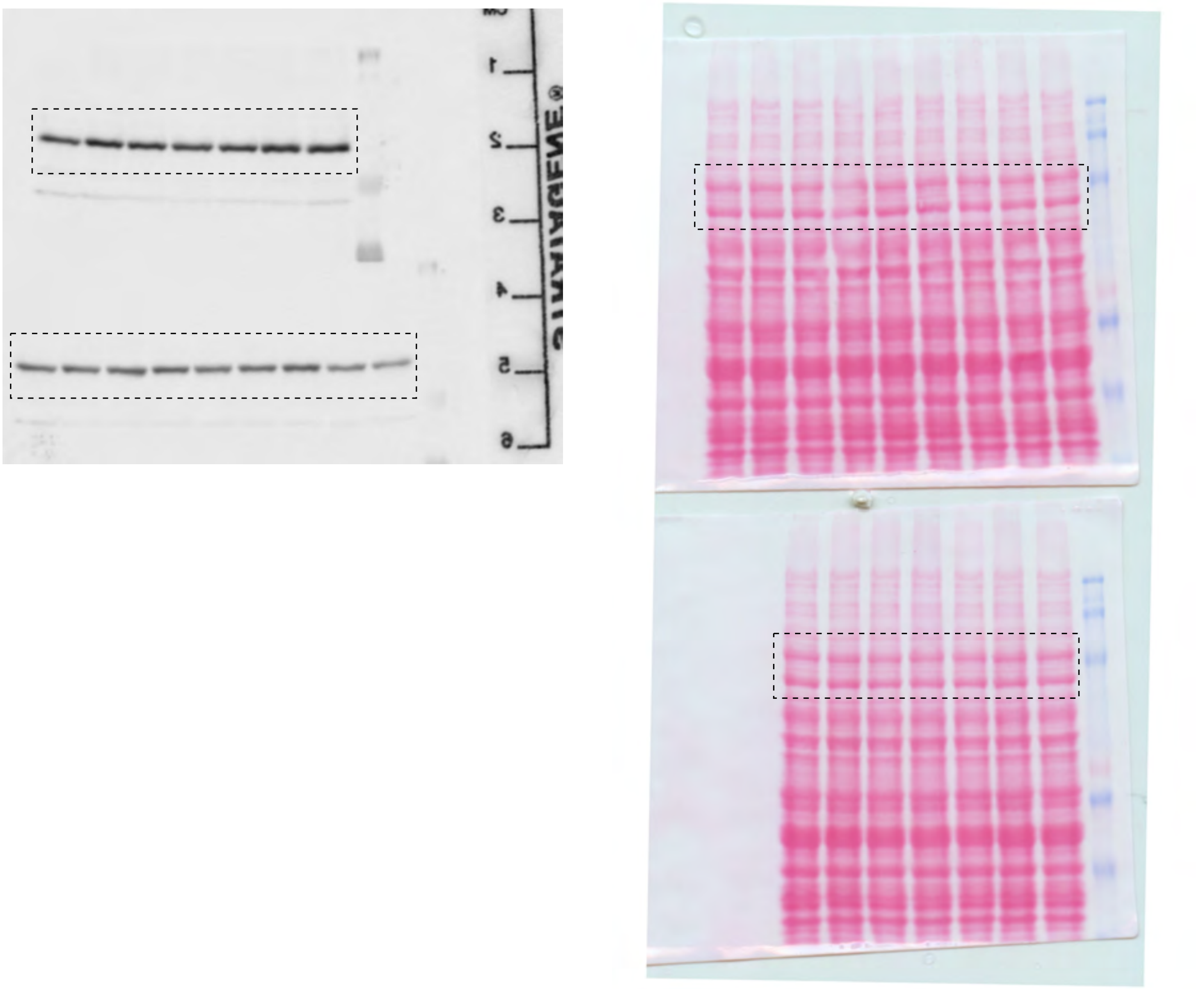

Fig S4A

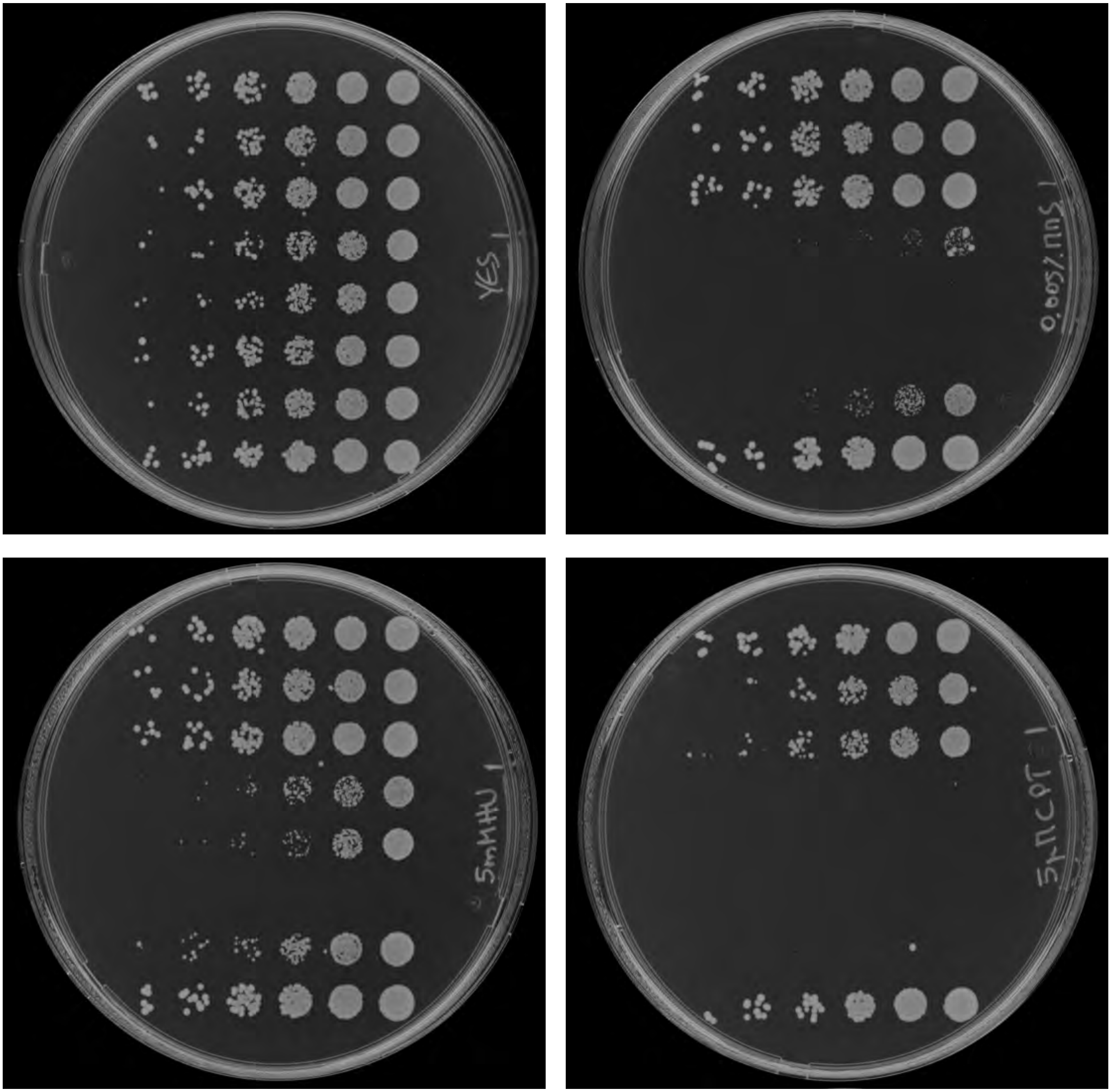

Fig S4B

*ssb3-GFP*

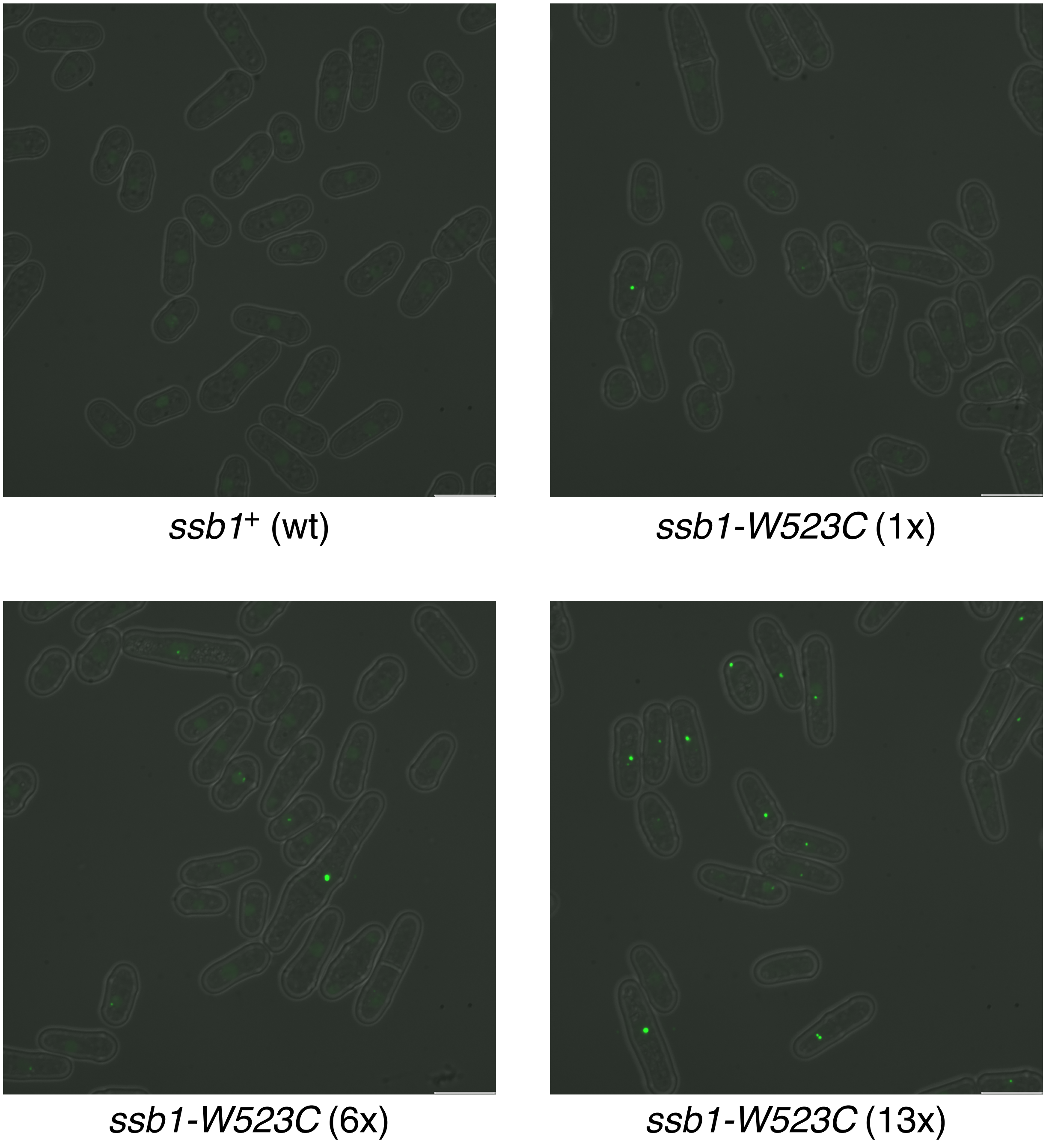

Fig S5B

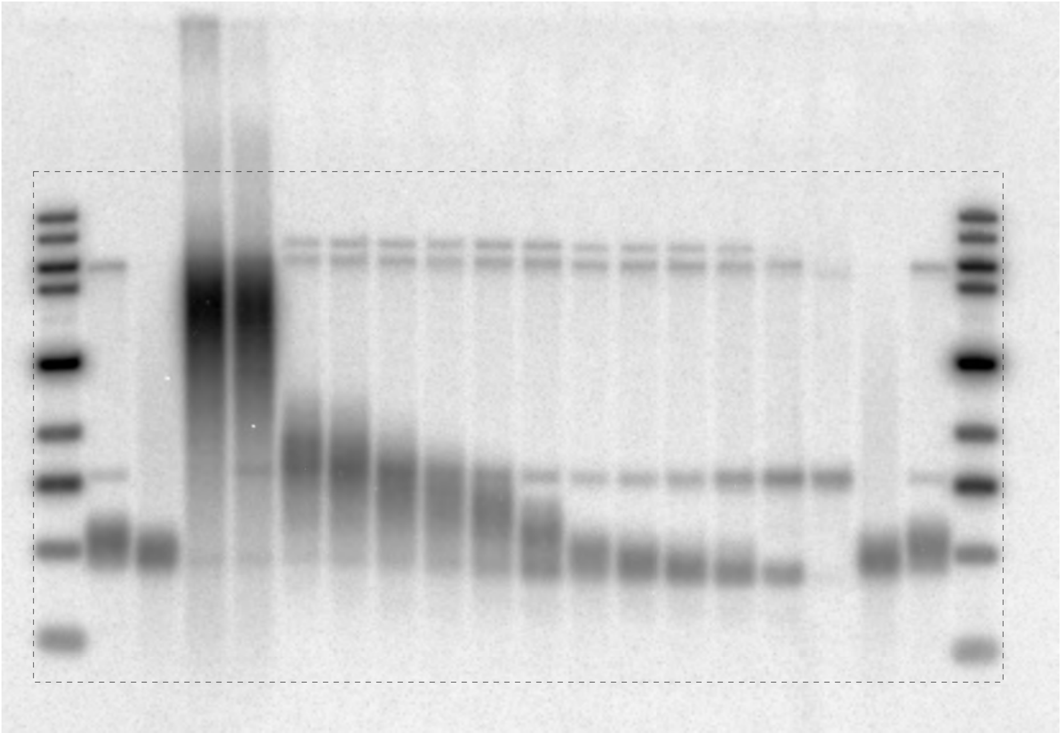

Fig S5C

Fig S5D

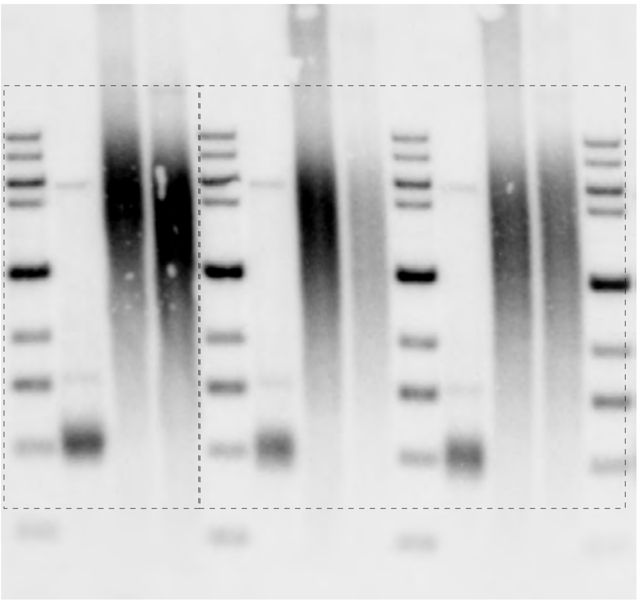

Fig S5E

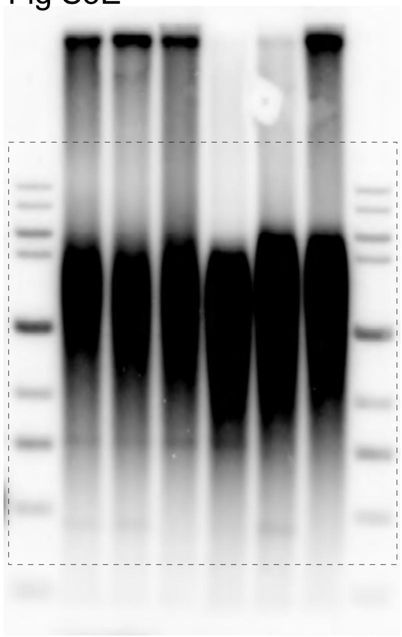

Fig S7B

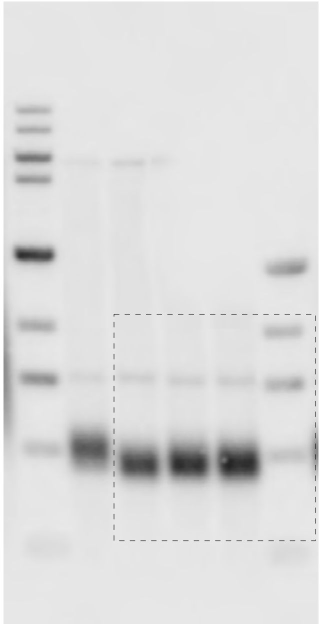

Fig S7C

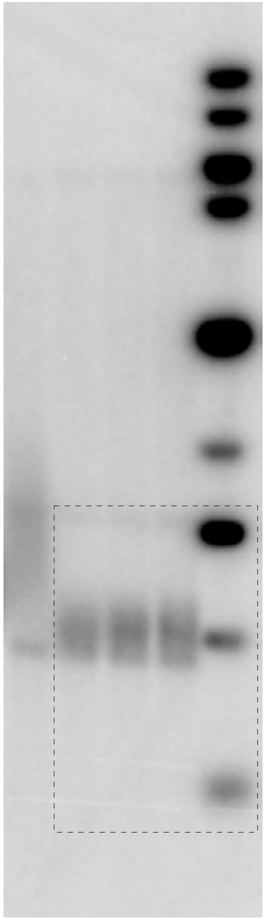

Fig S8A

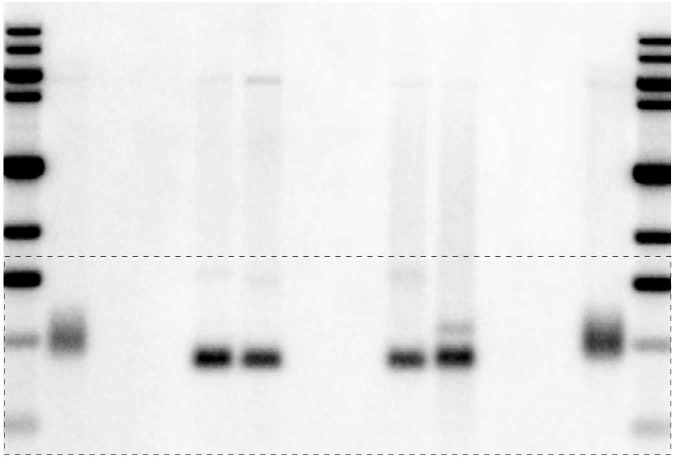

Fig S8B

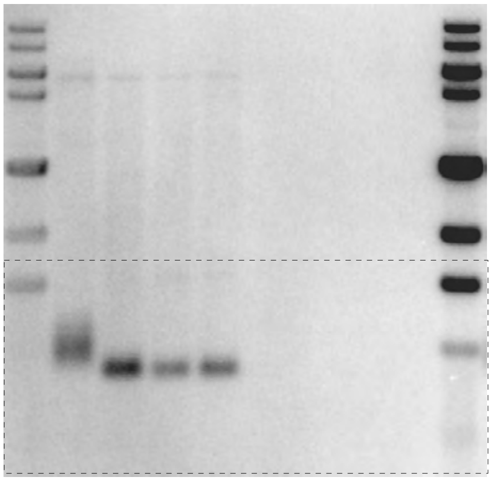

Fig S8C

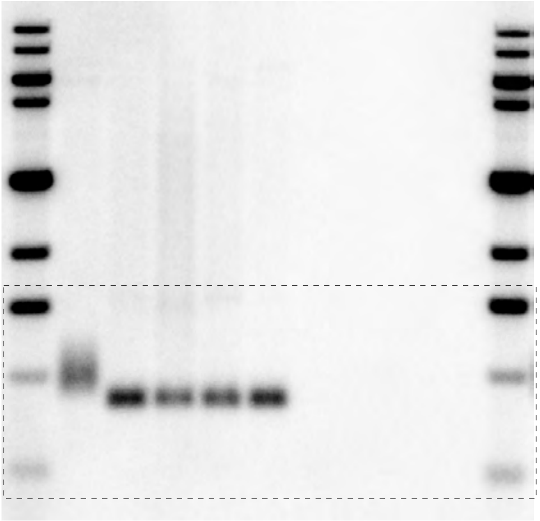

Fig S8D

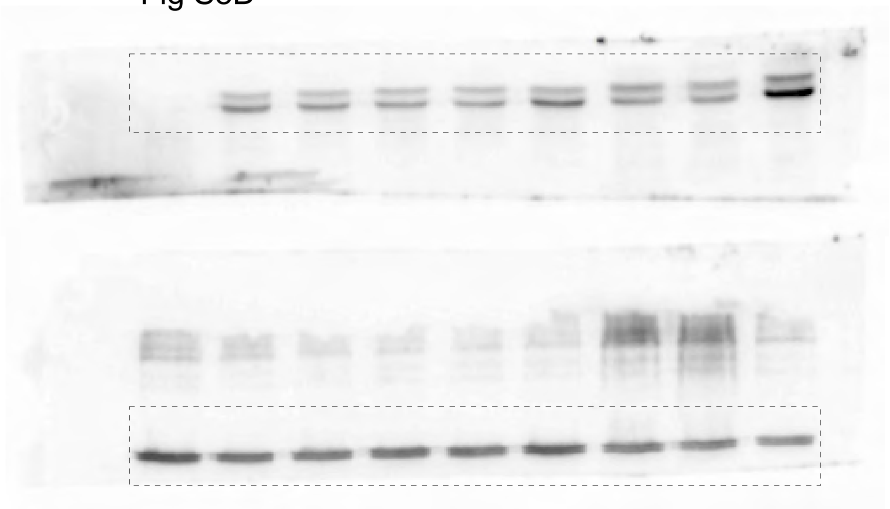

Fig S8G

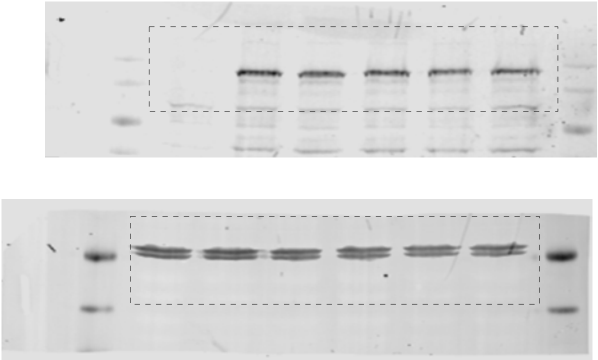

Fig S8E

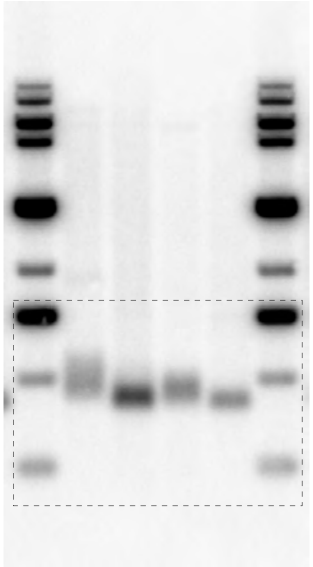

Fig S8F (left)

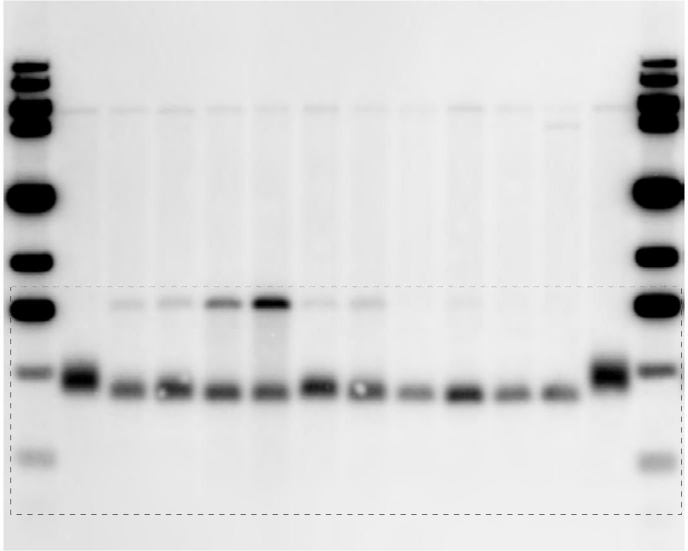

Fig S8F (right)

Fig S8H (right)

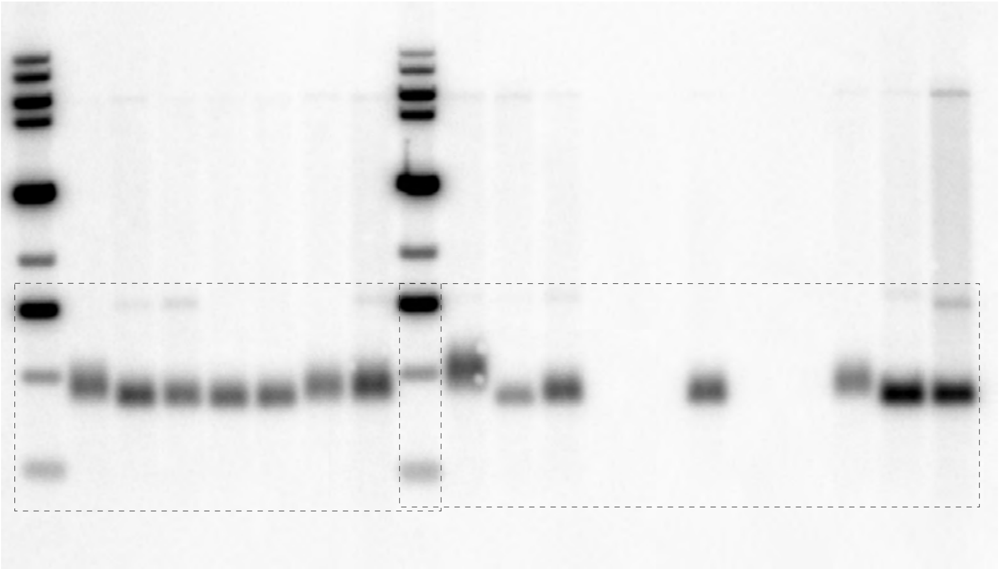

Fig S8H (left)

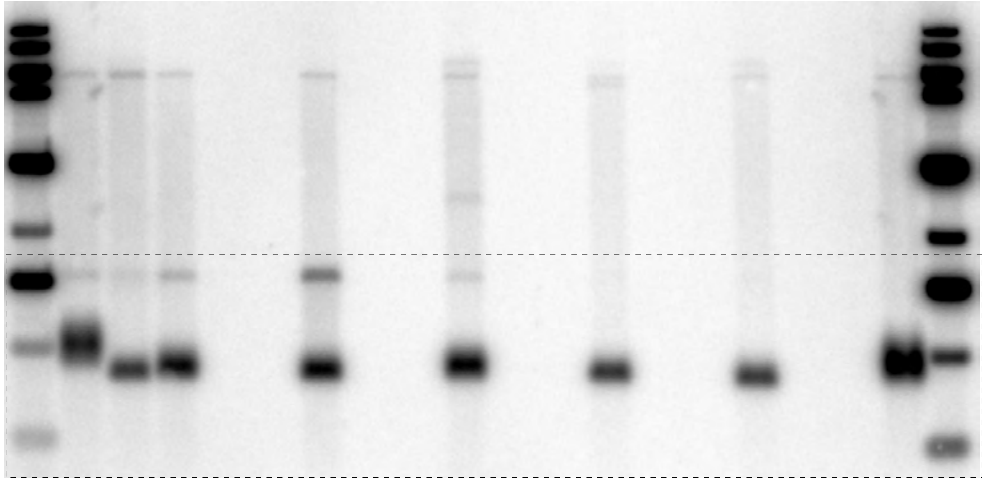

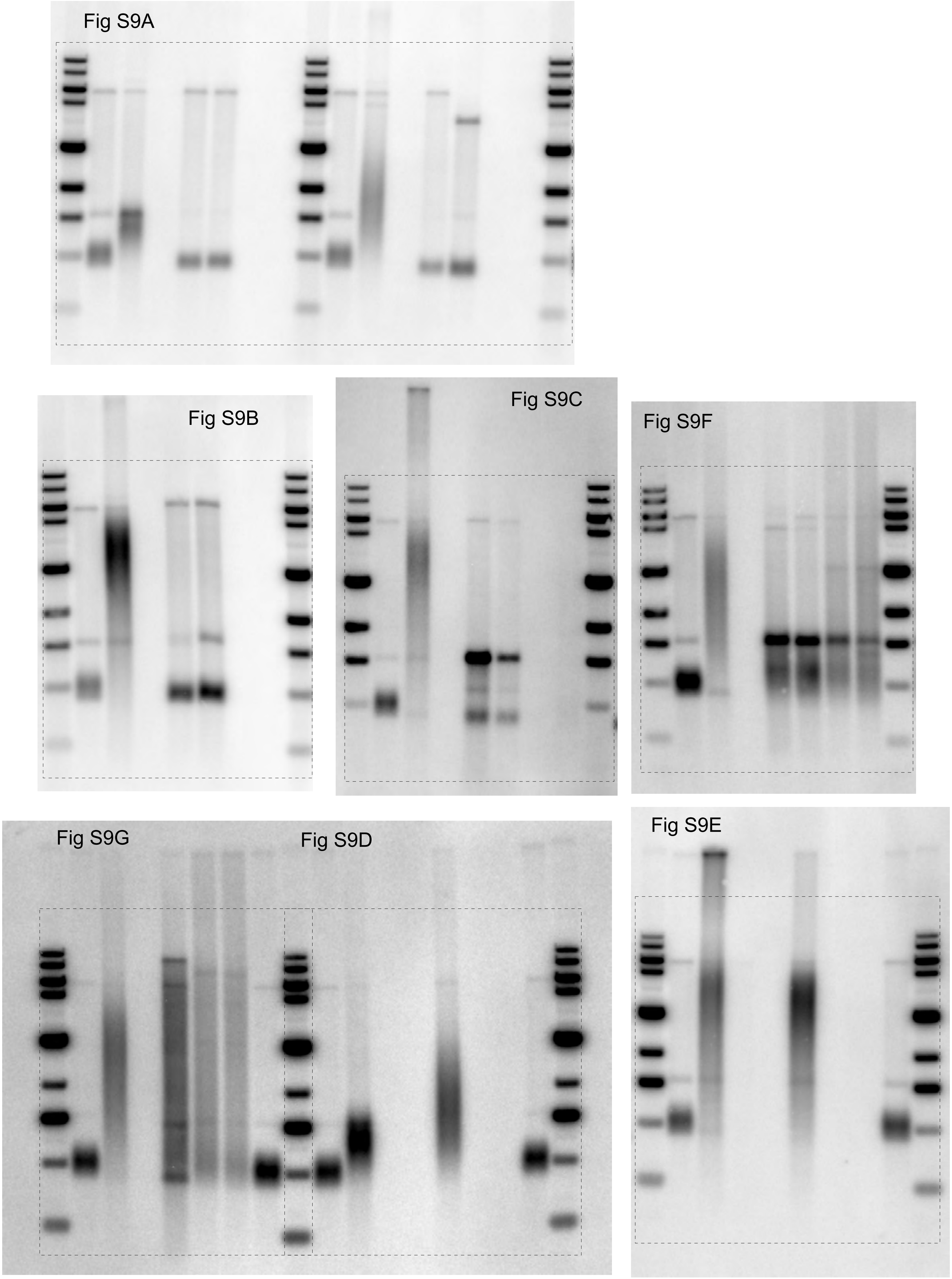
Fig S9A
Fig S9B
Fig S9C
Fig S9F
Fig S9G
Fig S9D
Fig S9E

Fig S10

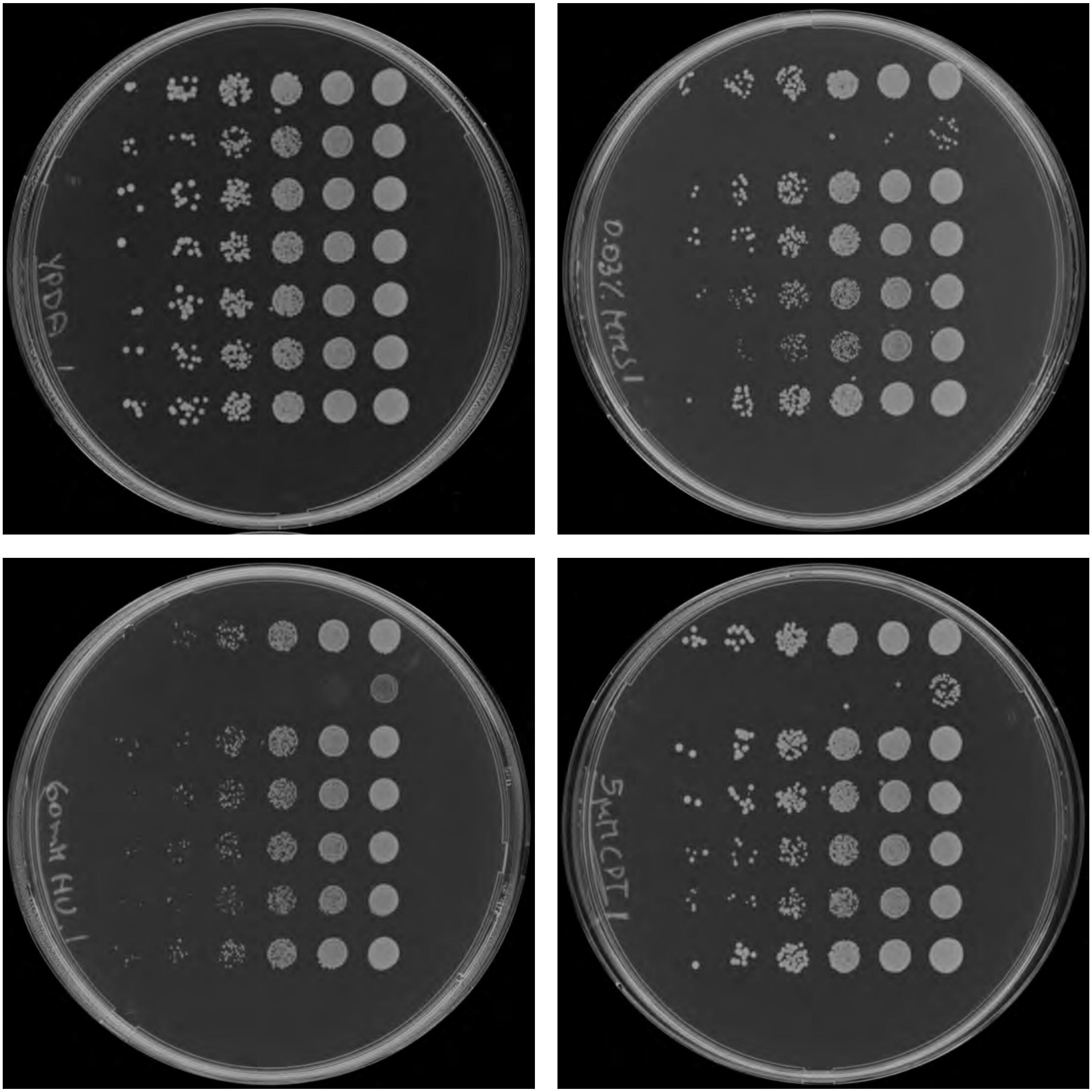
